# A reduced SNP panel optimised for non-invasive genetic assessment of a genetically impoverished conservation icon, the European bison

**DOI:** 10.1101/2023.04.01.535110

**Authors:** Gerrit Wehrenberg, Małgorzata Tokarska, Berardino Cocchiararo, Carsten Nowak

**Affiliations:** Centre for Wildlife Genetics, Senckenberg Research Institute and Natural History Museum Frankfurt, Clamecystraße 12, 63571 Gelnhausen, Germany; Department of Ecology and Evolution, Johann Wolfgang Goethe-University, Biologicum, Max-von-Laue-Straße 13, 60438 Frankfurt am Main, Germany; LOEWE Centre for Translational Biodiversity Genomics (LOEWE-TBG), Senckenberganlage 25, 60325 Frankfurt am Main, Germany; Mammal Research Institute PAS, Waszkiewicza 1, 17-230 Białowieża, Poland

## Abstract

The European bison was saved from the brink of extinction due to considerable conservation efforts since the early 20^th^ century. The current global population of > 9,500 individuals is the result of successful *ex situ* breeding based on a stock of only 12 founders, resulting in an extremely low level of genetic variability. Due to the low allelic diversity, traditional molecular tools, such as microsatellites, fail to provide sufficient resolution for accurate genetic assessments in European bison, let alone from non-invasive samples. Here, we present a SNP panel for accurate high-resolution genotyping of European bison, which is suitable for a wide variety of sample types. The panel accommodates 96 markers allowing for individual and parental assignment, sex determination, breeding line discrimination, and cross-species detection. Two applications were shown to be utilisable in further *Bos* species with potential conservation significance. The new SNP panel will allow to tackle crucial tasks in European bison conservation, including the genetic monitoring of reintroduced populations, and a molecular assessment of pedigree data documented in the world’s first studbook of a threatened species.

## Introduction

The European bison or wisent (*Bos bonasus* (Syn.: *Bison bonasus*) LINNAEUS, 1758) represents a textbook example of successful *ex situ* population management and reintroduction following severe bottlenecks and extinction in the wild in 1927. *Ex situ* and *in situ* population management is based on the world’s first studbook for a threatened species (European Bison Pedigree Book; EBPB) established for conservation purposes^1^. Today’s global population size of > 9,500 is the result of this successful population management during the last almost 100 years^1–3^. Despite this success, the species is still threatened by genetic erosion due to a small gene pool resulting from a total of only 12 founders with uneven founder representations^4–6^. Besides this massive bottleneck, the population went through several other contractions in population size before and after^7, 8^ the establishment of the breeding program in 1923, with the latest happening during World War II^9^. Additional bottlenecks still happen through initial founder effects when reintroducing a limited number of animals from captivity into the wild in the framework of reintroduction programmes. While it is presently not fully understood to which degree reduced genetic diversity hampers population fitness and adaptability to changing environmental conditions, an increased susceptibility to diseases, such as *posthitis* or *balanoposthitis* is commonly suspected to be a likely consequence of low genetic diversity and high inbreeding coefficients^1, 10^.

The current *B. bonasus* population is managed separately in two breeding lines: the lowland line (LL), representing the natural subspecies *Bos bonasus bonasus* LINNAEUS, 1758, originated from seven founders. The lowland-Caucasian line (LC) was founded by 11 founders of *B. b. bonasus*, including the seven founders of LL, and a single male of *Bos bonasus caucasicus* (TURKIN & SATUNIN, 1904). The LC is factually managed as an open population, whereas gene flow from LC into LL is undesired and its prevention is considered a priority in European bison conservation management^5^.

Because of the genetic issues mentioned above it is pivotal to track genetic diversity and relatedness in both *ex situ* as well as reintroduced populations of the European bison. However, due to genetic homogeneity of the species, standard approaches of using microsatellite markers for genetic monitoring as well as individual identification are not applicable in European bison conservation management. Tokarska *et al.*^11^ showed that single-nucleotide polymorphisms (SNPs) are more suitable to assess identity and paternity compared with microsatellites. Another important issue is DNA sampling: in contrast to often impractical and undesired invasive sampling, the ability to use non-invasive samples to assess viable genetic population data from appropriate numbers of individuals could be a valuable tool for e.g. monitoring wild species or for the use in behavioural studies^12–16^.

Consequently, a comprehensive genetic assessment with a reliable molecular method accompanying the existing conservation management in the wisent is needed to enable further preservation of genetic depletion of the already low intraspecific diversity in the long-term. Here, we present a novel reduced 96 SNP panel applicable for non-invasive samples of the European bison. The new modular marker panel tackles several conservation-relevant issues: (i) individual discrimination, (ii) parental assignment, (iii) sex determination, (iv) assessment of genetic diversity within the population, (v) breeding line discrimination and (vi) cross-species detection for European bison. Molecular resolution of parental assignment and genetic diversity in the wisent measures were evaluated with genealogical studbook data. Additionally, we evaluated the applicability of the SNP panel for further Bovini (GRAY, 1821) with potential conservation relevance in basic applications.

## Results

### General assay performance and selection of the final 96 SNP marker panel

From initially tested 231 SNP markers, 111 markers failed to amplify, showed no interpretable clusters or locus polymorphism in the European bison and were thus excluded after the first round of wet laboratory tests. A final set of 96 SNPs was selected from the remaining 120 SNP markers based on best performance with non-invasively collected samples. This final 96 SNP marker panel with overlapping subsets consisted of 90 autosomal markers for individual discrimination, 63 markers for parental assignment and the assessment of genetic diversity as well as 18 markers for breeding line discrimination between LL and LC. Six candidate SNP assays in the gonosomal amelogenin (AmelY1, AmelY2, AmelY3, AmelX1, AmelX2) and the zinc finger gene (ZFXY), respectively, were validated for sex determination in European bison and other bovines. Five assays showed consistent amplification for invasive samples, whereof four were excluded in later testing phase due to failing with non-invasive samples. Though no template controls (NTCs) were amplified within the X-chromosomal cluster, the locus AmelY1 was still found to be suitable due to the distinct Y-chromosomal-associated allele cluster and was finally included in the 96 SNP panel.

Subsets for parental assignment and genetic diversity assessment were tested for Hardy-Weinberg equilibrium (HWE) and linkage disequilibrium (LD) across 58 non-first-order relatives, resulting in a selection of 63 unlinked markers in HWE. The *R*^2^-based LD calculations estimated high linkage especially for *posthitis*-associated loci of the panel (Supplementary Fig. S3).

The mean call rate for non-invasive samples was 92.4 % and the mean genotyping error (GE) was 1.9 %, with allelic dropouts (ADOs) = 1.6 %, and false alleles (FAs) = 0.3 %. AmelY1 showed a GE rate of 0.044 across non-invasive samples. The mean call rate from invasive samples was 98 %, while the mean GE rate over all marker was close to 0 (Supplementary File *SNP_marker_list_details.xlsx*).

### Modular subsets of the 96 SNP panel

#### Individual discrimination

The microsatellite panel with 11 loci used in the pilot study did not reach sufficient resolution for the probability of identity (PID) and the probability of identity among siblings (PIDsib), which is considered to be a sufficiently low threshold for most applications involving natural populations^17^. In contrast, the SNP subset of 90 polymorphic markers reached a PID ≤ 0.0001 with ≤ 10 markers and PIDsib ≤ 0.0001 with ≤ 18 markers for *B. bonasus* (Figure 1).

**Figure 1:**
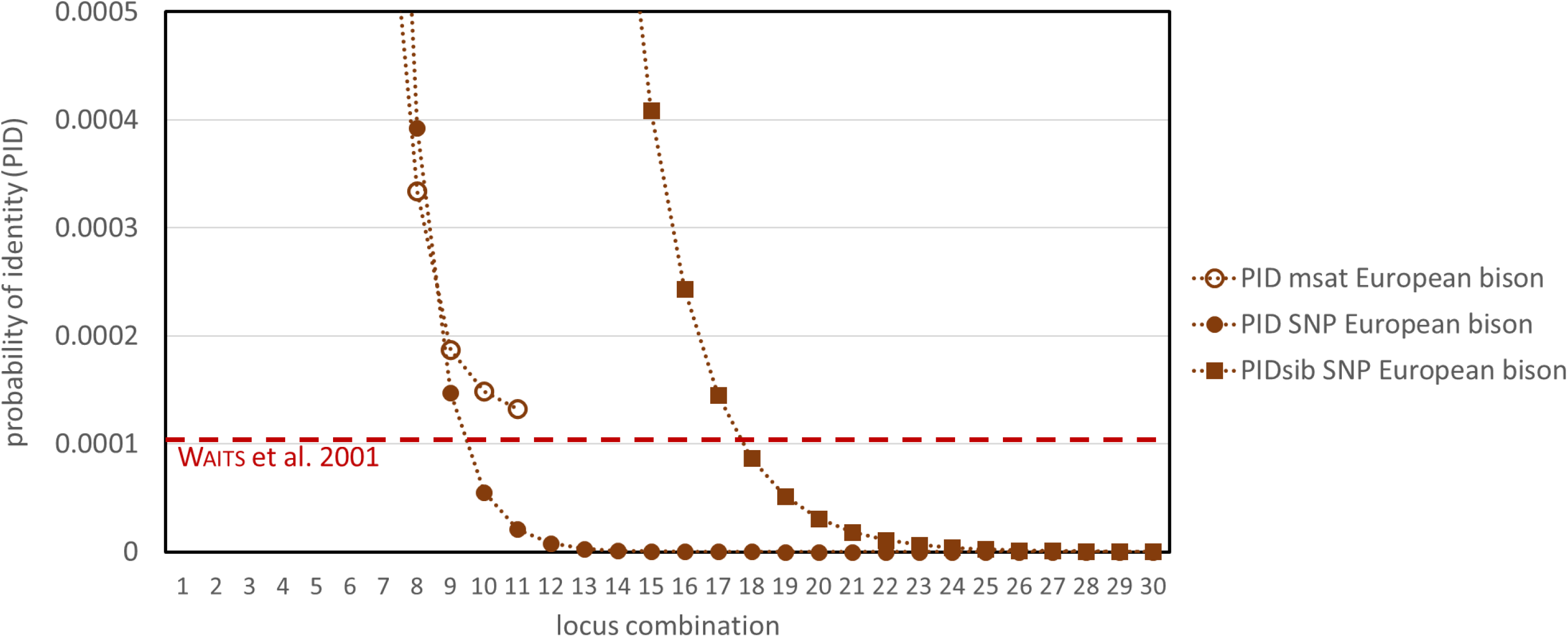
Probability of identity (PID) and probability of identity among siblings (PIDsib) of genotyped microsatellites (n = 11) and autosomal SNPs (n = 95) for European bison. Horizontal dashed red line: PID threshold for natural populations by Waits et al.^17^ is not overcome by the microsatellite panel. SNP-based PID reaches threshold at approx. 10, PIDsib at approx. 18 loci for the European bison. Approximations of PID and PIDsib close to zero are reached approx. with 13 and 24 loci, respectively. The x-axis was cut at locus combination of 30 loci for more conciseness whereby the approximation of the SNP-based PIDs does not change after 30 loci. PIDsibs estimations of the microsatellite panel are outside of the scale. PID and PIDsib for all other Bos species for which individualisation was possible based on 95 autosomal SNPs are provided in Supplementary Fig. S4.

The mean number of allele mismatches found between pairs of genotypes within the total wisent population were 28.2 (LC: 29.5; LL: 26.5), for American bison 11.2, for gaur 6, for banteng 4.1 and highest for domestic cattle with 34.9. The lowest value for European bison (= 17) was found between two first-degree relatives. The lowest number of allele mismatches in the American bison was 7, for domestic cattle 23, for gaur and banteng 4, also all between two first-degree relatives each (*Figure 2*).

**Figure 2:**
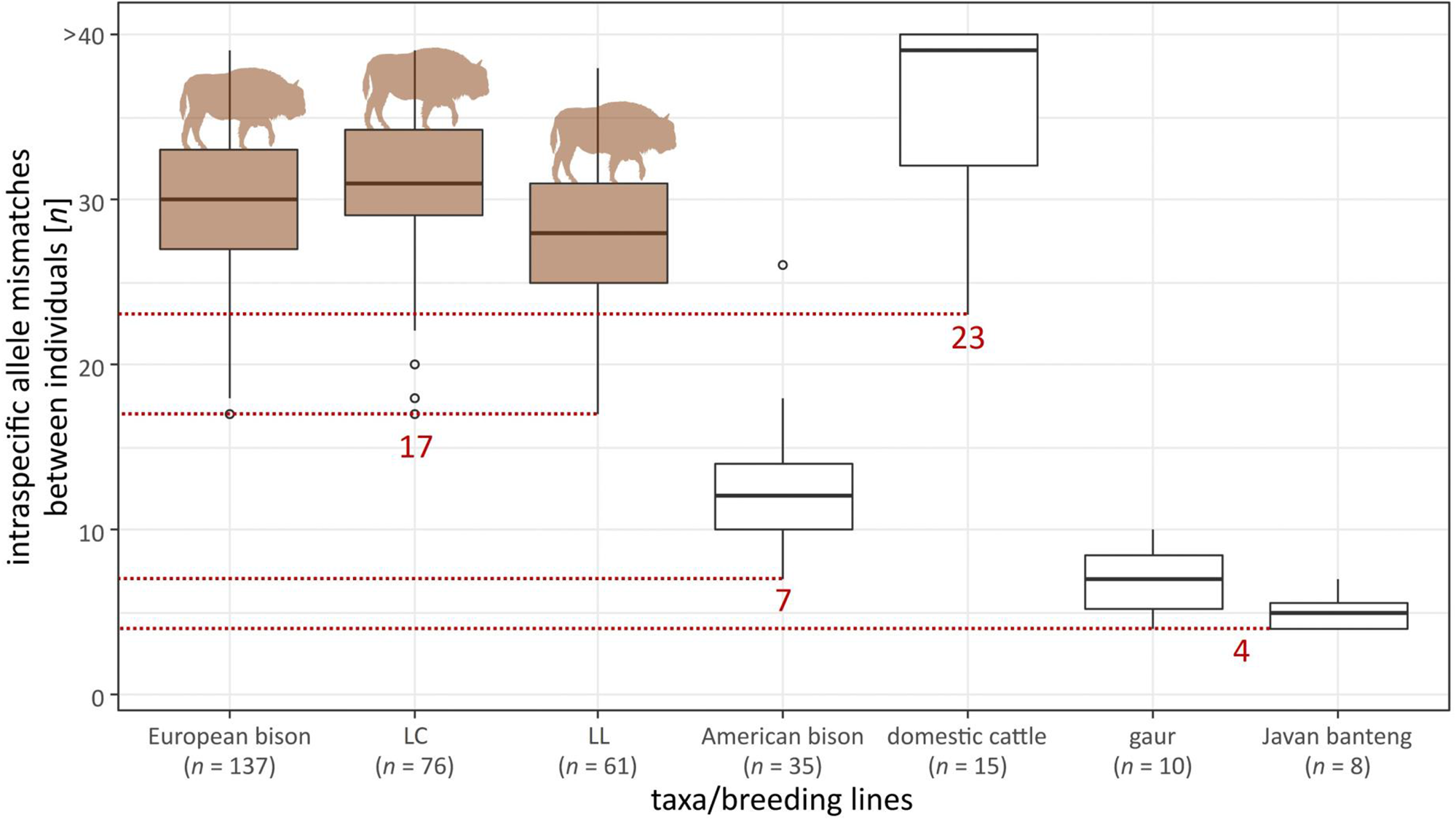
Detected number of mean allele mismatches between individual genotypes (genotypes consisting of 95 loci) of European bison (both breeding lines separately) as well as American bison, domestic cattle, gaur and banteng. Lowest allele mismatches are highlighted in red. Individual sample size per group is noted (*n*). Allele mismatches between genotypes of five unrelated cattle individuals are > 40 loci.

#### Sex determination

Correct sex determination failed for six European bison cows out of a total of 137 individuals (4.4 %). These six individual samples showed three to four FAs in the Y-chromosomal cluster within six replicates. Sex determination was also possible with American bison, yak, domestic cattle, gaur, banteng, water buffalo, lowland anoa, mountain anoa, Cape buffalo and forest buffalo. Over all 11 species (235 individuals) 92.9 % were correctly determined, 4.4 % were false positive and 2.8 % not determinable.

#### Parental assignment

Parental assignment for comparison with the pedigree data was conducted for 137 individual genotypes (see exemplary family network with 23 relatives in *Figure 3*). Of those, 128 were individually assigned during sampling in the field, while nine individual genotypes originate from not individually assigned samples. According to the studbook, 48 parental assignments were expected to be detected between the available genotypes. From these, 41 maternal and paternal relationships were correctly identified. In eight cases, the parent-offspring (PO) relationship was detected but the offspring was assumed to be the parent or vice versa. In all of those latter cases the genotype of the second parent was unknown. In seven cases the expected PO relationship was not identified. In eight cases, PO relationships were estimated false-positively compared to pedigree data. Five of these false positives were assigned to second-degree relatives, one to a third-degree and one to a fourth-degree relative with recent inbreeding involved. Despite one case of a second-degree relative all false-positive parental assignments between individuals were obtained if no true parental genotypes were available in the molecular sample set. No false-positive parental assignments between individuals of the two breeding lines were estimated.

**Figure 3:**
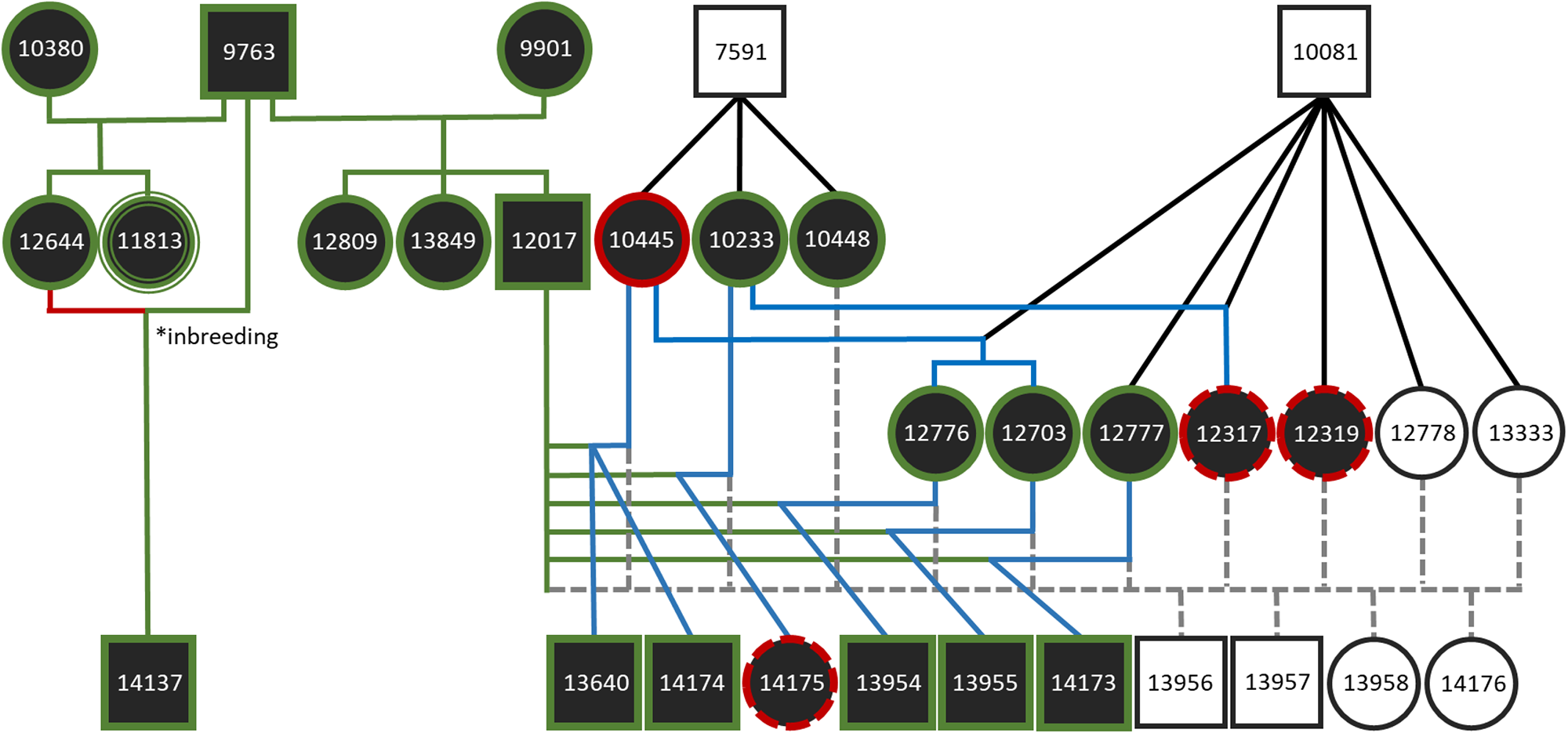
An exemplary family network to document the integration of molecular kinship analysis into the present pedigree data from the European Bison Pedigree Book (EBPB). Three generations of 23 individuals assigned to LL were sampled and genotyped from three holders in the Netherlands and Germany (Lelystad (Natuurpark), Duisburg (Zoo), and Springe (Wisentgehege)). Circles represent female individuals and squares male individuals (filled symbols: genotyped). Green edges around the individuals represent successful molecular sex verification, whereas solid red edges represent falsely positive sex assignments and dashed red edges, where no molecular sex assignment was possible. All genotypes are based on a single sample per individual. Triple edges: sample was not individually assignable in the field but was assigned to an individual with the genotype based on sex determination and parental assignment. Different colours of the genealogical lineages represent different verification states: green: genetically verified kinships from the EBPB; blue: genetically assigned kinships with lacking data in the EBPB; red: kinship from the EBPB not genetically verified; black: kinships genetically not verifiable due to missing genotypes. 10 parental assignments (sired by ‘EBPB#7591’ and ‘EBPB#10081’) with unknown maternities from the EBPB were included to visualise the high degree of at least half-sibling relationships of the females/potential mothers in Lelystad; grey dashed: presumed kinships not verifiable due to missing genotypes and missing data in the EBPB. Asterisk: case of inbreeding. All breeding line assignments of the displayed individuals were genetically verified (not noted here).

Two out of twelve originally individually unassignable field samples were assigned to known individuals documented in the EBPB through their as well genotyped parents: ‘Durana’ (EBPB#11813) and ‘Odila’ (EBPB#13951).

#### Genetic diversity

All 63 non-linked markers in HWE (Supplementary File *SNP_marker_list_details.xlsx*) were used for the assessment of genetic diversity in the European bison in comparison to pedigree-derived values. Generally, gene diversity (GD) and heterozygosity values (*H*_S_/u*H*_E_) were stable within but not consistent between molecular and pedigree data, whereas the *F*-statistics showed comparable values between both data sets (Table *1*). The *F*-statistics tend to be variable even based on same molecular or pedigree data depending on the utilised software and its calculation method. Notably lower genotype samples sizes negatively affected mostly *heterozgosities* and *F*-statistics and caused erroneous calculations most prominently in the *F_IS_* (Table 1). If calculated per breeding line, LC showed a consistently higher genetic diversity than LL (Supplementary Table S4).

**Table 1:** Genetic diversity measures based on SNP genotypes and pedigree data for different sets of European bison individuals. SNP genotype values are based on unlinked 63 SNPs in HWE. All 277 of 338 sampled individuals with known genealogy were used to generate pedigree-based genetic values. As genealogical information was not available for all successfully genotyped individuals, molecular and pedigree-based genetic diversity values were calculated for an overlapping set of 99 successfully SNP-genotyped individuals with available genealogical data. Sample sizes [n] in squared brackets show the number of individuals included in the associated pedigree up to the founders. Values in parentheses next to the genetic values represent the associated standard errors (SE). F-statistics were calculated using either arithmetic averages^1^ or based on the average HS and HT over loci^2^. Pedigree-based genetic diversity values in PMx were calculated based on kinship matrix^3^ or gene drop^4^. A more detailed table including genetic diversity values of each both breeding lines is provided in the Supplementary Table S4.

| set of individuals | *n* | SNP genotypes |  |  |  |  |  |  | pedigree |  |  |  |
| --- | --- | --- | --- | --- | --- | --- | --- | --- | --- | --- | --- | --- |
|  |  | Allelic richness | *H<sub>O</sub>*<sup>FSTAT</sup> | *H<sub>S</sub>*<sup>FSTAT</sup> | *H<sub>T</sub>*<sup>FSTAT</sup> | *F<sub>IT</sub>*<sup>GenAlEx</sup> | *F<sub>IS</sub>*<sup>GenAlEx1</sup><br>*F<sub>IS</sub>*<sup>GenAlEx2</sup><br>*F<sub>IS</sub>*<sup>FSTAT</sup> | *F<sub>ST</sub>*<sup>GenAlEx1</sup><br>*F<sub>ST</sub>*<sup>GenAlEx2</sup><br>*F<sub>ST</sub>*<sup>FSTAT</sup> | GD<sup>PMx3</sup><br>GD<sup>PMx4</sup> | *F<sub>IT</sub>*<sup>ENDOG</sup> | *F<sub>IS</sub>*<sup>ENDOG</sup> | *F<sub>ST</sub>*<sup>PMx</sup><br>*F<sub>ST</sub>*<sup>ENDOG</sup> |
| all sampled with pedigree (total) | 227 [1,296] | - | - | - | - | - | - | - | 0.825<br>0.825 | 0.059 | 0.022 | 0.024<br>0.038 |
| all genotyped | 137 | 126 | 0.400 (0.015) | 0.409 (0.014) | 0.422 (0.014) | 0.049 (0.012) | 0.017 (0.011)<br>0.015 (0.011)<br>0.024 (0.010) | 0.034 (0.005)<br>0.033 (0.005)<br>0.030 (0.005) | - | - | - | - |
| all genotyped with pedigree | 99 [982] | 126 | 0.400 (0.015) | 0.401 (0.014) | 0.417 (0.014) | 0.036 (0.015) | -0.006 (0.013)<br>-0.008 (0.013)<br>0.004 (0.013) | 0.043 (0.006)<br>0.043 (0.006)<br>0.037 (0.006) | 0.803<br>0.804 | 0.057 | 0.011 | 0.055<br>0.047 |

#### Breeding line discrimination

A subset of 18 SNP markers provided the lowest false-positive rate in breeding line assignments. This marker subset with the highest resolution was identified when the *F*_ST_ threshold per locus was set to a minimum of 0.075. It includes two out of six loci with private alleles found in LC among 137 individuals in this study (Supplementary File *SNP_marker_list_details.xlsx*).

Seven individuals (5.1 %) with the Bayesian genetic clustering (*STRUCTURE*) and five individuals (3.6 %) with the maximum likelihood genetic clustering (*adagenet*) were false-positively assigned to a breeding line (Bayesian: total: *n* = 5, LC: *n* = 4, LL: *n* = 1; Maximum Likelihood: total: *n* = 4, LC: *n* = 4, LL: *n* = 0) or were not clearly assignable (Bayesian: total: *n* = 2, LC: *n* = 1, LL: *n* = 1; Maximum Likelihood: total: *n* = 1, LC: *n* = 0, LL: *n* = 1; Figure 4). Four samples from Russia were constantly false-positively assigned to LL based on the given breeding line assignment.

**Figure 4:**
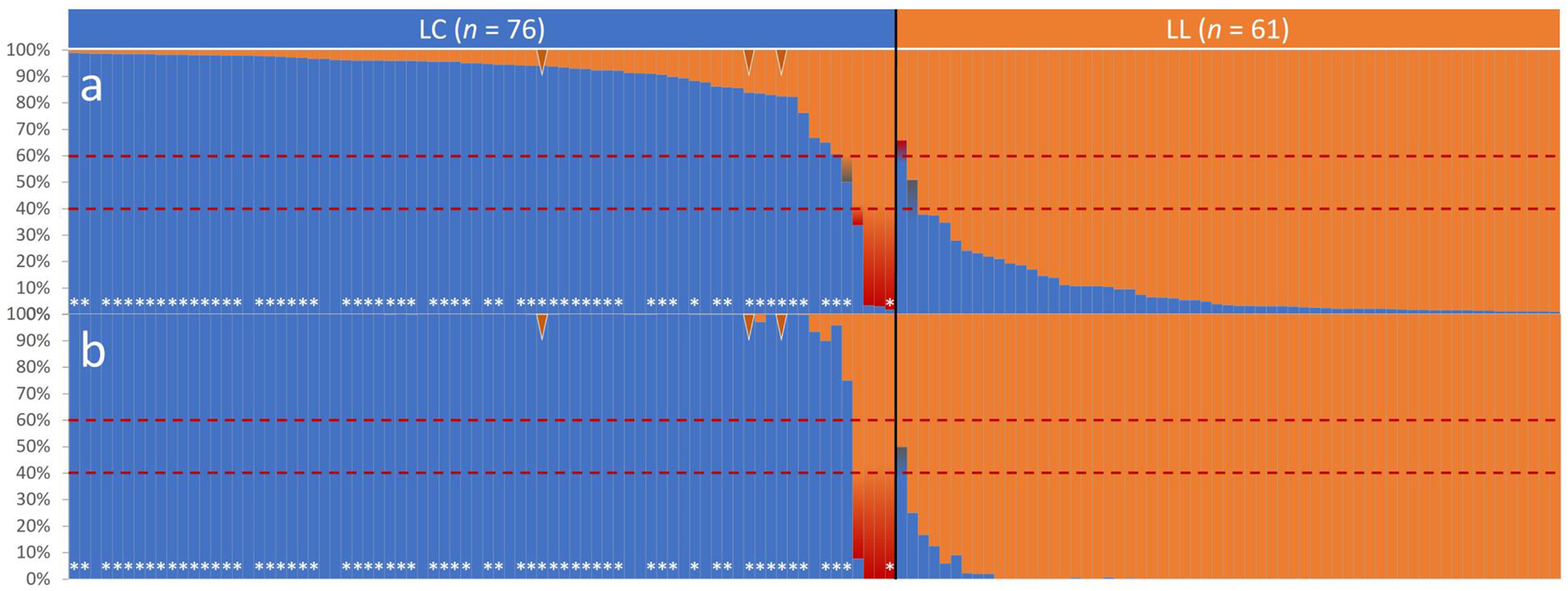
Assignment probabilities [%] based on 18 loci selected for breeding line discrimination between LC (*n* = 76) and LL (*n* = 61) in the European bison: (a) Bayesian genetic clustering computed with *STRUCTURE*; (b) Maximum-likelihood genetic clustering computed with *adegenet*. The black line shows the previously assigned lineage distinction (LC: blue; LL: orange). Dashed red lines indicate assignment thresholds. Bars tarnished red mark individuals with unexpected lineage assignment; bars tarnished grey mark individuals not assignable with genotypic data according to the assignment threshold. Brown arrows: F1 breeding line hybrids. White asterisks: LC individuals with at least one of the six private alleles found in LC. See Supplementary Table S5 for the order of individuals shown here.

#### cross-species detection

All non-target taxa with SNP call rates > 80 % (16 evolutionarily significant units (ESUs) in 10 Bovini species; *Figure 5*) could be distinguished from *B. bonasus* in a Principal Coordinates Analysis (PCoA) based on 95 or 31 (for domestic cattle) loci (Figure 6). Samples from more distantly related taxa showed generally much lower call rates and less SNP polymorphism (*Figure 5*). See Supplementary File *SNP_marker_list_details.xlsx* for SNP subsets suited for cross-species identification between several other ESUs within Bovini along with provided reference genotypes from a broad phylogenetic diversity of this tribe (Supplementary File *Genotype_lists.xlsx*).

**Figure 5:**
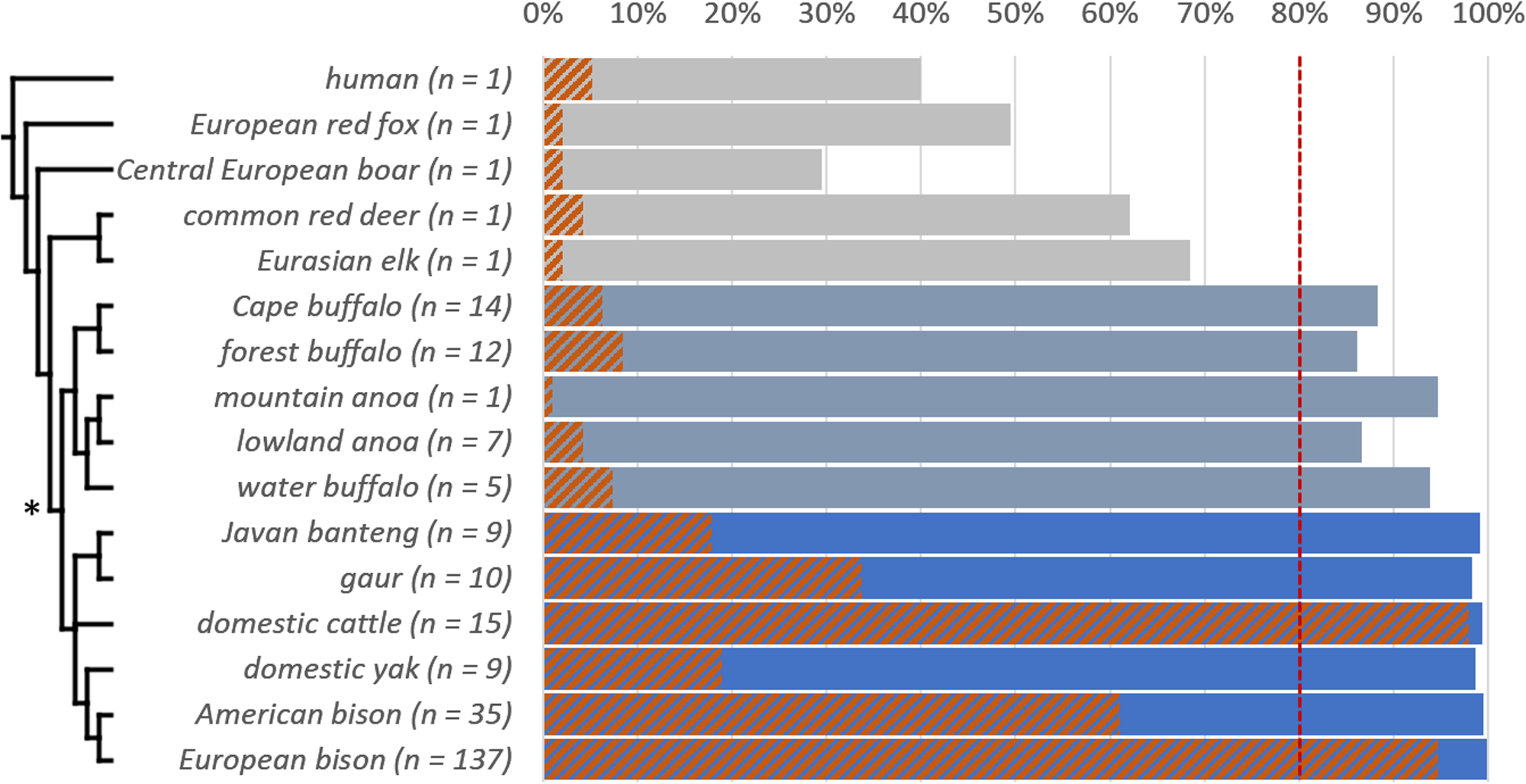
SNP call rate [%] for 95 autosomal SNPs in the European bison and 15 non-target species with corresponding numbers of individuals (*n*). The length of a solid bar indicates the mean SNP call rate for each analysed species. Blue bars reflect all groups classified to the genus *Bos*, blue-grey bars groups classified to the subtribe Bubalina and grey bars species outside of Bovini. A SNP call rate of at least 80 % call rate (red dashed line) is the threshold for inclusion into further analysis. The orange-hatched bars show the percentage of found polymorphism over 95 loci within the groups. The cladogram reflects known evolutionary relationships between the species^18^. The asterisk points out the tribe of Bovini.

**Figure 6:**
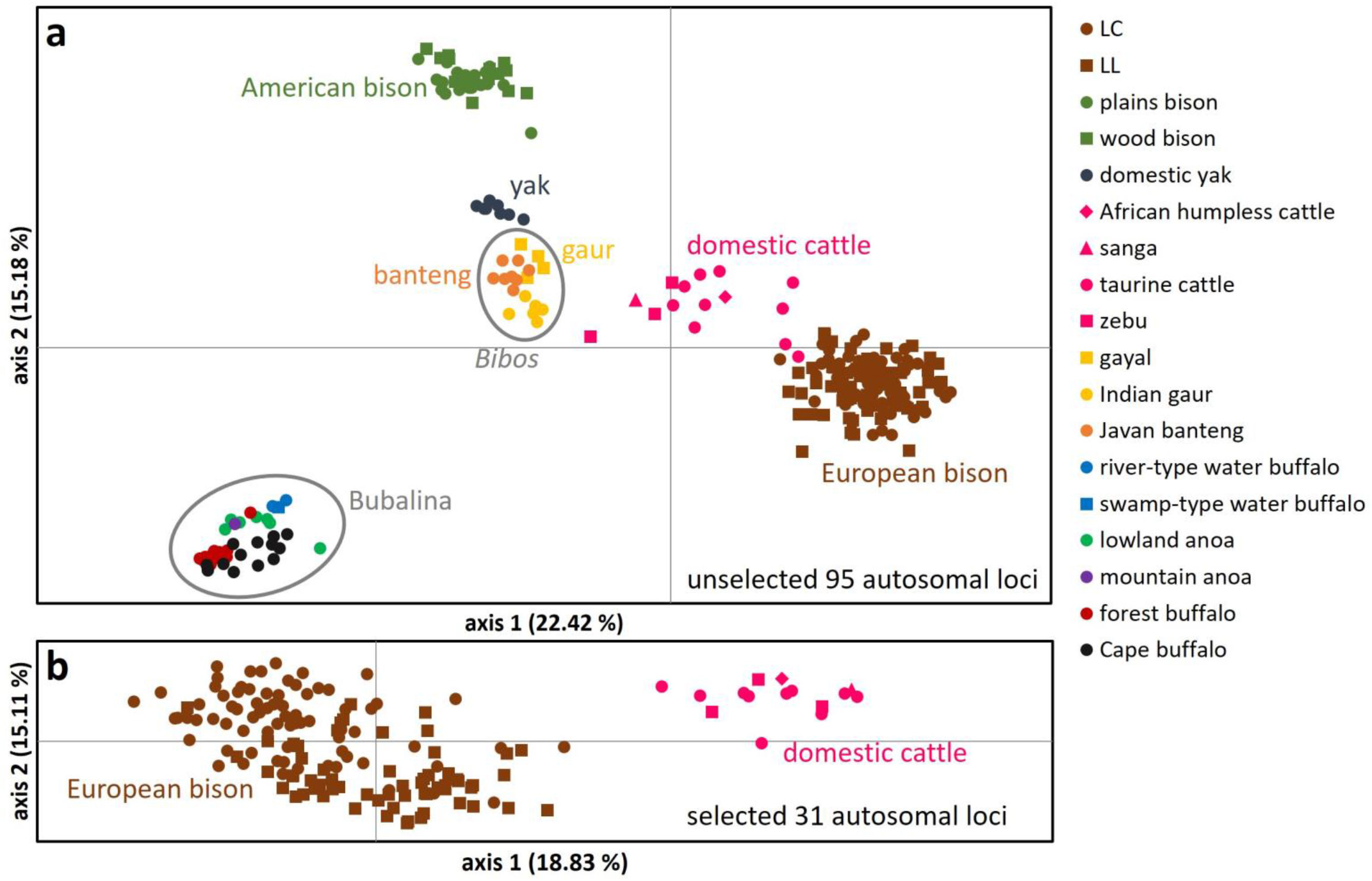
(a) PCoA of 137 European bison (both breeding lines) and 116 individuals of 10 non-target Bovini species (16 ESUs) with a SNP call rate over 80 % utilising all 95 autosomal SNP loci. (b) PCoA of 137 European bison and 15 domestic cattle (four major lineages) utilising a subset of selected 31 SNP loci. Clusters containing higher taxa like the subgenus *Bibos* (HODGSON, 1837) and the subtribe Bubalina (RÜTIMEYER, 1865) are marked in grey circles. Eigenvalues (a): axis 1: 233.68; axis 2: 158.19; Eigenvalues (b): axis 1: 33.46; axis 2: 26.85

## Discussion

### Resolution of the new SNP panel

The genetic assessment of wildlife populations via non-invasive samples reduces undesired anthropogenic interference as much as possible and consequently became common practice in wildlife genetic studies^16^. Once developed, such reduced marker panels for genotyping of non-invasive samples with genome-wide SNPs provide a standardised, fast-applicable, and low-cost genomic approach for conservation^19^. Previously, it has been shown that genotype recovery for non-invasive samples is overall higher using microfluid SNP panels compared with frequently utilised microsatellites^20^. In line with von Thaden *et al.*^20^ we also found high informative content, reliability and reproducibility of genotypes of our microfluidic SNP panel, with a high average genotyping quality across samples (average call rate = 92.4 %, GE rate = 1.9 %).

To gain for maximum resolution of the panel, we decided to accept increased amplification rates in NTCs for some selected loci. Occasional fluorescence of NTCs are known in SNP genotyping and is considered to be no major concern due to marker-specificity and inconsistency in genotype yields from NTCs^19^. With the marker GTA0242130 all NTCs showed fluorescence and solely clustered with the homozygous YY cluster. Nevertheless, this marker was kept because of the overall good clustering. If the downstream analysis was not negatively impacted lower call rates were also tolerated: a single autosomal marker (GTA0250956) showed a drastically lower call rate of 76.2 %. Since this marker is highly informative for breeding line discrimination (*F*_ST_ = 0.112 in a set of 58 individuals not in a first-degree relationship) with GE rate of 2.2 %, it was kept. Invasive samples generally showed complete call rates and minor GE rates and thus no need to be replicated with the current SNP panel.

The European bison is the only recent wild cattle in its current distribution^21^. However, within all native regions of the European bison, domestic cattle and partly domestic water buffalos occur as livestock^22, 23^ and their faeces could thus be confused during field sampling (see Supplementary Discussion for a more extensive discussion on bovid dung as a considerable genetic sample type). Therefore, it is important that obtained genotypes can be reliably assigned to the correct species to avoid biased results in a genetic monitoring. With the SNP panel presented here all genotyped Bovini could be distinguished from the European bison and furthermore, clustered according to their ESU. The proximity of the cattle cluster to the European bison cluster can be attributed to the fact that all autosomal SNPs in this study were originally detected in *B. primigenius* (Figure 6 (a)). This also causes the strikingly high degree of SNP polymorphism in this species (Figure 5). With a subset of 31 selected SNPs from the novel marker panel it is possible to genetically distinguish *B. primigenius* from *B. bonasus* (Figure 6 (b); Supplementary File *SNP_marker_list_details.xlsx*).

The new SNP panel allowed for safe individual discrimination, with considerable allele differences between most individuals. The lowest number of allele mismatches between individuals of European bison was 17 loci between first-degree relatives. This is two to three times higher than allele mismatch thresholds allowing individual discrimination known from similar SNP panels for other species^24, 25^, resulting in a high degree of confidence. This is roughly consistent if considering the commonly used probability threshold for natural populations by Waits *et al.*^17^: approx. 18 SNPs would be sufficient for reliable individual discrimination (Figure 1).

The GE rate of 0.04 for the sex marker led to six failed individual sex determinations (three false positives and three not determinable) out of a total of 137 European bison. Despite occasional misidentifications, which typically occur in genetic information derived from non-invasively collected samples^26^, this marker set will be helpful in assessing sex ratios and sex-related behaviour in free roaming European bison populations.

Reliable individual genotypes can be used for parentage analysis, which is highly susceptible towards genotyping errors^27^. Previous studies conclude that 50 – 60 SNPs selected for high heterozygosity would be sufficient to resolve paternity in the European bison^11, 28^. The number of required loci depends on the breeding line and the grade of information regarding the parents^11, 28, 29^. A 100 SNP panel has been published for parental assignment for LL exclusively^29^, of which a portion of markers were included in the current panel. Wojciechowska *et al.*^3^ developed a subset of 50 SNPs for parental assignment applied for both breeding lines. With the reduced 63 SNP subset in our study, parental assignment was also successful for LC and additionally proved effective for non-invasively collected samples. In difficult cases as shown in the exemplary family network (Figure 3), where recent inbreeding meets low genetic diversity in LL, which is expected to require more loci to resolve PO relationships^28^ the panel resolution reaches its limits and partially fails to disentangle kinship. Such cases show that only the combination of genetic assessment with available studbook and other metadata will allow to resolve patterns of relatedness with high certainty^30, 31^. This combined approach is state of the art in other comprehensive genetic population monitoring assessments^32^ and in line with the conclusion of other studies that parental assignment is strongly facilitated in case of one known parent^11, 28, 29^.

Despite the high genetic similarity due to recent origin from an overlapping subset of founders and ongoing one-directional gene flow from LL to LC^5^, the presented SNP panel allowed for reliable discrimination of the two breeding lines as an overarching requirement for conservation actions^5^. While breeding line discrimination has previously been achieved with sets of 1,536^28^ and 30 selected SNPs^3^, our subset of just 18 markers achieved a comparable resolution including F_1_ breeding line hybrids, which are formally assigned to LC following the official management definition^5^ (Figure 4).

Among the tested samples only four individuals from ‘Russia’ documented as LC individuals clearly clustered in LL regardless of the utilised clustering method. While wild herds founded only by LL individuals in Russia are known^33^, we have no detailed information regarding those particular samples, and thus the reason for the apparent incongruency cannot be deduced here.

The finding of six private alleles within LC is not surprising since this breeding line carries genetic material of five additional founders including one bull from a separate subspecies^5^. The absence of any of six private alleles in 16 LC individuals (Figure 4) shows the low information content just relying on those markers and the need for a more discriminative markers if aiming for a robust breeding line separation as the one presented here. The discriminative value for SNP alleles published by Kamiński *et al.*^34^, which were described to be private for one breeding line, could not be confirmed in our study. This can be explained by the small and presumably not representative sample size of only ten individuals genotyped in the aforementioned study.

Neither the private alleles nor the other discriminative markers have assignable genetic origins from one of the two subspecies, *B. b. bonasus* or *B. b. caucasicus* and could be a consequence of distinct breeding management during past decades. The marker subset for breeding line discrimination presented here is thus not suitable for a validation of both the breeding lines as ESUs. Solely designed to assign individuals to the currently predefined anthropogenic breeding lines it cannot be applied to argue for or against the separate management of the two breeding lines within the European bison.

### Comparing genetic diversity estimates between studbook and molecular data

Given its history of consecutive bottlenecks and genetic depletion, an appropriate genetic marker system for *B. bonasus* should, besides individual discrimination and parentage analysis, allow for measures of genetic diversity in order to aid population management^35^. For this we selected 63 autosomal unlinked loci in HWE found to be polymorphic in the European bison. The SNPs utilised in this panel were originally detected in domestic cattle^3, 11, 28, 29, 34, 36–39^. Though common practice^40–42^, it is obvious that such a reduced number of SNPs found in a related species as well as an ascertainment bias from selecting for high polymorphism in our target species will not allow for unbiased estimates of genetic diversity^43, 44^. Thus, any results regarding genetic diversity using this SNP panel need to be interpreted with caution.

Different aspects were considered to reduce an ascertainment bias in the current SNP panel. Studies assessing genetic diversity often face the problem of incomplete population sampling^35^. In this study, the pedigree-based founder representation of the genotype set (*n* = 99) was compared with a larger pedigree data set of in total 1,296 individuals including all genotyped individuals up to all known founders to validate its population representativity beforehand (Table 1; Supplementary Fig. S1). An overall ascertainment bias can be reduced when ancestral populations are used to develop SNP panels applied on derived populations^45^. Until today, reintroductions of European bison are largely sourced from the captive population, which therefore resembles an ancestral population from which the majority of individuals for the SNP selection process originated. Overall, our genotyped individuals represent approx. 1.5 % of the current generally highly admixed^46^ global ancestral population (status 2020).

Not surprisingly, estimates of relatedness or inbreeding based on sufficient pedigree data are generally more accurate than marker-based estimates^47^. However, often no pedigree data is available for conservation-related population studies. Even for the otherwise well documented European bison, this is the case for reintroduced free-roaming herds. Additionally, pedigree-based estimations may suffer from underestimated inbreeding in the founder population^48^ as well as uncertainties towards the correctness of parental assignments, which can result in an accumulation of errors over time. This concern has been raised as well for the EBPB^49, 50^. It is also known that genetic diversity estimates, whether based on pedigree or molecular data, suffer from small sample sets especially with small gene pools caused by inbred populations and/or sample sets with high portions of closely related individuals^51, 52^. Thus, estimation accuracy will be increased by larger sample sizes and decreasing sampling variance of reference genotypes^47^, particularly within the breeding lines.

Since the 63 SNPs utilised for genetic diversity estimations were specifically selected for high polymorphism, it is obviously not appropriate to directly compare pedigree-based GD values with molecular-based heterozygosities. Still, it is interesting to note that SNP-based fixation indices resemble the pedigree-based values (Table 1). The relatively low *F*_IS_ is caused by high intermixture within the breeding lines, whereas rare gene flow between LC and LL is manifested in the second highest fixation estimated in the *F*_ST_. Overall, we found a high degree of admixture over the population, despite of the species’ strongly reduced gene pool. This finding, which is consistent with a recent study utilising 22,602 SNPs^46^ is a consequence of the successful population management during the last decades. The highest fixation seen in the *F*_IT_ is caused by different allele frequencies within the breeding lines compared to the total population and is known as the Wahlund effect^53^. Changes in fixation indices among populations can be caused by dynamic processes such as genetic drift, gene flow, migration or bottleneck events^54^. Since one of the biggest threats for the European bison is genetic erosion, the new SNP panel can be used to effectively track such trends and changes in genetic diversity and aid conservation efforts aiming at the establishment of stable populations in the wild. Thus, long-term monitoring of genetic diversity will also enable an evaluation of laborious and costly reintroduction efforts for decision makers.

### Potential application on other Bovini species

The IUCN red list contains 12 Bovini species (*Syncerus* spp. included in this study are recognised as conspecific) of which 9 species are listed as threatened (VU: *n* = 2; EN: *n* = 4; CR: *n* = 3)^55–66^. A genetic assessment of those wild cattle, similar to the European bison is therefore of considerable interest. The SNP marker panel presented here was solely developed for *B. bonasus*. However, as all autosomal SNPs were originally discovered in *B. primigenius* but are still polymorphic in the European bison, those to some degree evolutionary conserved orthologous SNPs may allow for utilisation in closely related species. Demonstrably, this SNP panel can be utilised for sex determination in all Bovini species (success rate of 92.9 %) as well as for individualisation in American bison (both subspecies), domestic cattle (with all four major lineages), gaur (including gayal) and banteng from non-invasive samples. Thus, the new SNP panel developed for the European bison has instant potential for basic population genetics or conservation applications in other threatened wild cattle and may serve as basis for further optimised panels.

### Implementation in conservation and research of European bison

The SNP panel presented here has been specifically developed for current questions and needs in *ex* and *in situ* conservation of the European bison. Free-roaming European bison are not listed individually in the EBPB and therefore lack genealogical documentation^2^. The new SNP panel provided here allows the assessment of relationships between wild individuals without the need to catch or harm the animals, and allows for continuous, systematic genetic monitoring, which is recommended to improve *in situ* conservation efforts^67^. Genetic population monitoring generates important information for decision makers and can also help raise public awareness^68, 69^. The panel may be as well used to generate sound data in human-wildlife conflicts, which may arise due to damages in forestry or agriculture^70^. To allow for an effective long-term monitoring of wild populations, it is strongly recommended to genotype all reintroduced founder individuals. Complementing this approach with a subsequent continuous non-invasive genetic monitoring will allow to track population developments over time and help disentangle the effects of e.g. genetic drift, population isolation, migration, and/or changes in (effective) populations sizes, home ranges and social structure following reintroductions^71^.

Even more than 50 years since the first reintroductions, the captive wisent population is still the source for current rewilding efforts. Therefore, an assessment of the *ex situ* population must go hand in hand with the *in situ* conservation actions re-establishing Europe’s last species of wild bovines. *Ex situ* breeding strategies based on pedigrees are tested to be efficient if sufficient genealogical data is available for a species^48, 72^. Until today, this pedigree data is utilised for breeding, culling and reintroduction recommendations^5, 73^. However, due to the above-mentioned weaknesses of pedigree-based estimations on genetic diversity an independent assessment is needed. Further unintended documentation errors in the EBPB are still possible due to certain husbandry conditions, unknown paternity in herds with several mature bulls or natural behaviours like alloparental care, especially non-maternal suckling, known in European bison^1^. Formally unknown maternal relationships, genetically identified with the new SNP panel presented here, already have found their way into the EBPB. SNP-based marker-assisted breeding strategies in addition to the traditional practice based on the EBPB have been recommended before^36, 74^. This might be especially true for populations with high inbreeding, where it is presumably more important to practice population management based on genetic diversity instead of management purely based on heredity.

Besides its obvious application in population monitoring, the SNP panel may as well serve in research projects aiming at studying various aspects of conservation-relevant European bison biology, e.g. to investigate the influence of dominant male mating behaviour on the genetic structure and effective population size of the species. Furthermore, due to its robustness towards low quality samples, the analysis of collection specimens^75^ and historical hunting trophies^76, 77^ could provide interesting insights into the development of genetic diversity over time. Recently, the focus on *posthitis*-associated SNPs^39, 78^ paves the way for an utilisation of genetic assessments of this disease important for wisent conservation management. In prospect, twelve *posthitis*-associated markers were included into the current SNP panel (Supplementary File *SNP_marker_list_details.xlsx*). Due to the lack of presence-absence information of *posthitis* in the genotyped individuals of this study, further investigation is needed.

Despite of the moderate marker number our SNP panel provides a viable tool to monitor reintroductions, validate, revise and construct pedigrees, and assess population structures where no pedigree data is available. Thus, the new SNP panel represents an optimised compromise between the needed non-invasive sampling method, cost-efficiency needed for the application in conservation and the resulting informative accuracy, which is demonstrably and reasonably sufficient for the purpose it was developed for. While other recently presented SNP panels lack implementation in appropriate assays^3, 11, 28, 29, 34, 79^ the presented marker panel is non-invasive genotyping approach for the European bison ready to be used in conservation and monitoring studies. Ongoing real-world application comprises dung-based genetic monitoring of the reintroduced European bison in the Țarcu Mountains, Romania (LIFE RE-Bison; LIFE14 NAT/NL/000987). We propose the wider use of this panel both for *ex situ* population management as well as genetic monitoring of reintroduced European bison.

## Methods

All statistical analyses and most graphical visualisations were conducted using *R* v3.6.0^80^ within *RStudio* v1.0.43^81^.

### Pedigree data

All EBPB editions from 1947 to 2018 were reviewed to assess genealogical data and to create a total pedigree data set of all European bison sampled in this study (*n* = 337) up to the founders. The software *mPed*^82^ was used to convert the pedigree data into a readable format for *PMx* v1.5.20180429^83^.

### Sampling and sample storage

This study focused mainly on the collection of faecal samples, however, hair, urine, saliva and nasal secretion as valuable non-invasive sample types were also collected. Invasive sample types like muscle tissue were used as reference samples and originated from study-unrelated samplings. No harmful sampling was undertaken in the framework of this study. Within this study 253 individual genotypes from European bison (*n* = 137; LC: *n* = 76; LL: *n* = 61) and additional 15 species were analysed: ten Bovini species in 16 ESUs: American bison (*Bos bison* (Linnaeus, 1758): *n* = 35; plains bison (*B. b. bison* (Linnaeus, 1758)): *n* = 22; wood bison (*B. b. athabascae* (Rhoads, 1897): *n* = 13), domestic yak (*Bos mutus grunniens* (Linnaeus, 1766): *n* = 9), domestic cattle in four ESUs (*Bos primigenius* (Bojanus, 1827): *n* = 15; taurine cattle (*B. p. taurus* (Linnaeus, 1758)) in eight breeds: *n* = 10; African humpless shorthorn cattle (*n* = 1); sanga: *n* = 1; indicine cattle/zebu (*B. p. indicus* (Linnaeus, 1758)) in three breeds: *n* = 3^84, 85^), gaur (*Bos gaurus* (Smith, 1827): *n* = 10; Indian gaur (*B. g. gaurus* (Smith, 1827)): *n* = 6; gayal (*B. g. frontalis* (Lambert, 1804)): *n* = 4), Javan banteng (*Bos javanicus javanicus* D’alton, 1823: *n* = 8), water buffalo (*Bubalus arnee bubalis* (Linnaeus, 1758): *n* = 5; river-type: *n* = 4; swamp-type: *n* = 1^86, 87^), lowland anoa (*Bubalus depressicornis* (Smith, 1827): *n* = 7), mountain anoa (*Bubalus quarlesi* (Ouwens, 1910): *n* = 1), Cape buffalo (*Syncerus caffer* (Sparrman, 1779): *n* = 14) and forest buffalo (*Syncerus nanus,* (Boddaert 1785): *n* = 12). For cross-species tests five further species with each one individual were included: Eurasian elk (*Alces alces alces* (Linnaeus, 1758)), common red deer (*Cervus elaphus elaphus* Linnaeus, 1758), Central European wild boar (*Sus scrofa scrofa* Linnaeus, 1758), European red fox (*Vulpes vulpes crucigera* (Bechstein, 1789)) and human (*Homo sapiens* Linnaeus, 1758) (Supplementary File *Sample_list.xlsx*).

Captive sampling was done in 37 institutions from eight European countries. Samples from free-roaming LL individuals originate from the Białowieża and Knyszyńska forests in Poland and a single bull shot near Lebus in Germany in 2017. Samples from free-roaming LC individuals were collected in Russia and the Rothaar mountains in Germany between 1990 and 2017. Samples from non-Bovini species were taken from our internal collection of wildlife samples.

For sampling of faeces, hair, body liquids like urine, saliva, nasal secretion or blood from environmental surfaces sterile gloves and cotton swabs were used. Beside storage of faecal swab samples in InhibitEx buffer (Qiagen, Germany) all swabs and hair samples were stored in a filter paper and pressure lock bags including a silica gel sachet. Most pure urine samples were collected from urine-soaked snow in winter^88^. In order to test optimised faecal sampling for genetic analysis, several sampling and preservation methods were previously validated in a pilot study (Supplementary Information), resulting in two equally-suited approaches: (i) collection of 10 – 15 g of interior faecal matrix with a one-way forceps and storage in 33 ml of 96 % EtOH, (ii) swabbing the interior part of faeces and storage in InhibitEX buffer. For this study no tissue samples were invasively collected, unless as by-product from occasionally conducted mandatory earmarking by zoo personnel.

All samples were stored at room temperature (RT; 20 – 21 °C), except blood samples in Ethylenediaminetetraacetic acid (EDTA), which were stored at -20 °C. Beside from dead individuals some fresh blood samples independently originate from veterinarian procedures occurring alongside this study. Some beforehand stored blood samples were also provided by some holders (collected between 2014 – 2019).

### DNA extraction

DNA extraction of non-invasive or minimally invasive samples (hairs, scats, saliva swabs) was conducted in a laboratory dedicated to processing of non-invasively collected sample material^12^. The QIAamp Fast DNA Stool Mini Kit (Qiagen) for faecal samples and the QIAamp DNA Investigator Kit (Qiagen) for all other non-invasive sample types, respectively, were used to extract DNA on the QIAcube system (Qiagen) generally following manufacturer’s instructions with some adjustments (Supplementary Tables S8 – S10). DNA from invasive samples was extracted with the Blood&Tissue Kit (Qiagen) according to the manufacturer’s protocol. Nucleic acid concentrations of DNA extracts from invasive samples were measured with a Nanodrop spectrophotometer. Isolated DNA was stored at 4°C until use.

### Pilot study: faecal sampling, preservation and sample storage methodology

To account for the aforementioned methodological challenges, we tested for best practice in faecal sampling, sample preservation and DNA extraction from wisent dung. Mainly faeces, but other invasive and non-invasive sample types of the European bison were analysed with a set of 14 polymorphic out of 21 microsatellite markers from non-coding regions originally developed for different even-toed ungulate species and a sex determination marker^89^ to evaluate the applicability of the different sampling and storage methods. In the present study, 16 of these markers were applied for the first time to European bison. Using Generalised Linear Mixed Models (GLMMs), we statistically evaluated sampling, sample preservation and DNA extraction of wisent dung and used these results to extrapolate the finally used best practice (Supplementary Information).

### Selection of SNP loci and SNPtype assay design

All autosomal SNP loci tested in this study originate from the BovineSNP50 Genotyping BeadChip and BovineHD Genotyping BeadChip (Illumina). A set of 231 informative SNP loci for the European bison was selected from available publications for initial testing (Supplementary File *SNP_marker_list_details.xlsx*): 14 SNPs with the strongest association to *posthitis*^78^, 43 most polymorphic SNPs from Kamiński *et al.*^34^, respective 43 loci from Oleński *et al.*^29^ filtered by PID, additionally 44 SNP loci from unpublished data by high polymorphic information content (PIC) and 81 SNPS for breeding line discrimination using loci with highest contrary allele frequencies between LL and LC. It is noted that further promising SNP loci from the study Wojciechowska *et al.*^28^ for more accurate breeding line discrimination were not available due to missing indication of used loci. For sex determination, a SNP (ZFXY) found in the homologous zinc finger gene distinguishing between the gonosomal ZFX and ZFY with a C/T transition^90^ was included. Five gonosomal SNPs were identified in the amelogenin gene of European bison, plains bison, taurine cattle and zebu, yak, banteng and gayal using sequence information from GenBank® (www.ncbi.nlm.nih.gov/genbank; Supplementary File *SNP_marker_list_details.xlsx*). Subsequently, SNPtype assays were designed based on sequence information of approx. 300 bp for each SNP locus using the web-based D3 assay design tool (Fluidigm corp.). SNPs were rejected from the initial selection if not traceable at the European Bioinformatics Institute (EMBL-EBI; http://www.ebi.ac.uk) to avoid SNP duplicates or if primer design by Fluidigm corp. failed.

### SNP panel development and genotyping

We followed the development guidelines for genotyping degraded samples with reduced SNP panels provided in von Thaden *et al.*^25^ to obtain a final 96 SNP panel for implementation into a microfluidic chip system. The following sample set was used during the entire testing phase: 46 invasive reference samples (LL: *n* = 17; LC: *n* = 21; taurine cattle: *n* = 6; plains bison: *n* = 2) and 90 non-invasively collected samples. For initial wet laboratory tests, we used 150 *in silico* SNPtype assays in two partitioned genotyping runs to filter for markers with (i) proper amplification and (ii) high informative value. Assays showing failed amplification or indistinct clustering were excluded for final panel selection. All reference samples were normalised before genotyping towards the recommended concentration of 60 ng/µl (Fluidigm). Those samples did not undergo a STA (specific target amplification) pre-amplification step to enrich the target regions for SNP genotyping.

In the next step, serial dilutions of the reference sample set were prepared to concentrations of 5 ng/µl, 1 ng/µl and 0.2 ng/µl and genotyped with the remaining pool of SNPs after filtering to test the markers’ applicability on low template concentrations and subsequent pre-amplification.

### Specific target amplification and SNP genotyping

The SNP genotyping procedure using 96.96 Dynamic Arrays™ with integrated fluidic circuits (IFCs)^91^ was conducted according to the manufacturer’s protocol for genotyping with SNPtype^TM^ Assays (Advanced Development Protocol 34, Fluidigm corp.). Low DNA samples were pre-amplified in a modified STA for enrichment of the target loci before the SNP genotyping PCR. The pre-amplification of the target regions was conducted using 14 cycles for invasive samples and 28 cycles with extracts from non-invasive samples according to von Thaden *et al.*^25^.

All experiments and sample setups included NTCs (no template controls) and STA NTCs. In all experiments NTCs and samples were replicated.

#### Validation of SNP markers and scoring procedure

Raw data analyses of all runs were conducted with *Fluidigm SNP Genotyping Analysis* v4.1.2 software (Fluidigm) after 38 thermal cycles. Automated clustering and allele scoring of every SNP marker was manually checked and corrected if needed according to the guidelines suggested by von Thaden *et al.*^20^. During the development phase every SNP cluster was compared to its profile in former chip runs to keep uniformity in allele scoring. If the clustering pattern of SNP markers diverged to the pattern in former runs the complete marker was disregarded and scored as ‘No Call’ for all samples. Alleles appearing too far from the centre of a cluster were ranked as FAs and were also scored as ‘No Call’.

#### Validation of genotyping errors

Genotyping errors (GE) of each single replicate were calculated based on a consensus multilocus genotype (subsequently called reference genotype) which was built using all replicates of a sample (for consensus genotypes see Supplementary File *Genotype_lists.xlsx*). Accordingly, the following rules were applied: In general, the majority rule was applied across replicates. Loci equally scored as homo-and heterozygous were considered heterozygous. For all autosomal loci: if a locus was scored partly to be heterozygous and both opposite homozygous genotypes were found at least twice in other replicates, the genotype was defined as heterozygous. If every possible zygosity was shown in triplicates, the locus was considered to be heterozygous as well. If both homozygous genotypes were scored the more frequent zygosity was assigned. If both homozygosities were scored with 50 %, no zygosity was assigned in the consensus. Sex information for the tested individual was used as reference for calculation of the sex markers’ GE.

### Characteristics of the final 96 SNP panel

The 96 SNPs of the final panel are distributed throughout all *B. primigenius* chromosomes except autosome 25, which was not represented in the initially tested 231 SNPs as well (Supplementary File *SNP_marker_list_details.xlsx*). With 2*n* = 60, the European bison carries the same number of chromosomes^92^, which suggest a similar distribution of the used SNPs in both species.

Several applications of *GenAlEx* v6.5^93^ were used for evaluation and assessment of the molecular data as explicitly noted below. A test for LD of the 90 autosomal markers polymorphic in the European bison was conducted using squared allelic correlation (*R*^2^) utilising the *R* package *LDheatmap*^94^.

#### Cross-species detection

Five cross-species markers (GTA0250958, GTA0250953, GTA0250963, GTA0250909, GTA0250962) were selected to be monomorphic in the European bison and polymorphic in the most common sympatric bovine species (domestic cattle) or sister species (American bison), respectively. Those five markers were utilised for cross-species detection only.

In total, 24 taxa/ESUs were selected for the cross-species test on the basis of the following criteria: potentially sympatric with the European bison^95, 96^ and represent candidates for potential confusion in environmental traces such as faeces and stripping damage or sample contamination due to faecal wallowing. All further Bovini, representing the closest living relatives up to the tribe level collectable in Europe, were also included for cross-species detection. Human was included to test for methodological contamination. All samples with a SNP call rate over 80 % were analysed with a PCoA using all 95 autosomal loci executed in *GenAlEx*.

#### Individualisation

The discriminative power of the polymorphic autosomal SNP set (90 loci) and of the microsatellite panel (11 loci, data from pilot study) was assessed by estimating PID and PIDsib in *GenAlEx*. The loci were sorted according to the highest expected heterozygosity (*H*_E_).

The number of allele mismatches between individual genotypes were compared: the lowest number of allowed allele mismatches were expected between close relatives and were used as a guidance threshold for individual discrimination. Except for the sole mountain anoa all genotype sets per species contained first-degree relatives. Only those Bovini species were considered with an allele mismatches ≥ 1.

#### Parental assignment

The software *Colony* v2.0.6.6^97^, using the Full-likelihood analysis method was utilised to estimate Parent-Offspring (PO) relationships between all 137 individuals with a subset of 63 SNPs in HWE and without loci in LD. The Full-likelihood method was chosen because it was shown to be the most accurate method of *Colony*^98^. The estimations were computed with default assumptions except the following settings: male and female polygamy and inbreeding were assumed since both cases were present in the data set. *Very high likelihood precision* with *allele frequency updates* in a *very long* run was executed. All 137 individuals were put in as offspring and assigned to their sex with the probability of a sire or a dam in the data set = 0.5. No parental sibling inclusion or exclusion were added. It was only excluded for every individual to be its own parent. These settings were chosen to simulate a blind genetic monitoring study where only information is available from the genotypes and the sex determination marker. Genotyping error rates were assumed to be 0.0001 per locus because the used consensus genotypes were generated from at least triplicates and assumed to be reliable.

For validation, an exemplary family network of 23 individuals was chosen, whereof relationships of a bigger part were known. This showcase included three generations from different parks (different sample types from different collectors), many possible parents in siblinghoods, a case of inbreeding, individually assigned and not assigned samples as well as individuals with undocumented maternities and thus, visualise the full range of applications for parental assignment (*Figure 3*).

#### Breeding line discrimination

Based on 58 individual genotypes without first-degree relatives *GenAlEx* was used to identify markers with highest *F*_ST_ in each of the breeding lines to minimise an allele frequency bias by relatedness. If both parents were genotyped, the offspring were removed to obtain the highest allele variation possible. Two methods for genetic clustering were applied to the descriptive markers to test the robustness of the breeding line marker subset across different statistical approaches. Thus, the subsequent analysis was conducted assuming *K* = 2. A minimum breeding line discrimination threshold of 60 % probability was set for both genetic clustering methods.

##### Bayesian genetic clustering

To infer the presence of a distinct breeding line structure the systematic Bayesian clustering approach of *STRUCTURE* v2.3.4^99–101^ was used for microsatellite (Supplementary Fig. S6) and SNP genotypes (*Figure 4*) with burn-in periods of 250 000 repetitions and 500 000 MCMC (Markov Chain Monte Carlo) repeats. The simulations were set with *K* = 1 – 10 each with 10 iterations. *STRUCTURE HARVESTER*^102^ was used to select the most likely *K* value. *CLUMPP* v1.1.2 was used to combine the iterations of the most likely *K* value with the *FullSearch* algorithm among 10 *K*^103^.

##### Maximum-likelihood genetic clustering

The function *snapclust*^104^ implemented in the *R* package *adagenet* v2.1.1^105, 106^ was used to infer the presence of distinct genetic structures between the two breeding lines. The Bayesian information criterion (BIC) among *K* = 1 – 10 was used to estimate the most likely *K* value (Supplementary Fig. S5).

#### Assessment of molecular genetic diversity

To select a marker subset for the assessment of genetic diversity in the European bison all markers deviating from HWE within 58 non-first-degree-relatives were discarded utilising *χ*^2^ test in *GenAlEx* and *Arlequin* visualised in ternary plots (Supplementary Fig. S2) performed with the *R* package *HardyWeinberg* v1.6.3^107, 108^. Allelic richness, expected (*H*_E_), unbiased expected (u*H*_E_) and observed heterozygosity (*H*_O_) as well as the *F*-statistics were measured for all European bison individuals and for each breeding line. Molecular based heterozygosities and *F*-statistics (*F*_IT_, *F*_ST_, *F*_IS_) were calculated in *GenAlex* and *FSTAT* v2.9.4^109^.

*PMx*^110^ was used to generate genetic values from pedigree data. *PMx* provides two methods to calculate pedigree-based gene diversity (GD): from kinship matrix as well as gene drop method^111^. For the latter method genetic default assumptions (1 000 gene drop iterations, autosomal mendelian inheritance mode) were used. GD is equivalent to *H*_E_ ^111, 112^ and was therefore used for pedigree versus molecular data comparisons. For clarification and as it is output by each software, GD will always refer to the pedigree-based values within this study, whereas *H*_E_ is referring to molecular-based values. Additionally, pedigree-based *F*_ST_, *F*_IS_ and *F*_IT_ were generated in *ENDOG* v4.8^113^.

The pedigree-based and SNP-based *F-*statistics were also compared. In order to do this, two pedigree data sets were used for *PMx*: for a direct comparison the pedigree-based genetic values were computed including only the successfully SNP-genotyped individuals with known genealogy (*n* = 99) and their assigned ancestors (*n* = 982) up to the founders. To evaluate the representativeness of those pedigree-based genetic values, the same calculations were conducted with all sampled individuals with known genealogy in this study (*n* = 227) and their assigned ancestors up to the founders (*n* = 1,296).

### Visualisation and data set conversion

Boxplots were generated with the *R* packages *ggplot2* v3.2.0^114^ and *gridExtra* v2.3^115^. The cladogram of the Bovini and other non-target species was conducted in *Mesquite* v3.61 (build 927)^116^. *CONVERT* v1.31^117^ was used to adjust data sets for implementation in several analysis programs. The *R* package *genetics* v1.3.8.1.2^118^ was used to transform data sets into partly required genotype data sets.

## Supporting information

SNP marker list

Genotype lists

Sample list

## Acknowledgments

Funding for this study was generated by the Hessen State Ministry of Higher Education, Research and the Arts (HMWK) via the LOEWE Centre for Translational Biodiversity Genomics (LOEWE-TBG) and through the centre for wildlife genetics at the Senckenberg Conservation Genetics Section, Gelnhausen. G.W. received generous funding from the Karl und Marie Schack-Stiftung. Special thanks to the regional EBCCs represented by the Wisentgehege und Falkenhof in Springe, Wisentgehege in Warburg-Hardehausen, Wisentreservat Damerower Werder and the Wisentprojekt Donaumoos. Furthermore, we thank several sample providers: Wisent-Welt Wittgenstein e.V., Wildpark Alte Fasanerie Hanau-Klein-Auheim, Wilhelma in Stuttgart, Tier-und Pflanzenpark Fasanerie Wiesbaden, Hanover Zoo, Zoologischer Garten Köln, Wildpark Köln-Dünnwald, Natuurpark and Het Flevo-Landschap in Lelystad, Duisburg Zoo, Tierpark Gera, Tiergarten Bernburg, Zoological Garden Berlin, Tierpark Berlin-Friedrichsfelde, Serengenti-Park Hodenhagen, Rostock Zoo, Bavarian Forest National Park, Alpenzoo in Innsbruck, Tierpark Suhl, Wisentpark Kropp, Lower Oder Valley National Park, faunaforst in Bryrup, Krefeld Zoo, Tiergarten Nürnberg, Dierenpark Planckendael in Mechelen, Tierpark Chemnitz, Wildgatter Oberrabenstein, Tierpark Neumünster, Opel-Zoo in Kronberg, Wildtierpark Edersee in Edertal-Hemfurth, SafariZoo in Thoiry, Tierpark Oberwald, Karlsruhe Zoo, Wildpark Leipzig-Connewitz, Warszawa Zoo, Tierpark Sababurg in Hofgeismar and Praha Zoo.

## Author contributions

This study was designed by G.W., C.N. and B.C. Sampling and sample organisation were done by G.W., M.T., B.C. and C.N. Laboratory work was performed by G.W. under B.C. supervision. All microsatellite and SNP data were generated and scored by G.W and B.C. Data analyses were performed by G.W. G.W. wrote the original manuscript draft. All authors contributed to the preparation of the final draft and approved it.

## Additional information

### Data availability statement

The authors confirm that the data supporting the findings of this study are available within the published article and its supplementary materials.

## Legends of figures and tables

**Figure 7: Probability of identity (PID) and probability of identity among siblings (PIDsib) of genotyped microsatellites (n = 11) and autosomal SNPs (n = 95) for European bison.**

Horizontal dashed red line: PID threshold for natural populations by Waits et al.^17^ is not overcome by the microsatellite panel. SNP-based PID reaches threshold at approx. 10, PIDsib at approx. 18 loci for the European bison. Approximations of PID and PIDsib close to zero are reached approx. with 13 and 24 loci, respectively. The x-axis was cut at locus combination of 30 loci for more conciseness whereby the approximation of the SNP-based PIDs does not change after 30 loci. PIDsibs estimations of the microsatellite panel are outside of the scale. PID and PIDsib for all other Bos species for which individualisation was possible based on 95 autosomal SNPs are provided in Supplementary Fig. S4.

**Figure 8: Detected number of mean allele mismatches between individual genotypes (genotypes consisting of 95 loci) of European bison (both breeding lines separately) as well as American bison, domestic cattle, gaur and banteng.**

Lowest allele mismatches are highlighted in red. Individual sample size per group is noted (*n*). Allele mismatches between genotypes of five unrelated cattle individuals are > 40 loci.

**Figure 9:** An exemplary family network to document the integration of molecular kinship analysis into the present pedigree data from the European Bison Pedigree Book (EBPB).

Three generations of 23 individuals assigned to LL were sampled and genotyped from three holders in the Netherlands and Germany (Lelystad (Natuurpark), Duisburg (Zoo), and Springe (Wisentgehege)). Circles represent female individuals and squares male individuals (filled symbols: genotyped). Green edges around the individuals represent successful molecular sex verification, whereas solid red edges represent falsely positive sex assignments and dashed red edges, where no molecular sex assignment was possible. All genotypes are based on a single sample per individual. Triple edges: sample was not individually assignable in the field but was assigned to an individual with the genotype based on sex determination and parental assignment. Different colours of the genealogical lineages represent different verification states: green: genetically verified kinships from the EBPB; blue: genetically assigned kinships with lacking data in the EBPB; red: kinship from the EBPB not genetically verified; black: kinships genetically not verifiable due to missing genotypes. 10 parental assignments (sired by ‘EBPB#7591’ and ‘EBPB#10081’) with unknown maternities from the EBPB were included to visualise the high degree of at least half-sibling relationships of the females/potential mothers in Lelystad; grey dashed: presumed kinships not verifiable due to missing genotypes and missing data in the EBPB. Asterisk: case of inbreeding. All breeding line assignments of the displayed individuals were genetically verified (not noted here).

**Table 2: Genetic diversity measures based on SNP genotypes and pedigree data for different sets of European bison individuals.**

SNP genotype values are based on unlinked 63 SNPs in HWE. All 277 of 338 sampled individuals with known genealogy were used to generate pedigree-based genetic values. As genealogical information was not available for all successfully genotyped individuals, molecular and pedigree-based genetic diversity values were calculated for an overlapping set of 99 successfully SNP-genotyped individuals with available genealogical data. Sample sizes [n] in squared brackets show the number of individuals included in the associated pedigree up to the founders. Values in parentheses next to the genetic values represent the associated standard errors (SE). F-statistics were calculated using either arithmetic averages^1^ or based on the average H_S_ and H_T_ over loci^2^. Pedigree-based genetic diversity values in PMx were calculated based on kinship matrix^3^ or gene drop^4^. A more detailed table including genetic diversity values of each both breeding lines is provided in the Supplementary Table S4.

**Figure 10:** Assignment probabilities [%] based on 18 loci selected for breeding line discrimination between LC (*n* = 76) and LL (*n* = 61) in the European bison:

(a) Bayesian genetic clustering computed with *STRUCTURE*; (b) Maximum-likelihood genetic clustering computed with *adegenet*. The black line shows the previously assigned lineage distinction (LC: blue; LL: orange). Dashed red lines indicate assignment thresholds. Bars tarnished red mark individuals with unexpected lineage assignment; bars tarnished grey mark individuals not assignable with genotypic data according to the assignment threshold. Brown arrows: F_1_ breeding line hybrids. White asterisks: LC individuals with at least one of the six private alleles found in LC. See Supplementary Table S5 for the order of individuals shown here.

**Figure 11:** SNP call rate [%] for 95 autosomal SNPs in the European bison and 15 non-target species with corresponding numbers of individuals (*n*). The length of a solid bar indicates the mean SNP call rate for each analysed species. Blue bars reflect all groups classified to the genus *Bos*, blue-grey bars groups classified to the subtribe Bubalina and grey bars species outside of Bovini. A SNP call rate of at least 80 % call rate (red dashed line) is the threshold for inclusion into further analysis. The orange-hatched bars show the percentage of found polymorphism over 95 loci within the groups. The cladogram reflects known evolutionary relationships between the species^18^. The asterisk points out the tribe of Bovini.

**Figure 12:** (a) PCoA of 137 European bison (both breeding lines) and 116 individuals of 10 non-target Bovini species (16 ESUs) with a SNP call rate over 80 % utilising all 95 autosomal SNP loci. (b) PCoA of 137 European bison and 15 domestic cattle (four major lineages) utilising a subset of selected 31 SNP loci.

Clusters containing higher taxa like the subgenus *Bibos* (HODGSON, 1837) and the subtribe Bubalina (RÜTIMEYER, 1865) are marked in grey circles. Eigenvalues (a): axis 1: 233.68; axis 2: 158.19; Eigenvalues (b): axis 1: 33.46; axis 2: 26.85

## References

1. Krasińska, M. & Krasiński, Z. A. European bison: The nature monograph. (Springer Berlin Heidelberg, 2013).

2. Raczyński, J. European Bison Pedigree Studbook. (2021).

3. Wojciechowska, M. et al. From Wisent to the Lab and Back Again—A Complex SNP Set for Population Management as an Effective Tool in European Bison Conservation. Diversity 15, 116 (2023).

4. Slatis, H. M. An Analysis of Inbreeding in the European Bison. Genetics 45, 275–87 (1960).

5. European bison: Status survey and conservation action plan. (IUCN, 2004).

6. Tokarska, M., Pertoldi, C., Kowalczyk, R. & Perzanowski, K. Genetic status of the European bison Bison bonasus after extinction in the wild and subsequent recovery. Mammal Review 41, 151– 162 (2011).

7. Kuemmerle, T., Hickler, T., Olofsson, J., Schurgers, G. & Radeloff, V. C. Reconstructing range dynamics and range fragmentation of European bison for the last 8000 years. Diversity and Distributions 18, 47–59 (2012).

8. Gautier, M. et al. Deciphering the Wisent Demographic and Adaptive Histories from Individual Whole-Genome Sequences. Mol Biol Evol 33, 2801–2814 (2016).

9. Belousova, I. P. & Kudriavtsev, I. V. Genetic structure of captive and free-living European bison populations through Pedigree analysis. Zeitschrift für Säugetierkunde; Proceedings of the 1st International Symposium on Physiology an Ethology of Wild and Zoo Animals; Supplementum II 62, 12–13 (1997).

10. Willi, Y., van Buskirk, J. & Hoffmann, A. A. Limits to the Adaptive Potential of Small Populations. Annu. Rev. Ecol. Evol. Syst. 37, 433–458 (2006).

11. Tokarska, M. et al. Effectiveness of microsatellite and SNP markers for parentage and identity analysis in species with low genetic diversity: The case of European bison. Heredity (Edinb) 103, 326–32 (2009).

12. Taberlet, P., Waits, L. P. & Luikart, G. Noninvasive genetic sampling: Look before you leap. Trends in Ecology & Evolution 14, 323–327 (1999).

13. Mills, L. S., Citta, J. J., Lair, K. P., Schwartz, M. K. & Tallmon, D. A. Estimating animal abundance using noninvasive DNA sampling: promise and pitfalls. Ecological Applications 10, 283–294 (2000).

14. Eggert, L. S., Eggert, J. A. & Woodruff, D. S. Estimating population sizes for elusive animals: the forest elephants of Kakum National Park, Ghana. Mol Ecol 12, 1389–1402 (2003).

15. Piggott, M. P. & Taylor, A. C. Remote collection of animal DNA and its applications in conservation management and understanding the population biology of rare and cryptic species. Wildl. Res. 30, 1 (2003).

16. Waits, L. P. & Paetkau, D. Noninvasive Genetic Sampling Tools for Wildlife Biologists:: A Review of Applications and Recommendations for Accurate Data Collection. Journal of Wildlife Management 69, 1419–1433 (2005).

17. Waits, L. P., Luikart, G. & Taberlet, P. Estimating the probability of identity among genotypes in natural populations: cautions and guidelines. Mol Ecol 10, 249–56 (2001).

18. The genetics of cattle. (CAB International, 2015).

19. Kraus, R. H. S. et al. A single-nucleotide polymorphism-based approach for rapid and cost-effective genetic wolf monitoring in Europe based on noninvasively collected samples. Mol Ecol Resour 15, 295–305 (2015).

20. von Thaden, A. et al. Assessing SNP genotyping of noninvasively collected wildlife samples using microfluidic arrays. Sci Rep 7, 10768 (2017).

21. Groves, C. P., et al. Family Bovidae (Hollow-horned Ruminants). in Hoofed mammals (ed. Mittermeier, R. A.) vol. 2 (Lynx, 2011).

22. Felius, M. Cattle breeds: An encyclopedia. (Misset, 1995).

23. Borghese, A. & Mazzi, M. Buffalo population and strategies in the world. Buffalo production and research 67, 1–39 (2005).

24. Nussberger, B., Wandeler, P. & Camenisch, G. A SNP chip to detect introgression in wildcats allows accurate genotyping of single hairs. Eur J Wildl Res 60, 405–410 (2014).

25. von Thaden, A. et al. Applying genomic data in wildlife monitoring: Development guidelines for genotyping degraded samples with reduced single nucleotide polymorphism panels. Mol Ecol Resour (2020) doi:10.1111/1755-0998.13136.

26. Taberlet, P. et al. Noninvasive genetic tracking of the endangered Pyrenean brown bear population. Mol Ecol 6, 869–876 (1997).

27. Morin, P. A., Luikart, G., Wayne, R. K. & group, the S. workshop. SNPs in ecology, evolution and conservation. Trends in Ecology & Evolution 19, 208–216 (2004).

28. Wojciechowska, M. et al. Panel of informative SNP markers for two genetic lines of European bison: Lowland and Lowland-Caucasian. ANIMAL BIODIVERSITY AND CONSERVATION 40, 17–25 (2017).

29. Oleński, K., Kamiński, S., Tokarska, M. & Hering, D. M. Subset of SNPs for parental identification in European bison Lowland-Białowieża line (Bison bonasus bonasus). Conservation Genet Resour 10, 73–78 (2018).

30. Jones, O. R. & Wang, J. Molecular marker-based pedigrees for animal conservation biologists. Animal Conservation 13, 26–34 (2010).

31. Taylor, H. R., Kardos, M. D., Ramstad, K. M. & Allendorf, F. W. Valid estimates of individual inbreeding coefficients from marker-based pedigrees are not feasible in wild populations with low allelic diversity. Conserv Genet 16, 901–913 (2015).

32. Mueller, S. A. et al. The rise of a large carnivore population in Central Europe: genetic evaluation of lynx reintroduction in the Harz Mountains. Conserv Genet 21, 577–587 (2020).

33. Sipko, T. P. European bison in Russia-past, present and future. European Bison Conservation Newsletter 2, 148–159 (2009).

34. Kamiński, S., Olech, W., Oleński, K., Nowak, Z. & Ruść, A. Single nucleotide polymorphisms between two lines of European bison (Bison bonasus) detected by the use of Illumina Bovine 50 K BeadChip. Conservation Genet Resour 4, 311–314 (2012).

35. Witzenberger, K. A. & Hochkirch, A. Ex situ conservation genetics: a review of molecular studies on the genetic consequences of captive breeding programmes for endangered animal species. Biodivers Conserv 20, 1843–1861 (2011).

36. Pertoldi, C. et al. Depauperate genetic variability detected in the American and European bison using genomic techniques. Biol Direct 4, 4848 (2009).

37. Tokarska, M., Kawałko, A., Wójcik, J. M. & Pertoldi, C. Genetic variability in the European bison (Bison bonasus) population from Białowieża forest over 50 years. Biol J Linn Soc Lond 97, 801– 809 (2009).

38. Pertoldi, C. et al. Genome variability in European and American bison detected using the BovineSNP50 BeadChip. Conserv Genet 11, 627–634 (2010).

39. Oleński, K. et al. A refined genome-wide association study of posthitis in lowland Białowieza population of the European bison (Bison bonasus). Eur J Wildl Res 66, 6410 (2020).

40. Launhardt, K., Epplen, C., Epplen, J. T. & Winkler, P. Amplification of microsatellites adapted from human systems in faecal DNA of wild Hanuman langurs (Presbytis entellus). Electrophoresis 19, 1356–61 (1998).

41. Smith, K. L. et al. Cross-species amplification, non-invasive genotyping, and non-Mendelian inheritance of human STRPs in Savannah baboons. Am. J. Primatol. 51, 219–227 (2000).

42. Ogden, R., Baird, J., Senn, H. & McEwing, R. The use of cross-species genome-wide arrays to discover SNP markers for conservation genetics: a case study from Arabian and scimitar-horned oryx. Conservation Genet Resour 4, 471–473 (2012).

43. Albrechtsen, A., Nielsen, F. C. & Nielsen, R. Ascertainment biases in SNP chips affect measures of population divergence. Mol Biol Evol 27, 2534–47 (2010).

44. Malomane, D. K. et al. Efficiency of different strategies to mitigate ascertainment bias when using SNP panels in diversity studies. BMC Genomics 19, 22 (2018).

45. Schlötterer, C. & Harr, B. Single nucleotide polymorphisms derived from ancestral populations show no evidence for biased diversity estimates in Drosophila melanogaster. Mol Ecol 11, 947– 950 (2002).

46. Druet, T. et al. Genomic footprints of recovery in the European bison. J Hered (2020) doi:10.1093/jhered/esaa002.

47. Wang, J. Pedigrees or markers: Which are better in estimating relatedness and inbreeding coefficient? Theor Popul Biol 107, 4–13 (2016).

48. Rudnick, J. A. & Lacy, R. C. The impact of assumptions about founder relationships on the effectiveness of captive breeding strategies. Conserv Genet 9, 1439–1450 (2008).

49. Olech, W. European bison EEP Annual Report 2004. in EAZA Yearbook 2004 (eds. van Lint, W., de Man, D., Garn, K., Hiddinga, B. & Brouwer, K.) 529–531 (2006).

50. Olech, W. European bison EEP Annual Report 2005. in EAZA Yearbook 2005 (eds. de Man, D., van Lint, W., Garn, K. & Hiddinga, B.) 561–564 (2007).

51. Gutiérrez, J. P., Goyache, F. & Cervantes, I. User’s Guide: ENDOG v4.8: A Computer Program for Monitoring Genetic Variability of Populations Using Pedigree Information. (2010).

52. Harris, A. M. & DeGiorgio, M. An Unbiased Estimator of Gene Diversity with Improved Variance for Samples Containing Related and Inbred Individuals of any Ploidy. G3 (Bethesda) 7, 671–691 (2017).

53. Wahlund, S. Zusammensetzung von Populationen und Korrelationserscheinungen vom Standpunkt der Vererbungslehre ausbetrachtet. Hereditas 11, 65–106 (1928).

54. Frankham, R., Ballou, J. D. & Briscoe, D. A. Introduction to conservation genetics. (Univ. Press, 2015).

55. Boyles, R., Schutz, E. & de Leon, J. Bubalus mindorensis: The IUCN Red List of Threatened Species 2016: e.T3127A50737640. (2016).

56. Burton, J., Wheeler, P. & Mustari, A. Bubalus depressicornis: The IUCN Red List of Threatened Species 2016: e.T3126A46364222. (2016).

57. Burton, J., Wheeler, P. & Mustari, A. Bubalus quarlesi: The IUCN Red List of Threatened Species 2016: e.T3128A46364433. (2016).

58. Buzzard, P. & Berger, J. Bos mutus: The IUCN Red List of Threatened Species 2016: e.T2892A101293528. (2016).

59. Duckworth, J. W., Sankar, K., Williams, A. C., Samba Kumar, N. & Timmins, R. J. Bos gaurus: The IUCN Red List of Threatened Species 2016: e.T2891A46363646. (2016).

60. Gardner, P., Hedges, S., Pudyatmoko, S., Gray, T. N. E. & Timmins, R. J. Bos javanicus: The IUCN Red List of Threatened Species 2016: e.T2888A46362970. (2016).

61. Timmins, R. J., Burton, J. & Hedges, S. Bos sauveli: The IUCN Red List of Threatened Species 2016: e.T2890A46363360. (2016).

62. Aune, K., Jørgensen, D. & Gates, C. C. Bison bison: The IUCN Red List of Threatened Species 2017: e.T2815A123789863. (2018).

63. IUCN SSC Antelope Specialist Group. Syncerus caffer: The IUCN Red List of Threatened Species 2019: e.T21251A50195031. (2019).

64. Kaul, R., Williams, A. C., Rithe, K., Steinmetz, R. & Mishra, R. Bubalus arnee: The IUCN Red List of Threatened Species 2019: e.T3129A46364616. (2019).

65. Plumb, G., Kowalczyk, R. & Hernandez-Blanco, J. A. IUCN Red List of Threatened Species 2020: Bison bonasus. (2020). doi:10.2305/IUCN.UK.2020-3.RLTS.T2814A45156279.en.

66. Timmins, R. J., Hedges, S. & Robichaud, W. Pseudoryx nghetinhensis: The IUCN Red List of Threatened Species 2020: e.T18597A166485696. (2020).

67. Wilson, G. A., Nishi, J. S., Elkin, B. T. & Strobeck, C. Effects of a recent founding event and intrinsic population dynamics on genetic diversity in an ungulate population. Conserv Genet 6, 905–916 (2006).

68. Sutherland, W. J., Pullin, A. S., Dolman, P. M. & Knight, T. M. The need for evidence-based conservation. Trends in Ecology & Evolution 19, 305–8 (2004).

69. Brooks, J. S., Franzen, M. A., Holmes, C. M., Grote, M. N. & Mulder, M. B. Testing hypotheses for the success of different conservation strategies. Conservation Biology 20, 1528–38 (2006).

70. Schröder, F., Oldorf, M. A. P. & Heising, K. L. Spatial relation between open landscapes and debarking hotspots by European bison (Bison bonasus) in the Rothaar Mountains. European Bison Conservation Newsletter 12, 5–16 (2019).

71. Hagemann, L. et al. Long-term inference of population size and habitat use in a socially dynamic population of wild western lowland gorillas. Conserv Genet 143, 1780 (2019).

72. Giglio, R. M., Ivy, J. A., Jones, L. C. & Latch, E. K. Pedigree-based genetic management improves bison conservation. Jour. Wild. Mgmt. 82, 766–774 (2018).

73. Olech, W. & Perzanowski, K. A genetic background for reintroduction program of the European bison (Bison bonasus) in the Carpathians. Biological Conservation 108, 221–228 (2002).

74. Pertoldi, C. et al. Phylogenetic relationships among the European and American bison and seven cattle breeds reconstructed using the BovineSNP50 Illumina Genotyping BeadChip. Acta Theriologica 55, 97–108 (2010).

75. Rowe, K. C. et al. Museum genomics: low-cost and high-accuracy genetic data from historical specimens. Mol Ecol Resour 11, 1082–92 (2011).

76. Hoffmann, G. S. & Griebeler, E. M. An improved high yield method to obtain microsatellite genotypes from red deer antlers up to 200 years old. Mol Ecol Resour 13, 440–6 (2013).

77. Hoffmann, G. S., Johannesen, J. & Griebeler, E. M. Population dynamics of a natural red deer population over 200 years detected via substantial changes of genetic variation. Ecol Evol 6, 3146–53 (2016).

78. Oleński, K. et al. Genome-wide association study for posthitis in the free-living population of European bison (Bison bonasus). Biol Direct 10, 2 (2015).

79. Kunvar, S., Czarnomska, S., Pertoldi, C. & Tokarska, M. In Search of Species-Specific SNPs in a Non-Model Animal (European Bison (Bison bonasus))-Comparison of De Novo and Reference-Based Integrated Pipeline of STACKS Using Genotyping-by-Sequencing (GBS) Data. Animals (Basel) 11, (2021).

80. R Core Team. R: A language and environment for statistical computing. (2019).

81. RStudio Team. RStudio: Integrated Development Environment for R. (2016).

82. Jansson, M., Ståhl, I. & Laikre, L. mPed: a computer program for converting pedigree data to a format used by the PMx-software for conservation genetic analysis. Conservation Genet Resour 5, 651–653 (2013).

83. Ballou, J. D., Lacy, R. C. & Pollak, J. P. PMx: Software for demographic and genetic analysis and management of pedigreed populationsChicago. (2018).

84. Klös, H.-G. & Wünschmann, A. Die Rinder. in Säugetiere 4 (eds. Bannikow, A. G., et al.) vol. 13 368–436 (Deutscher-Taschenbuch-Verl., 1993).

85. Mwai, O., Hanotte, O., Kwon, Y.-J. & Cho, S. African Indigenous Cattle: Unique Genetic Resources in a Rapidly Changing World. Asian-australas J Anim Sci 28, 911–21 (2015).

86. Kumar, S. et al. Mitochondrial DNA analyses of Indian water buffalo support a distinct genetic origin of river and swamp buffalo. Anim Genet 38, 227–32 (2007).

87. Yindee, M. et al. Y-chromosomal variation confirms independent domestications of swamp and river buffalo. Anim Genet 41, 433–5 (2010).

88. Valiere, N. & Taberlet, P. Urine collected in the field as a source of DNA for species and individual identification. Mol Ecol 9, 2150–2152 (2003).

89. Westekemper, K., Signer, J., Cocchiararo, B., Nowak, C. & Balkenhol, N. Understanding effective isolation of intensively managed red deer populations across Germany.

90. Aasen, E. & Medrano, J. F. Amplification of the Zfy and Zfx Genes for Sex Identification in Humans, Cattle, Sheep and Goats. Nat Biotechnol 8, 1279–1281 (1990).

91. Wang, J. et al. High-throughput single nucleotide polymorphism genotyping using nanofluidic Dynamic Arrays. BMC Genomics 10, 561 (2009).

92. Nguyen, T. T. et al. Phylogenetic position of the saola (Pseudoryx nghetinhensis) inferred from cytogenetic analysis of eleven species of Bovidae. Cytogenet Genome Res 122, 41–54 (2008).

93. Peakall, R. & Smouse, P. E. GenAlEx 6.5: genetic analysis in Excel. Population genetic software for teaching and research--an update. Bioinformatics 28, 2537–9 (2012).

94. Shin, J.-H., Blay, S., Graham, J. & McNeney, B. LDheatmap : An R Function for Graphical Display of Pairwise Linkage Disequilibria Between Single Nucleotide Polymorphisms. J. Stat. Soft. 16, (2006).

95. Mammal species of the world: A taxonomic and geographic reference. (Johns Hopkins Univ. Press, 2005).

96. Handbook of the mammals of the world. (Lynx, 2009).

97. Jones, O. R. & Wang, J. COLONY: a program for parentage and sibship inference from multilocus genotype data. Mol Ecol Resour 10, 551–5 (2010).

98. Wang, J. Computationally efficient sibship and parentage assignment from multilocus marker data. Genetics 191, 183–94 (2012).

99. Pritchard, J. K., Stephens, M. & Donnelly, P. Inference of population structure using multilocus genotype data. Genetics 155, 945–59 (2000).

100. Falush, D., Stephens, M. & Pritchard, J. K. Inference of population structure using multilocus genotype data: linked loci and correlated allele frequencies. Genetics 164, 1567–87 (2003).

101. Pritchard, J. K., Wen, X. & Falush, D. Documentation for structure software: Version 2.3. (2010).

102. Earl, D. A. & von Holdt, B. M. STRUCTURE HARVESTER: a website and program for visualizing STRUCTURE output and implementing the Evanno method. Conservation Genet Resour 4, 359– 361 (2012).

103. Jakobsson, M. & Rosenberg, N. A. CLUMPP: a cluster matching and permutation program for dealing with label switching and multimodality in analysis of population structure. Bioinformatics 23, 1801–6 (2007).

104. Beugin, M.-P., Gayet, T., Pontier, D., Devillard, S. & Jombart, T. A fast likelihood solution to the genetic clustering problem. Methods Ecol Evol 9, 1006–1016 (2018).

105. Jombart, T. adegenet: a R package for the multivariate analysis of genetic markers. Bioinformatics 24, 1403–5 (2008).

106. Jombart, T. & Ahmed, I. adegenet 1.3-1: new tools for the analysis of genome-wide SNP data. Bioinformatics 27, 3070–1 (2011).

107. Graffelman, J. & Camarena, J. M. Graphical tests for Hardy-Weinberg equilibrium based on the ternary plot. Hum Hered 65, 77–84 (2008).

108. Graffelman, J. Exploring Diallelic Genetic Markers: The HardyWeinberg Package. J. Stat. Soft. 64, (2015).

109. Goudet, J. Fstat: a program to estimate and test population genetics parameters. (2003).

110. Lacy, R. C., Ballou, J. D. & Pollak, J. P. PMx: software package for demographic and genetic analysis and management of pedigreed populations. Methods Ecol Evol 3, 433–437 (2012).

111. PMx Users Manual Version 1.0. (2011).

112. Nei, M. Analysis of gene diversity in subdivided populations. Proceedings of the National Academy of Sciences 70, 3321–3 (1973).

113. Gutiérrez, J. P. & Goyache, F. A note on ENDOG: a computer program for analysing pedigree information. J Anim Breed Genet 122, 172–6 (2005).

114. Wickham, H. ggplot2. (Springer International Publishing, 2016). doi:10.1007/978-3-319-24277-4.

115. Auguie, B. & Antonov, A. gridExtra. (2017).

116. Maddison, W. P. & Maddison, D. R. Mesquite: a modular system for evolutionary analysis. (2019).

117. Glaubitz, J. C. convert: A user-friendly program to reformat diploid genotypic data for commonly used population genetic software packages. Mol Ecol Notes 4, 309–310 (2004).

118. Warnes, G. genetics: Population Genetics (R package). (2012).

