## Supplementary Information for "A reduced SNP panel optimised for non-invasive genetic assessment of a genetically impoverished conservation icon, the European bison"

#### Index

#### Index for supplementary tables

Supplementary Table S5: Assignment probabilities [%] based on 18 loci selected for breeding line discrimination between the LC (n = 76) and LL line (n = 61) in the European bison. Two methods are compared: Bayesian genetic clustering computed with STRUCTURE; Maximum-likelihood genetic clustering computed with adegenet. Individuals (EBPB# and study internal names) are ordered within their breeding line (according to the metadata) after assignment probabilities computed with the Bayesian clustering (same order as in the barplot (Fig. 4)). .......... 14
Supplementary Table S6: Overview of all utilised microsatellite markers. .......... 39
Supplementary Table S7: Overview of primer concentrations in the multiplex mixes A to C (µl/reaction) and ratios of the not labelled forward primers (For) and fluorescence-labelled forward primers (ForLab). .......... 41
Supplementary Table S8: Protocol adjustments for preparing the 10X STA Primers for SNP genotyping. Font colour code: grey = protocols during testing phase; black = protocols for general genotyping of this 96× SNP panel (recommended for adopting in other laboratoires). Abbreviations: LSP = Locus-specific primer (Rev); STA = Specific target amplification primer. Nomenclature was taken from Fluidigm protocol for comparison. For not adjusted protocols see manufacture's documentation for SNPtypeTM Assays for SNP Genotyping (Advanced Development Protocol 34, Fluidigm corp.). .. 45
Supplementary Table S9: Protocol adjustments for preparing the 10X STA Primers for SNP genotyping. Colour code: grey = protocols during testing phase; black = protocols for general genotyping of this 96× SNP panel (recommended for adopting in other laboratoires). Abbreviations: STA = Specific target amplification primer; gDNA = genomic DNA. Nomenclature was taken from Fluidigm protocol for comparison. For not adjusted protocols see manufacture's documentation for SNPtypeTM Assays for SNP Genotyping (Advanced Development Protocol 34, Fluidigm corp.). ........ 46
Supplementary Table S10: Protocol adjustments for preparing the 10X STA Primers for SNP genotyping. Colour code: grey = protocols during testing phase; black = protocols for general genotyping of this 96× SNP panel (recommended for adopting in other laboratoires). Abbreviations: ASP = Allele-Specific Primers; LSP = Locus-specific primer (Rev). Nomenclature was taken from Fluidigm protocol for comparison. For not adjusted protocols see manufacture's documentation for SNPtypeTM Assays for SNP Genotyping (Advanced Development Protocol 34, Fluidigm corp.). ........ 47

### Index for supplementary figures

Supplementary Figure S1: Pie charts of founder representations in (a) all 337 sampled and (b) all genotyped individuals with pedigree information based on genealogy documented in the EBPB. Brown: Founders of both breeding lines; green: founders exclusive for the LC line. Darker colours: males; lighter colours: females. Detached pie piece resembles the Caucasian bison founder 'Kaukasus' (EBPB#100). The founder representations found in our comprehensive sample set are congruent with Tokarska et al.[1] for the global in situ and ex situ populations.................................... 5
Supplementary Figure S2: Ternary Hardy-Weinberg-Equilibrium (HWE) plots showing heterozygote deficiencies or excesses of 90 autosomal markers of 58 non-first-degree relatives of European bison.8

### Overview of the supplementary material

**Supplementary Table S1: Overview of all additional digital files of the supplementary material with short content descriptions.**

| File name | content |
|---|---|
| **Supplementary Information** | Contains additional Material & Methods, Results and Discussion in text, tables, and figures (the document open at the moment) |
| **Sample_list.xlsx** | Complete sampling list of all samples collected and/or used in the framework of this study with individual details |
| **Genotype_list.xlsx** | Complete SNP genotype (96 loci) and microsatellite genotype (11 loci) lists with individual details |
| **SNP_marker_list_details.xlsx** | List of all SNP markers tested in this study with further details and additional data for all 96 loci of the final panel. Details on sex marker design. |

### Supplementary Results

#### Pedigree data

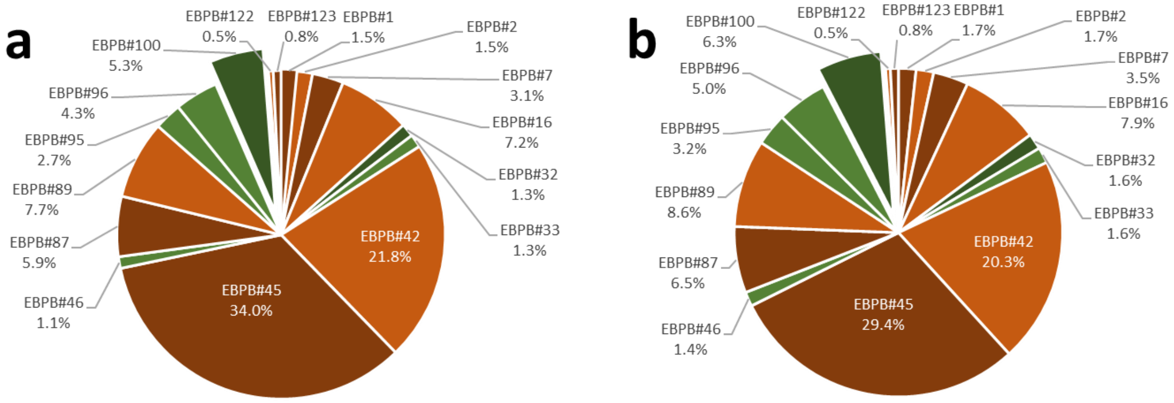

**Supplementary Figure S1: Pie charts of founder representations in (a) all 337 sampled and (b) all genotyped individuals with pedigree information based on genealogy documented in the EBPB.** Brown: Founders of both breeding lines; green: founders exclusive for LC. Darker colours: males; lighter colours: females. Detached pie piece resembles the Caucasian bison founder 'Kaukasus' (EBPB#100). The founder representations found in our comprehensive sample set are congruent with Tokarska *et al.*[1] for the global *in situ* and *ex situ* populations.

#### Microsatellites

Nearly all microsatellites used in wisent genetics so far are in non-coding regions. Only Flisikowski *et al.*[2] used a not named microsatellite located in the growth hormone receptor (GHR) gene given by Lucy *et al.*[3]. Mikhailova and Voitsukhovskaya[4] did not published their microsatellite marker set and therefore cannot compared with other studies.

Seven of 21 microsatellite markers were rejected by homozygosity or non-function: four microsatellites (NVHRT48, NVHRT21, CER14, INRA35) by non-function in *B. bonasus* and three (CSSM66, RT1, ETH225) were not suitable for the non-invasive approach. The gonosomal and twelve autosomal polymorphic microsatellites were viable for the analysis of European bison samples (Supplementary Table S2). IDVGA55 was the only monomorphic marker of the microsatellites applicable for the non-invasive approach and only used in evaluation of faecal sampling methodology.

**Supplementary Table S2: Characterisation of tested microsatellite markers appropriate for non-invasive samples from *Bos bonasus* found in 51 individuals from both breeding lines (LL: *n* = 22; LC: *n* = 29).** Allele frequencies are given for all individuals (wisent) and separately for both breeding lines (LC, LL). Private alleles per breeding line in the genotyped individuals are underlined. In two individuals IDVGA59 could not be successfully amplified and scored. In these both cases only non-invasive samples were available.

| *Locus* | *Multiplex* | *Allele* | | *wisent* | *LC* | *LL* |
|---|---|---|---|---|---|---|
| *DIK082* | **A** | | **N** | **51** | **22** | **29** |
| | | **100** | | 0.363 | 0.341 | 0.379 |
| | | **<u>112</u>** | | 0.039 | 0.000 | <u>0.069</u> |
| | | **124** | | 0.598 | 0.659 | 0.552 |
| *IDVGA59* | **A** | | **N** | **49** | **20** | **29** |
| | | **244** | | 0.265 | 0.275 | 0.259 |
| | | **264** | | 0.735 | 0.725 | 0.741 |
| *BM4208* | **A** | | **N** | **51** | **22** | **29** |
| | | **158** | | 0.167 | 0.114 | 0.207 |
| | | **160** | | 0.833 | 0.886 | 0.793 |
| *BM203* | **B** | | **N** | **51** | **22** | **29** |
| | | **218** | | 0.971 | 0.955 | 0.983 |
| | | **222** | | 0.029 | 0.045 | 0.017 |
| *CSSM19* | **B** | | **N** | **51** | **22** | **29** |
| | | **140** | | 0.118 | 0.182 | 0.069 |
| | | **142** | | 0.569 | 0.568 | 0.569 |
| | | **148** | | 0.314 | 0.250 | 0.362 |
| *CSSM14* | **B** | | **N** | **51** | **22** | **29** |
| | | **134** | | 0.627 | 0.614 | 0.638 |
| | | **136** | | 0.088 | 0.023 | 0.138 |
| | | **138** | | 0.284 | 0.364 | 0.224 |
| *CSSM22* | **B** | | **N** | **51** | **22** | **29** |
| | | **214** | | 0.990 | 0.977 | 1.000 |
| | | **<u>216</u>** | | 0.010 | <u>0.023</u> | 0.000 |
| *BMC1009* | **B** | | **N** | **51** | **22** | **29** |
| | | **276** | | 0.941 | 0.932 | 0.948 |
| | | **<u>278</u>** | | 0.010 | <u>0.023</u> | 0.000 |
| | | **280** | | 0.049 | 0.045 | 0.052 |
| *BM1818* | **B** | | **N** | **51** | **22** | **29** |
| | | **260** | | 0.480 | 0.614 | 0.379 |
| | | **264** | | 0.520 | 0.386 | 0.621 |
| *MM12* | **C** | | **N** | **51** | **22** | **29** |
| | | **108** | | 0.382 | 0.318 | 0.431 |
| | | **110** | | 0.618 | 0.682 | 0.569 |
| *Haut14* | **C** | | **N** | **51** | **22** | **29** |
| | | **142** | | 0.559 | 0.523 | 0.586 |
| | | **144** | | 0.441 | 0.477 | 0.414 |
| *CSSM16* | **C** | | **N** | **51** | **22** | **29** |
| | | **159** | | 0.196 | 0.159 | 0.224 |
| | | **171** | | 0.794 | 0.818 | 0.776 |
| | | **<u>173</u>** | | 0.010 | <u>0.023</u> | 0.000 |
| *IDVGA55* | **C** | | **N** | **51** | **22** | **29** |
| | | **199** | | 1.000 | 1.000 | 1.000 |
| *KY1/2 (sexmarker)* | **C** | | **N** | **51** | **22** | **29** |
| | | **170** | | 0.167 | 0.205 | 0.138 |
| | | **233** | | 0.833 | 0.795 | 0.862 |

No statistical evidence for null alleles (Supplementary Table S3) or scoring errors were found (level of significance: > 5 % ($\alpha$ = 0.05)).

**Supplementary Table S3: Comparison of estimated null allele frequencies of the microsatellites of this study on European bison (*n* = 51) using four algorithms[5–7].**

| *Locus* | *van Oosterhout* | *Chakraborty* | *Brookfield 1* | *Brookfield 2* |
|---|---|---|---|---|
| *DIK082* | 0.078 | 0.0856 | 0.0515 | 0.0515 |
| *IDVGA59* | -0.0328 | -0.0307 | -0.0181 | 0.1432 |
| *BM4208* | 0.035 | 0.0389 | 0.0167 | 0.0167 |
| *BM203* | -0.0311 | -0.0155 | -0.0018 | 0 |
| *CSSM19* | 0.0175 | 0.0123 | 0.0088 | 0.0088 |
| *CSSM14* | 0.0597 | 0.0694 | 0.0428 | 0.0428 |
| *CSSM22* | -0.0103 | -0.0051 | -0.0002 | 0 |
| *BMC1009* | -0.0626 | -0.0271 | -0.0058 | 0 |
| *BM1818* | -0.0534 | -0.0494 | -0.0346 | 0 |
| *MM12* | 0.0513 | 0.0557 | 0.0342 | 0.0342 |
| *Haut14* | 0.0628 | 0.0694 | 0.0428 | 0.0428 |
| *CSSM16* | -0.0593 | -0.0462 | -0.0233 | 0 |

#### 96 SNP Panel

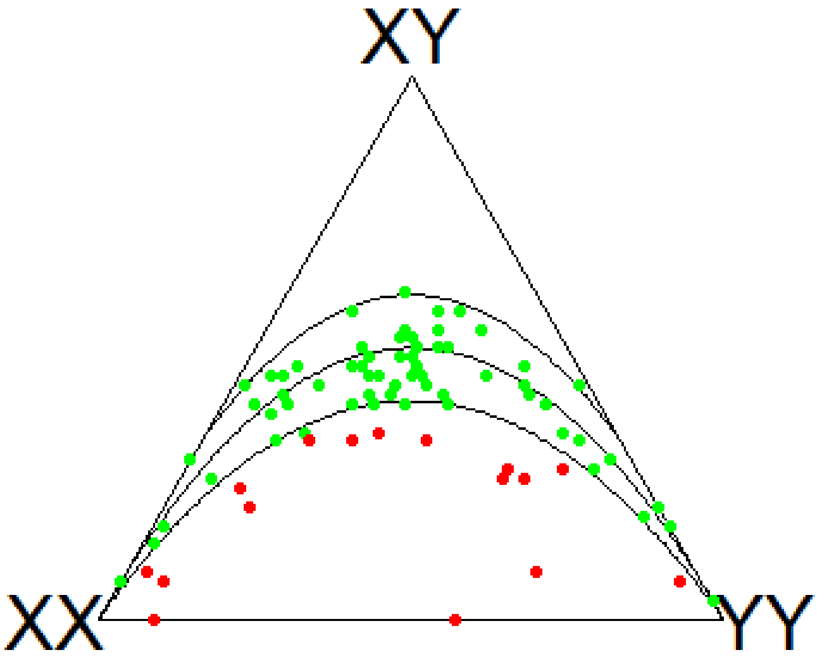

**Supplementary Figure S2: Ternary Hardy-Weinberg-Equilibrium (HWE) plots showing heterozygote deficiencies or excesses of 90 autosomal markers of 58 non-first-degree relatives of European bison.** Green dots represent the 74 loci in HWE whereas 16 loci (red dots) are deviating from HWE. The HWE parabola (intermediate curve) and acceptance region (between lower and upper curves) for the $\chi^2$ test ($\alpha = 0.05$) are shown. XX and XY symbolise both the monomorphic and XY the polymorphic states of the markers marking the genotype count vectors. Table with *p*-values per locus for HWE can be found in the supplementary file '*SNP_marker_list_details.xlsx*'.

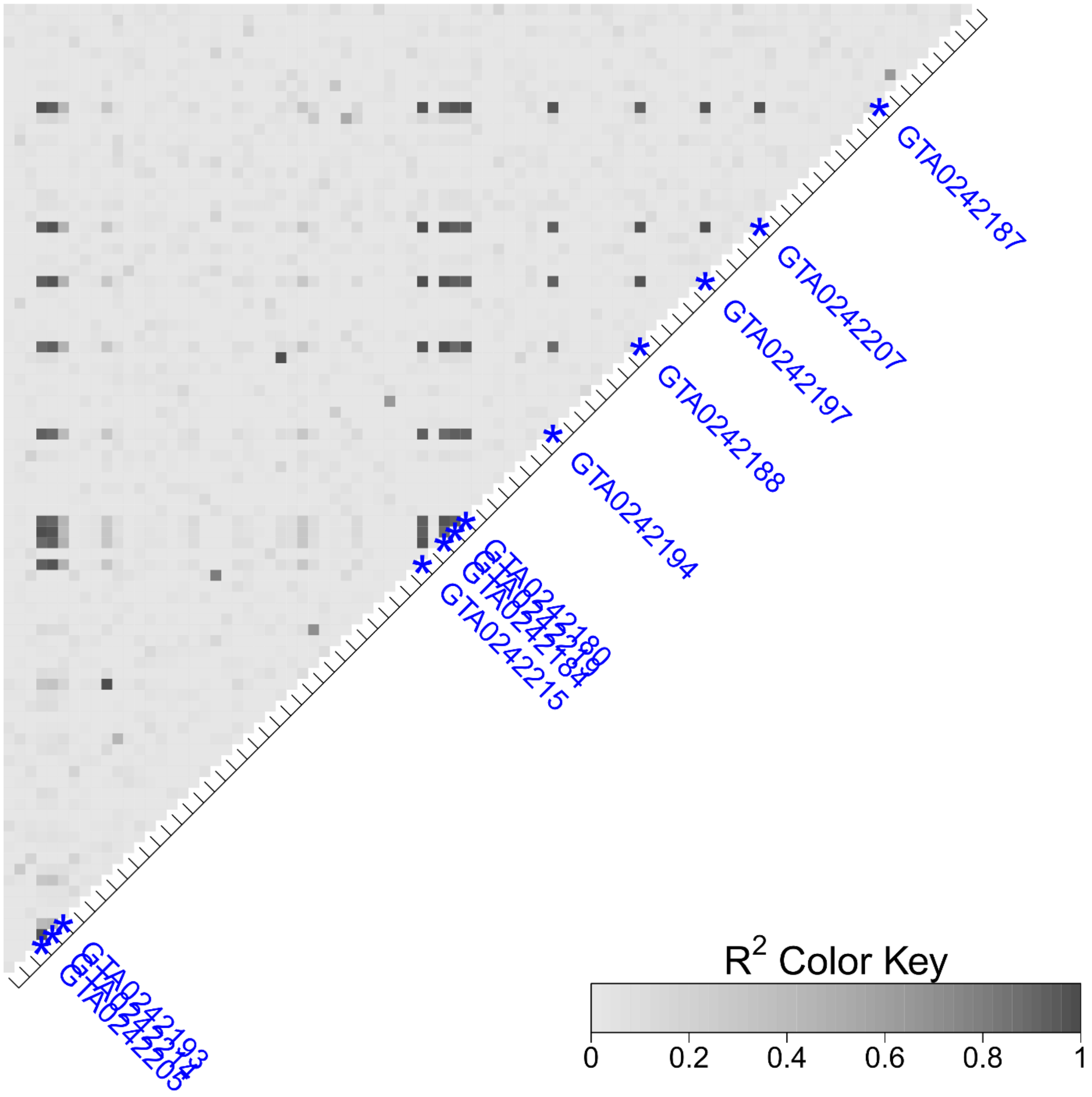

**Supplementary Figure S3: Pairwise linkage disequilibrium heatmap.** Pairwise linkage disequilibrium ($R^2$) calculated for 90 autosomal SNPs polymorphic in the European bison in 58 non-first-order relatives. Regardless of their LD, all 12 markers with an association to *posthitis*[8] are labelled in blue.

#### Individual identification

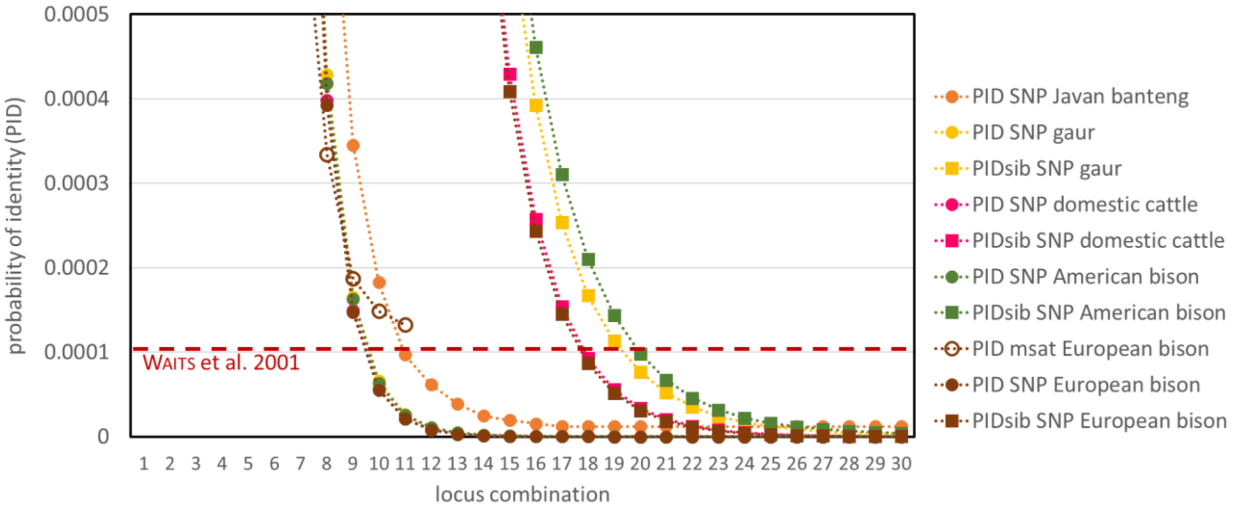

**Supplementary Figure S4:** ***Probability of identity (PID) and probability of identity among siblings (PIDsib) of genotyped microsatellites (n = 11) and SNPs (n = 95) for European bison, American bison, domestic cattle, gaur and banteng.*** Horizontal dashed red line: PID threshold for natural populations by Waits *et al.*[9] is not overcome by the microsatellite panel but is fulfilled by the SNP Panel for all shown species. The x-axis was cut at locus combination of 30 loci for more conciseness whereby the approximation of the SNP-based PIDs does not change after 30 loci. PIDsibs estimations of the microsatellite panel and the PIDsib SNP estimation for banteng are outside of the scale.

#### Assessment of genetic diversity

| | | SNP genotypes | | | | | | | pedigree | | | |
|---|---|---|---|---|---|---|---|---|---|---|---|---|
| **(sub)population/ set of individuals** | *n* | Allelic richness | $H_O^{GenAlEx}$<br>$H_O^{FSTAT}$ | $H_S^{GenAlEx}$<br>$uH_E^{GenAlEx}$<br>$H_S^{FSTAT}$ | $H_T^{GenAlEx}$<br>$H_T^{FSTAT}$ | $F_{IT}^{GenAlEx}$ | $F_{IS}^{GenAlEx1}$<br>$F_{IS}^{GenAlEx2}$<br>$F_{IS}^{FSTAT}$ | $F_{ST}^{GenAlEx1}$<br>$F_{ST}^{GenAlEx2}$<br>$F_{ST}^{FSTAT}$ | $GD^{PMx3}$<br>$GD^{PMx4}$ | $F_{IT}^{ENDOG}$ | $F_{IS}^{ENDOG}$ | $F_{ST}^{PMx}$<br>$F_{ST}^{ENDOG}$ |
| **Wisent (total)** | | | | | | | | | | | | |
| all sampled with pedigree (total) | 338<br>[1,296] | - | - | - | - | - | - | - | 0.8252<br>0.8248 | 0.0587 | 0.0219 | 0.0243<br>0.0376 |
| all genotyped | 137 | 126 | 0.400 (0.012)<br>0.400 (0.015) | 0.406 (0.011)<br>0.409 (0.011)<br>0.409 (0.014) | 0.420 (0.014)<br>0.422 (0.014) | 0.049 (0.012) | 0.017 (0.011)<br>0.015 (0.011)<br>0.024 (0.010) | 0.034 (0.005)<br>0.033 (0.005)<br>0.030 (0.005) | - | - | - | - |
| all genotyped with pedigree | 99<br>[982] | 126 | 0.400 (0.012)<br>0.400 (0.015) | 0.397 (0.011)<br>0.401 (0.011)<br>0.401 (0.014) | 0.414 (0.014)<br>0.417 (0.014) | 0.036 (0.015) | -0.006 (0.013)<br>-0.008 (0.013)<br>0.004 (0.013) | 0.043 (0.006)<br>0.043 (0.006)<br>0.037 (0.006) | 0.8034<br>0.8037 | 0.0574 | 0.0105 | 0.0546<br>0.0474 |
| **LC** | | | | | | | | | | | | |
| all sampled with pedigree (total) | 243<br>[1,032] | - | - | - | - | - | - | - | 0.8248<br>0.8209 | - | - | - |
| all genotyped | 76 | 126 | 0.403 (0.013)<br>0.406 (0.013) | 0.418 (0.012)<br>0.421 (0.012)<br>0.424 (0.012) | 0.389 (0.015)<br>0.424 (0.012) | - | - | - | - | - | - | - |
| all genotyped with pedigree | 59<br>[785] | 126 | 0.395 (0.022)<br>0.403 (0.014) | 0.360 (0.014)<br>0.413 (0.017)<br>0.419 (0.012) | 0.386 (0.016)<br>0.419 (0.012) | - | - | - | 0.8119<br>0.8074 | - | - | - |
| **LL** | | | | | | | | | | | | |
| all sampled with pedigree (total) | 95<br>[410] | - | - | - | - | - | - | - | 0.611<br>0.6041 | | - | - |
| all genotyped | 61 | 122 | 0.398 (0.019)<br>0.405 (0.018) | 0.395 (0.018)<br>0.398 (0.018)<br>0.405 (0.017) | 0.359 (0.019)<br>0.405 (0.017) | - | - | - | - | - | - | - |
| all genotyped with pedigree | 40<br>[340] | 122 | 0.432 (0.029)<br>0.406 (0.019) | 0.355 (0.019)<br>0.413 (0.018)<br>0.392 (0.017) | 0.379 (0.020)<br>0.392 (0.017) | - | - | - | 0.5673<br>0.5625 | - | - | - |

**Supplementary Table S4: Genetic diversity measures based on SNP genotypes and pedigree data for different sets of European bison individuals.** The molecular values are based on 63 SNP loci in HWE. For all 277 sampled individuals with known genealogy (total population) it was possible to generate pedigree-based genetic values (based on 338 individuals). Genealogical information was not available for all successfully genotyped individuals, whereby a complete pedigree-based assessment is not possible. Thus, molecular and pedigree-based genetic diversity values were calculated for an overlapping set of 99 successfully SNP-genotyped individuals with available genealogical data. Sample sizes [*n*] in squared brackets show the number of individuals included in the associated pedigree. Values in brackets below the genetic values represent the associated standard error (SE). *F*-statistics in *GenAlEx* were partly calculated in two different ways: [1] arithmetic averages; [2] calculated based on the average $H_S$ and $H_T$ over loci. Mean $H_E$ and $H_S$ calculated in *GenAlEx* are homologous. Pedigree-based genetic diversity values in *PMx* were calculated utilising two methods: [3] based on kinship matrix; [4] based on gene drop.

### Breeding line discrimination

#### Breeding line discrimination based on the SNP panel

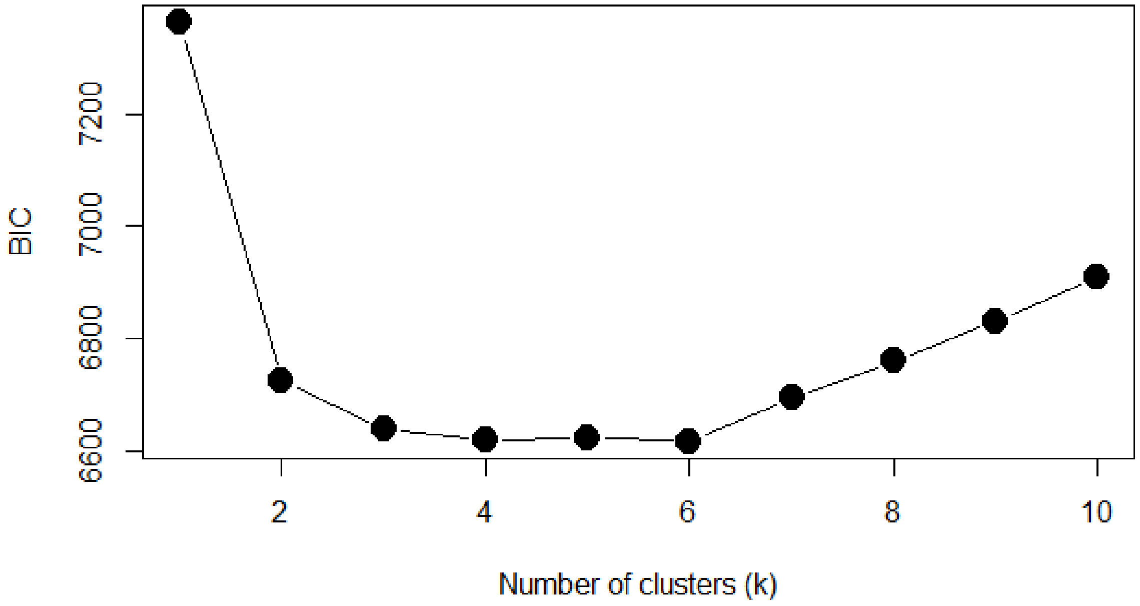

**Supplementary Figure S5: BIC for one to ten assumed *K* from maximum-likelihood genetic clustering with 18 SNP markers and 137 European bison.** The subset of 18 markers was selected to discriminate between two breeding lines in the wisent. The lower BICs for *K* = 3 to 6 might reflect genetic structures of closely related individuals.

**Supplementary Table S5: Assignment probabilities [%] based on 18 loci selected for breeding line discrimination between the LC (*n* = 76) and LL line (*n* = 61) in the European bison.** Two methods are compared: Bayesian genetic clustering computed with *STRUCTURE*; Maximum-likelihood genetic clustering computed with *adegenet*. Individuals (EBPB# and study internal names) are ordered within their breeding line (according to the metadata) after assignment probabilities computed with the Bayesian clustering (same order as in the barplot (Fig. 4)).

| Assignment probabilities [%] | | | | |
|---|---|---|---|---|
| | **Bayesian genetic clustering (*STRUCTURE*)** | | **Maximum-likelihood genetic clustering (*adegenet*)** | |
| **ID** | **LC** | **LL** | **LC** | **LL** |
| **EBPB#9912** | 0.9884 | 0.0116 | 1 | 1.85199E-12 |
| **EBPB#12963** | 0.987 | 0.013 | 1 | 6.83411E-11 |
| **BB_WILD2** | 0.9866 | 0.0134 | 1 | 2.59997E-09 |
| **EBPB#13088** | 0.9859 | 0.0141 | 1 | 9.64312E-10 |
| **EBPB#13177** | 0.9851 | 0.0149 | 1 | 9.58942E-10 |
| **EBPB#12657** | 0.985 | 0.015 | 1 | 1.02125E-10 |
| **EBPB#13636** | 0.9849 | 0.0151 | 1 | 8.8023E-10 |
| **EBPB#13761** | 0.9844 | 0.0156 | 0.9999998 | 1.98083E-07 |
| **EBPB#11843** | 0.9824 | 0.0176 | 1 | 2.56769E-11 |
| **EBPB#9952** | 0.9816 | 0.0184 | 0.9999999 | 1.12899E-07 |
| **EBPB#13700** | 0.9814 | 0.0186 | 1 | 4.13591E-08 |
| **1100_LC** | 0.9811 | 0.0189 | 0.9999998 | 1.83973E-07 |
| **EBPB#14139** | 0.9806 | 0.0194 | 1 | 4.42529E-08 |
| **EBPB#12102** | 0.9799 | 0.0201 | 1 | 2.79501E-12 |
| **EBPB#13321** | 0.9798 | 0.0202 | 1 | 2.29168E-08 |
| **EBPB#13634** | 0.9792 | 0.0208 | 0.9999917 | 8.27643E-06 |
| **EBPB#9318** | 0.9791 | 0.0209 | 1 | 1.19423E-09 |
| **EBPB#9186** | 0.9773 | 0.0227 | 0.9999996 | 3.89099E-07 |
| **EBPB#11517** | 0.9762 | 0.0238 | 0.9999996 | 3.5699E-07 |
| **EBPB#13637** | 0.9749 | 0.0251 | 1 | 7.25235E-09 |
| **EBPB#13659** | 0.973 | 0.027 | 0.9999994 | 5.50732E-07 |
| **EBPB#13407** | 0.9716 | 0.0284 | 0.9999999 | 6.99615E-08 |
| **EBPB#13323** | 0.9673 | 0.0327 | 0.9999956 | 4.39437E-06 |
| **1103_LC** | 0.9669 | 0.0331 | 0.9999862 | 1.37735E-05 |
| **EBPB#11939** | 0.9623 | 0.0377 | 0.999998 | 1.9855E-06 |
| **EBPB#11042** | 0.9618 | 0.0382 | 0.9999964 | 3.59094E-06 |
| **EBPB#10211** | 0.9607 | 0.0393 | 0.9999874 | 1.26271E-05 |
| **EBPB#11853** | 0.9605 | 0.0395 | 0.9999943 | 5.72498E-06 |
| **EBPB#11336** | 0.96 | 0.04 | 0.9999997 | 3.00423E-07 |
| **KUH_LC_1** | 0.9593 | 0.0407 | 0.9991749 | 0.000825083 |
| **EBPB#12629** | 0.9589 | 0.0411 | 0.9999936 | 6.41182E-06 |
| **EBPB#12045** | 0.9585 | 0.0415 | 0.9999204 | 7.96432E-05 |
| **EBPB#9707** | 0.9575 | 0.0425 | 0.9999838 | 1.61768E-05 |
| **EBPB#13358** | 0.9565 | 0.0435 | 1 | 1.5896E-08 |
| **EBPB#13826** | 0.9565 | 0.0435 | 0.999999 | 1.03027E-06 |
| **EBPB#12589** | 0.9559 | 0.0441 | 0.99999 | 1.00004E-05 |
| **EBPB#12223** | 0.9512 | 0.0488 | 0.9999978 | 2.21915E-06 |
| **EBPB#9940** | 0.9511 | 0.0489 | 0.999985 | 1.49954E-05 |

| | | | | |
|---|---|---|---|---|
| **EBPB#10152** | 0.948 | 0.052 | 0.9999554 | 4.4621E-05 |
| **1108_LC** | 0.9456 | 0.0544 | 0.999976 | 2.40096E-05 |
| **1111_LC** | 0.9446 | 0.0554 | 0.999981 | 1.90085E-05 |
| **EBPB#13966** | 0.9429 | 0.0571 | 0.9999571 | 4.29221E-05 |
| **1104_LC** | 0.9411 | 0.0589 | 0.9998706 | 0.000129416 |
| **EBPB#9823** | 0.941 | 0.059 | 0.9997972 | 0.000202758 |
| **EBPB#13082** | 0.9379 | 0.0621 | 0.9995719 | 0.000428082 |
| **EBPB#13517** | 0.9341 | 0.0659 | 0.9999772 | 2.28467E-05 |
| **KUH_LC_3** | 0.9297 | 0.0703 | 0.9999998 | 2.12087E-07 |
| **EBPB#12269** | 0.9285 | 0.0715 | 0.9999993 | 6.71157E-07 |
| **EBPB#13273** | 0.9236 | 0.0764 | 0.9999967 | 3.27532E-06 |
| **EBPB#11820** | 0.9235 | 0.0765 | 0.9999989 | 1.10022E-06 |
| **EBPB#13871** | 0.9219 | 0.0781 | 0.9999995 | 4.8839E-07 |
| **KUH_LC_4** | 0.9143 | 0.0857 | 0.9998719 | 0.000128057 |
| **EBPB#11338** | 0.9119 | 0.0881 | 0.9999052 | 9.48317E-05 |
| **EBPB#14062** | 0.9112 | 0.0888 | 0.9998983 | 0.00010174 |
| **EBPB#12097** | 0.9076 | 0.0924 | 0.9999998 | 1.53064E-07 |
| **BB_WILDA** | 0.8987 | 0.1013 | 0.9999606 | 3.94415E-05 |
| **EBPB#12801** | 0.8938 | 0.1062 | 0.9999977 | 2.28715E-06 |
| **1106_LC** | 0.8832 | 0.1168 | 0.9996025 | 0.000397456 |
| **EBPB#12415** | 0.8779 | 0.1221 | 0.9988138 | 0.00118624 |
| **KUH_LC_2** | 0.8625 | 0.1375 | 0.9993132 | 0.000686813 |
| **1102_LC** | 0.8592 | 0.1408 | 0.9997782 | 0.000221779 |
| **EBPB#9501** | 0.8569 | 0.1431 | 0.9996476 | 0.000352361 |
| **EBPB#9934** | 0.8394 | 0.1606 | 0.9978022 | 0.002197802 |
| **EBPB#9054** | 0.8363 | 0.1637 | 0.9705882 | 0.02941176 |
| **EBPB#13870** | 0.83 | 0.17 | 0.9998477 | 0.000152253 |
| **EBPB#9291** | 0.8257 | 0.1743 | 0.9998932 | 0.000106803 |
| **EBPB#13633** | 0.8243 | 0.1757 | 0.9998735 | 0.000126502 |
| **EBPB#11933** | 0.7626 | 0.2374 | 0.9974874 | 0.002512563 |
| **EBPB#11427** | 0.6679 | 0.3321 | 0.9333333 | 0.06666667 |
| **EBPB#11295** | 0.6505 | 0.3495 | 0.9 | 0.1 |
| **EBPB#8652** | 0.6076 | 0.3924 | 0.9583333 | 0.04166667 |
| **EBPB#10994** | 0.5008 | 0.4992 | 0.75 | 0.25 |
| **1107_LC** | 0.3368 | 0.6632 | 0.07692308 | 0.9230769 |
| **1109_LC** | 0.0339 | 0.9661 | 3.53782E-05 | 0.9999646 |
| **1105_LC** | 0.0306 | 0.9694 | 7.14321E-06 | 0.9999929 |
| **1113_LC** | 0.0185 | 0.9815 | 6.37961E-07 | 0.9999994 |
| **74** | 0.6587 | 0.3413 | 0.5 | 0.5 |
| **1301** | 0.5085 | 0.4915 | 0.25 | 0.75 |
| **EBPB#9964** | 0.3778 | 0.6222 | 0.1666667 | 0.8333333 |
| **EBPB#11943** | 0.3754 | 0.6246 | 0.125 | 0.875 |
| **EBPB#11944** | 0.3476 | 0.6524 | 0.05882353 | 0.9411765 |
| **86R** | 0.2794 | 0.7206 | 0.09090909 | 0.9090909 |
| **38K** | 0.2412 | 0.7588 | 0.02222222 | 0.9777778 |
| **73** | 0.2321 | 0.7679 | 0.02040816 | 0.9795918 |
| **EBPB#13913** | 0.2192 | 0.7808 | 0.02040816 | 0.9795918 |

| | | | | |
|---|---|---|---|---|
| **EBPB#10199** | 0.2102 | 0.7898 | 0.000255558 | 0.9997444 |
| **EBPB#14090** | 0.1939 | 0.8061 | 0.001329787 | 0.9986702 |
| **EBPB#10730** | 0.1876 | 0.8124 | 3.87522E-05 | 0.9999612 |
| **EBPB#12644** | 0.1705 | 0.8295 | 0.000898473 | 0.9991015 |
| **EBPB#12809** | 0.1461 | 0.8539 | 0.001338688 | 0.9986613 |
| **1305** | 0.1393 | 0.8607 | 0.001148106 | 0.9988519 |
| **1316** | 0.1116 | 0.8884 | 0.000102417 | 0.9998976 |
| **915** | 0.1083 | 0.8917 | 0.005235602 | 0.9947644 |
| **1266** | 0.1077 | 0.8923 | 0.001132503 | 0.9988675 |
| **1312** | 0.1072 | 0.8928 | 0.000387747 | 0.9996123 |
| **JUNGBULLE_LL** | 0.1058 | 0.8942 | 0.006756757 | 0.9932432 |
| **EBPB#9763** | 0.0959 | 0.9041 | 8.73591E-05 | 0.9999126 |
| **161R** | 0.095 | 0.905 | 0.003571429 | 0.9964286 |
| **1311** | 0.0753 | 0.9247 | 2.72546E-05 | 0.9999727 |
| **EBPB#10380** | 0.0658 | 0.9342 | 0.000236911 | 0.9997631 |
| **EBPB#11991** | 0.0643 | 0.9357 | 4.03177E-05 | 0.9999597 |
| **EBPB#14175** | 0.061 | 0.939 | 0.000483559 | 0.9995164 |
| **EBPB#14137** | 0.0548 | 0.9452 | 6.17742E-05 | 0.9999382 |
| **EBPB#13764** | 0.0544 | 0.9456 | 0.000267594 | 0.9997324 |
| **EBPB#12319** | 0.0498 | 0.9502 | 3.80154E-06 | 0.9999962 |
| **EBPB#9901** | 0.0392 | 0.9608 | 7.12048E-05 | 0.9999288 |
| **EBPB#13868** | 0.035 | 0.965 | 0.000316256 | 0.9996837 |
| **162R** | 0.0334 | 0.9666 | 3.10434E-05 | 0.999969 |
| **EBPB#10233** | 0.033 | 0.967 | 2.97708E-05 | 0.9999702 |
| **EBPB#12017** | 0.0321 | 0.9679 | 1.91157E-05 | 0.9999809 |
| **1300** | 0.0319 | 0.9681 | 1.93608E-06 | 0.9999981 |
| **EBPB#11159** | 0.0311 | 0.9689 | 1.74328E-05 | 0.9999826 |
| **925** | 0.0297 | 0.9703 | 1.62091E-06 | 0.9999984 |
| **EBPB#13293** | 0.0279 | 0.9721 | 2.26954E-07 | 0.9999998 |
| **EBPB#10448** | 0.026 | 0.974 | 7.23019E-06 | 0.9999928 |
| **EBPB#13849** | 0.0248 | 0.9752 | 5.34245E-06 | 0.9999947 |
| **KUH_LL_1** | 0.0221 | 0.9779 | 2.05724E-06 | 0.9999979 |
| **GOZUBR** | 0.022 | 0.978 | 3.15438E-06 | 0.9999968 |
| **EBPB#13640** | 0.0219 | 0.9781 | 1.31424E-06 | 0.9999987 |
| **EBPB#13954** | 0.0216 | 0.9784 | 2.17476E-07 | 0.9999998 |
| **KUH_LL_2** | 0.0213 | 0.9787 | 2.08553E-06 | 0.9999979 |
| **1306** | 0.0205 | 0.9795 | 1.78931E-06 | 0.9999982 |
| **EBPB#12317** | 0.0198 | 0.9802 | 2.23334E-07 | 0.9999998 |
| **BULLE** | 0.0184 | 0.9816 | 2.90653E-07 | 0.9999997 |
| **EBPB#10979** | 0.018 | 0.982 | 5.1014E-07 | 0.9999995 |
| **EBPB#9434** | 0.0169 | 0.9831 | 6.65341E-08 | 0.9999999 |
| **EBPB#14173** | 0.0165 | 0.9835 | 1.28488E-07 | 0.9999999 |
| **EBPB#11256** | 0.0163 | 0.9837 | 1.46793E-08 | 1 |
| **EBPB#10445** | 0.0159 | 0.9841 | 2.16867E-07 | 0.9999998 |
| **EBPB#11872** | 0.0158 | 0.9842 | 9.80688E-08 | 0.9999999 |
| **EBPB#14160** | 0.015 | 0.985 | 1.63525E-07 | 0.9999998 |
| **EBPB#12777** | 0.0127 | 0.9873 | 2.10459E-08 | 1 |

| | | | | |
|---|---|---|---|---|
| **EBPB#11951** | 0.0125 | 0.9875 | 2.14699E-09 | 1 |
| **EBPB#12776** | 0.0124 | 0.9876 | 3.25038E-09 | 1 |
| **EBPB#13955** | 0.012 | 0.988 | 1.0814E-08 | 1 |
| **EBPB#12703** | 0.0119 | 0.9881 | 1.14043E-08 | 1 |
| **EBPB#14174** | 0.0112 | 0.9888 | 4.63031E-09 | 1 |

#### Breeding line discrimination based on microsatellites

Beside the non-functional microsatellite markers all monomorphic and sex microsatellite markers (Supplementary Table S2) were excluded for further evaluation of genetic population structure. This results in a set of 12 heterozygous microsatellites markers.

We tested 51 European bison of which 22 individuals were assigned to LL and 29 individuals to LC. With the selected set of twelve autosomal microsatellites no discrimination of subpopulations or breeding lines were achievable (Supplementary Figure S6).

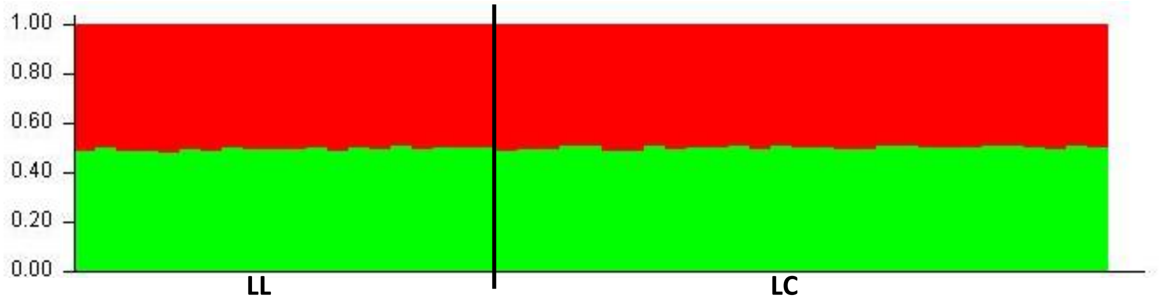

**Supplementary Figure S6:** ***STRUCTURE*** **barplot based on 12 microsatellite markers (Supplementary Table S2; not including the homozygous and sex markers) genotyping of 51 individuals of European bison including both breeding lines (*K* = 2).**

#### Pilot Study: best practice for faecal sampling, preservation and DNA extraction

##### *Faecal sampling and sample storage methodology*

Both utilised DNA extraction kits were developed for human faeces. However, the QIAgen DNA stool mini kit was already successfully used for DNA extraction from taurine frozen faeces and faecal swab samples for microbial investigation[10–12]. Additionally, we can verify the applicability for both extraction kits for all sampled species of Bovini.

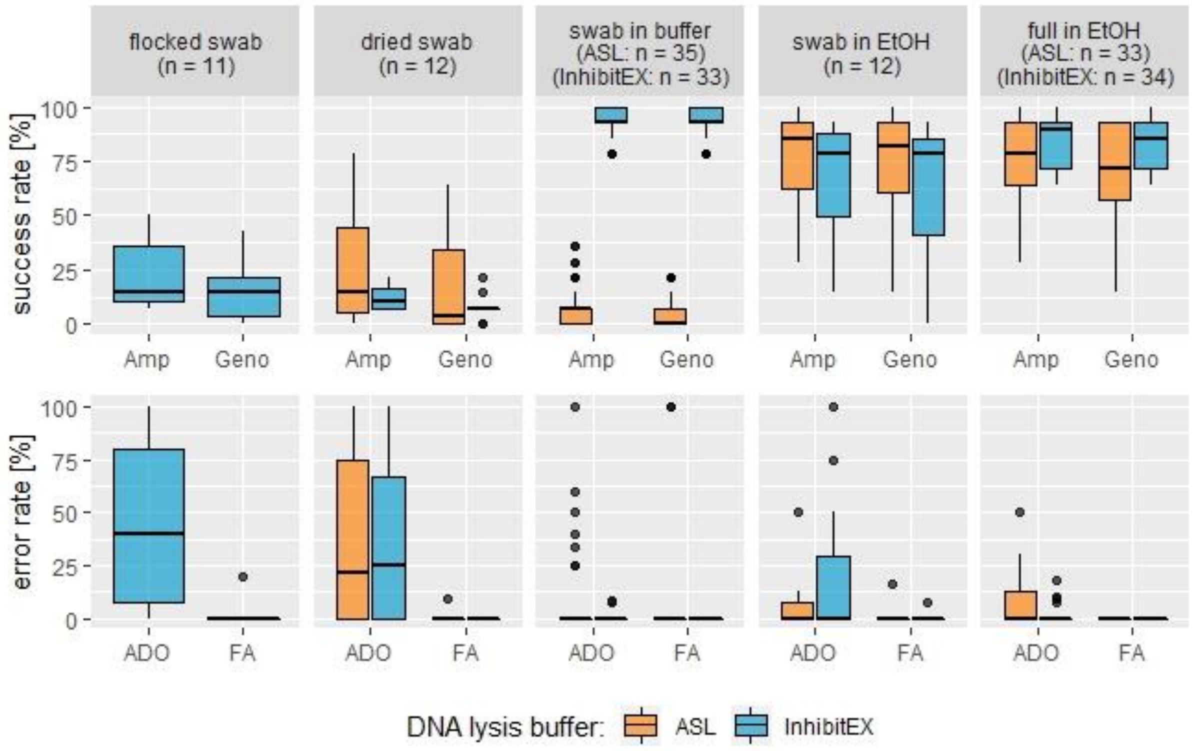

**Supplementary Figure S7: Success rates and genotyping error rates of triplicated genotypes from faecal samples (*n* = 194) collected with five sampling methods and two DNA lysis buffer.** Sample sizes per sampling method and DNA lysis buffer can be found above the boxplots and represent triplets of in total 68 DNA extractions. Those genotypes originate from two female individuals ('Falka' EBPB#9318: *n* = 94; 'Abia' EBPB#13637: *n* = 100). Amplification success rate (Amp): successful scored loci over total number of loci (*n* = 14). Genotyping success (Geno): genotyping errors (allelic dropout and false alleles) deducted from amplification success over the total number of loci. Allelic dropout rate (ADO): loci with not amplified alleles based on the consensus genotype over the number of successful amplified and scored loci (= amplification success). False allele rate (FA): amplified artefacts scored as allele based on the consensus genotype over the number of successful amplified and scored loci.

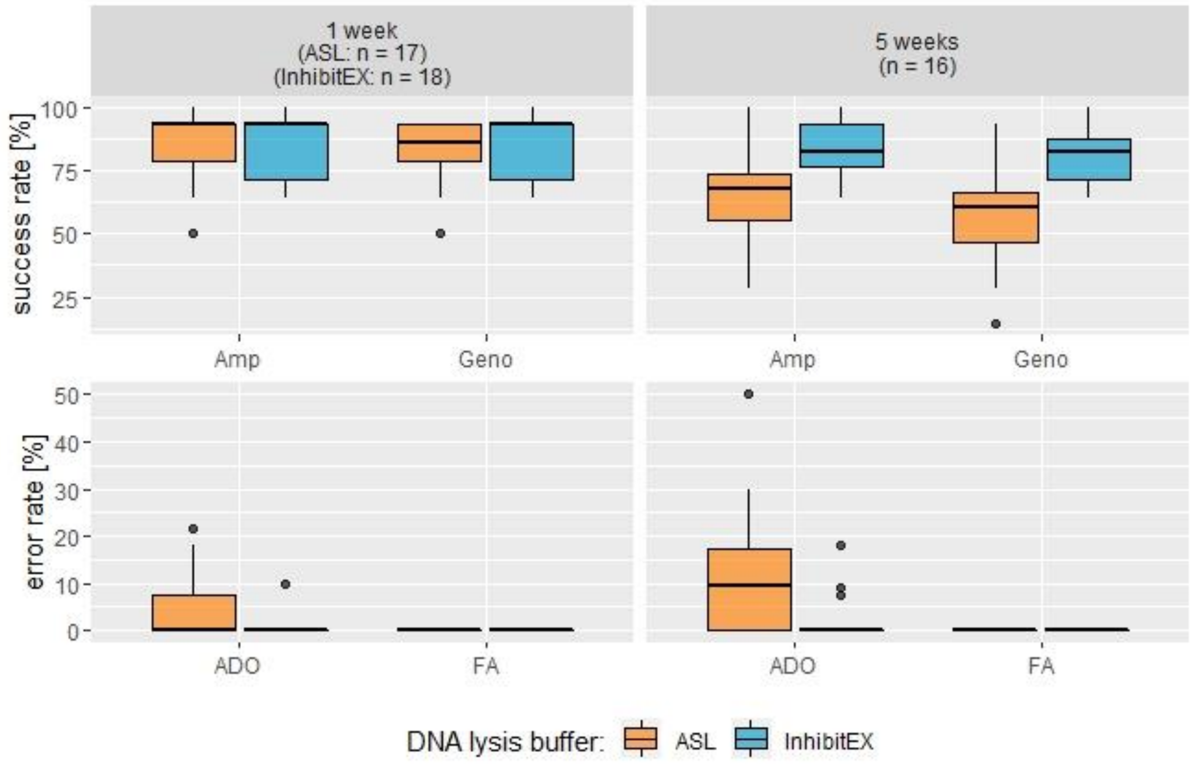

**Supplementary Figure S8: Success and genotyping error rates of triplicated genotypes from full faecal samples (*n* = 67) extracted after one and five weeks using two DNA lysis buffers corresponding to two DNA extraction kits.**

Sample sizes per storage duration and DNA lysis buffer can be found above the boxplots and represent triplets of in total 24 DNA extractions. Those genotypes originate from two female individuals ('Falka' EBPB#9318: *n* = 32; 'Abia' EBPB#13637: *n* = 35). Amplification success rate (Amp): successful scored loci over total number of loci (*n* = 14). Genotyping success (Geno): genotyping errors (allelic dropout and false alleles) deducted from amplification success over the total number of loci. Allelic dropout rate (ADO): loci with not amplified alleles based on the consensus genotype over the number of successful amplified and scored loci (= amplification success). False allele rate (FA): amplified artefacts scored as allele based on the consensus genotype over the number of successful amplified and scored loci.

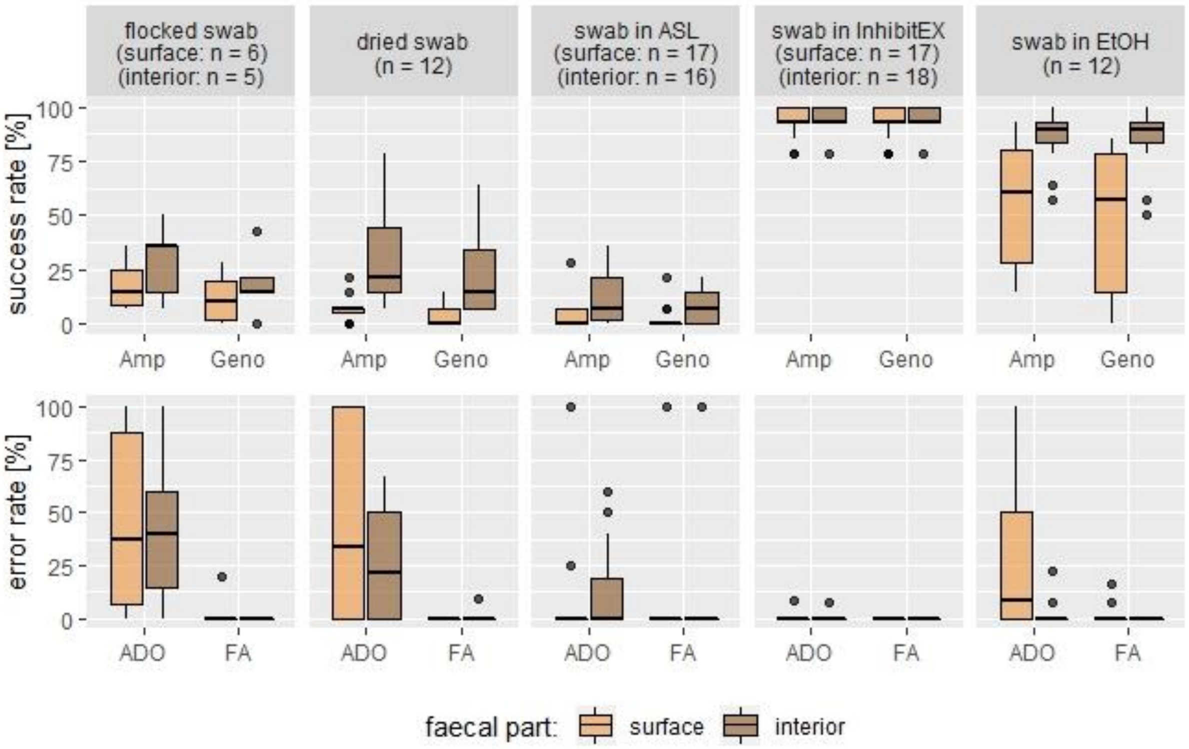

**Supplementary Figure S9: Success and genotyping error rates of triplicated genotypes from faecal samples (*n* = 127) collected with five sampling methods and two part of the wisent pat to evaluate faecal sampling methodology.** Sample sizes per sampling method and DNA lysis buffer can be found above the boxplots and represent triplets of in total 43 DNA extractions. Those genotypes originate from two female individuals (‘Falka’ EBPB#9318: *n* = 62; ‘Abia’ EBPB#13637: *n* = 65). Amplification success rate (Amp): successful scored loci over total number of loci (*n* = 14). Genotyping success (Geno): genotyping errors (allelic dropout and false alleles) deducted from amplification success over the total number of loci. Allelic dropout rate (ADO): loci with not amplified alleles based on the consensus genotype over the number of successful amplified and scored loci (= amplification success). False allele rate (FA): amplified artefacts scored as allele based on the consensus genotype over the number of successful amplified and scored loci.

**Supplementary Table S1: AICcs for the GLMMs of the success rates with/without the additional random effect 'QIAcube run', interaction terms and the null model.** Models in bold were selected. For the selected GLMMs the *p*-value on normality of the residuals (executed with the Shapiro-Wilk-test) are shown as well.

| Model | AICc | Shapiro-Wild-test (*p*-value) |
|---|---|---|
| <u>Amplification success rate</u> | | |
| **Sampling method + DNA extraction kit (Supplementary Equation S1)** | **652.7255** | **0.2055** |
| + random effect group (run) | 654.9254 | |
| Interaction terms | 655.1201 | |
| Null model | 740.2407 | |
| **Storage duration + DNA extraction kit (Supplementary Equation S2)** | **238.9371** | **0.005541** |
| + random effect group (run) | 241.2755 | |
| Interaction terms | 240.4805 | |
| Null model | 239.967 | |
| **Faecal part + sampling method + DNA extraction kit (Supplementary Equation S3)** | **404.5755** | **0.04195** |
| + random effect group (run) | 406.8936 | |
| Interaction terms | 410.2526 | |
| Null model | 474.2571 | |
| <u>Genotyping success rate</u> | | |
| **Sampling method + DNA extraction kit (Supplementary Equation S1)** | **635.1107** | **0.2134** |
| + random effect group (run) | 637.3107 | |
| Interaction terms | 636.4667 | |
| Null model | 720.6068 | |
| **Storage duration + DNA extraction kit (Supplementary Equation S2)** | **247.3894** | **0.5164** |
| + random effect group (run) | 249.7279 | |
| Interaction terms | 248.9009 | |
| Null model | 251.5641 | |
| **Faecal part + sampling method + DNA extraction kit (Supplementary Equation S3)** | **374.3496** | **0.02461** |
| + random effect group (run) | 376.6679 | |
| Interaction terms | 440.3392 | |
| Null model | 1726.25 | |

**Supplementary Table S2: AICcs for the GLMMs of the genotyping error rates with/without the additional random effect 'QIAcube run', interaction terms and the null model. Models in bold were selected.** The relationship of the storage duration and the DNA extraction kit on false alleles was not possible to determine because no false allele was detected in this experimental setup (grey). For the selected GLMMs the *p*-values on normality of the residuals (executed with the Shapiro-Wilk-test) are shown as well. No *p*-values concerning the false allele rates are given, because no GLMM was executed.

| Model | AICc | Shapiro-Wilk-test (*p*-value) |
|---|---|---|
| Allelic dropout rate | | |
| **Sampling method + DNA extraction kit (Supplementary Equation S1)** | **289.6554** | **3.789e-08** |
| + random effect group (run) | 291.8696 | |
| Interaction terms | 288.3367 | |
| Null model | 315.5074 | |
| **Storage duration + DNA extraction kit (Supplementary Equation S2)** | **106.6175** | **9.049e-05** |
| + random effect group (run) | 108.956 | |
| Interaction terms | 108.947 | |
| Null model | 115.7784 | |
| **Faecal part + sampling method + DNA extraction kit (Supplementary Equation S3)** | **177.7566** | **7.098e-07** |
| + random effect group (run) | 180.0747 | |
| Interaction terms | 187.4496 | |
| Null model | 199.4994 | |
| False alleles rate | | |
| Sampling method + DNA extraction kit (Supplementary Equation S1) | 58.31521 | |
| + random effect group (run) | 60.5151 | |
| Interaction terms | 62.04845 | |
| **Null model** | **54.26459** | |
| Storage duration + DNA extraction kit (Supplementary Equation S2) | NA | |
| + random effect group (run) | NA | |
| Interaction terms | NA | |
| Null model | NA | |
| **Faecal part + sampling method + DNA extraction kit (Supplementary Equation S3)** | **54.86338** | |
| + random effect group (run) | 57.18151 | |
| Interaction terms | 68.69846 | |
| Null model | 58.24566 | |

Effects of single samples or QIAcube runs were considered as random effects in the GLMMs. No explanatory improvements for the models were shown with the QIAcube runs as random effect groups and were subsequently excluded. Therefore, this variable is neglected in the following evaluation of best practice for the faecal sampling method.

**Supplementary Table S3: Summary of intercepts, standard errors, z-values and the *p*-value (Pr(>|z|)) for the predictor factors in the GLMM with the response variable 'amplification success rate'.** For each model sample sizes (*n*) are attached. The sample size (*n*) for the random effect groups represents the physical samples (divided by the triplicates used in the model). Significance codes: not significant 'ns', < 0.1 '.', < 0.05 '*', < 0.01 '**', < 0.001 '***'.

| predictor | intercept | Standard error | z-value | Pr(>\|z\|) | |
|---|---|---|---|---|---|
| **Sampling method + DNA extraction kit (Supplementary Equation S1)** | *n* = 194 | | | | |
| | -1.35625 | 0.68447 | -1.981 | 0.047538 | * |
| sampling_methodfull_EtOH | 2.97495 | 0.65128 | 4.568 | 4.93e-06 | *** |
| sampling_methodswab_buffer_ASL | -1.61580 | 0.78466 | -2.059 | 0.039471 | * |
| sampling_methodswab_buffer_InhibitEX | 4.50090 | 0.70075 | 6.423 | 1.34e-10 | *** |
| sampling_methodswab_dry | -0.46640 | 0.73464 | -0.635 | 0.525517 | ns |
| sampling_methodswab_EtOH | 2.49529 | 0.72857 | 3.425 | 0.000615 | *** |
| DNA_extraction_kitInhibitEX | 0.06884 | 0.37243 | 0.185 | 0.853345 | ns |
| Random effect groups (lab#): *n* = 68 | | | | | |
| **Storage duration + DNA extraction kit (Supplementary Equation S2)** | *n* = 67 | | | | |
| | 1.8247 | 0.3579 | 5.098 | 3.43e-07 | *** |
| storage_time5 | -0.8722 | 0.4082 | -2.137 | 0.0326 | * |
| DNA_extraction_kitInhibitEX | 0.4922 | 0.4071 | 1.209 | 0.2267 | ns |
| Random effect groups (lab#): *n* = 24 | | | | | |
| **Faecal part + sampling method + DNA extraction kit (Supplementary Equation S3)** | *n* = 127 | | | | |
| | -0.1002 | 0.7113 | -0.141 | 0.888016 | ns |
| faecal_partsurface | -1.2139 | 0.3281 | -3.700 | 0.000216 | *** |
| sampling_methodswab_buffer_ASL | -2.2727 | 0.7713 | -2.947 | 0.003212 | ** |
| sampling_methodswab_buffer_InhibitEX | 4.4421 | 0.5987 | 7.420 | 1.17e-13 | *** |
| sampling_methodswab_dry | -0.7923 | 0.6502 | -1.219 | 0.223017 | ns |
| sampling_methodswab_EtOH | 2.1832 | 0.6427 | 3.397 | 0.000682 | *** |
| DNA_extraction_kitInhibitEX | -0.5841 | 0.4971 | -1.175 | 0.239983 | ns |
| Random effect groups (lab#): *n* = 44 | | | | | |

**Supplementary Table S4: Summary of intercepts, standard errors, z-values and the *p*-value (Pr(>|z|)) for the predictor factors in the GLMM with the response variable 'genotyping success rate'.** For each model sample sizes (*n*) are attached. The sample size (*n*) for the random effect groups represents the physical samples (divided by the triplicates used in the model). Significance codes: not significant 'ns', < 0.1 '.', < 0.05 '*', < 0.01 '**', < 0.001 '***'.

| predictor | intercept | Standard error | z-value | Pr(>\|z\|) | |
|---|---|---|---|---|---|
| **Sampling method + DNA extraction kit (Supplementary Equation S1)** | *n* = 194 | | | | |
| | -2.3221 | 0.7780 | -2.985 | 0.002838 | ** |
| sampling_methodfull_EtOH | 3.6359 | 0.7415 | 4.904 | 9.41e-07 | *** |
| sampling_methodswab_buffer_ASL | -1.0694 | 0.8900 | -1.202 | 0.229519 | ns |
| sampling_methodswab_buffer_InhibitEX | 5.2927 | 0.7913 | 6.688 | 2.26e-11 | *** |
| sampling_methodswab_dry | -0.2897 | 0.8435 | -0.343 | 0.731253 | ns |
| sampling_methodswab_EtOH | 3.0608 | 0.8248 | 3.711 | 0.000206 | *** |
| DNA_extraction_kitInhibitEX | 0.2426 | 0.4118 | 0.589 | 0.555739 | ns |
| Random effect groups (lab#): *n* = 68 | | | | | |
| **Storage duration + DNA extraction kit (Supplementary Equation S2)** | *n* = 67 | | | | |
| | 1.75959 | 0.41008 | 4.291 | 1.78e-05 | *** |
| storage_time5 | -0.24583 | 0.09885 | -2.487 | 0.0129 | * |
| DNA_extraction_kitInhibitEX | 0.77095 | 0.39431 | 1.955 | 0.0506 | . |
| Random effect groups (lab#): *n* = 24 | | | | | |
| **Faecal part + sampling method + DNA extraction kit (Supplementary Equation S3)** | *n* = 127 | | | | |
| | -0.7504 | 0.8464 | -0.887 | 0.375330 | |
| faecal_partsurface | -1.4529 | 0.3867 | -3.757 | 0.000172 | *** |
| sampling_methodswab_buffer_ASL | -1.9827 | 0.9185 | -2.159 | 0.030876 | * |
| sampling_methodswab_buffer_InhibitEX | 5.2674 | 0.7075 | 7.445 | 9.72e-14 | *** |
| sampling_methodswab_dry | -0.7389 | 0.7854 | -0.941 | 0.346871 | ns |
| sampling_methodswab_EtOH | 2.6680 | 0.7661 | 3.483 | 0.000497 | *** |
| DNA_extraction_kitInhibitEX | -0.6320 | 0.5875 | -1.076 | 0.282042 | ns |
| Random effect groups (lab#): *n* = 44 | | | | | |

**Supplementary Table S5: Summary of intercepts, standard errors, z-values and the *p*-value (Pr(>|z|)) for the predictor factors in the GLMM with the response variable 'allelic dropout rate'.** For each model sample sizes (*n*) are attached. The sample size (*n*) for the random effect groups represents the physical samples (divided by the triplicates used in the model). Significance codes: not significant 'ns', < 0.1 '.', < 0.05 '*', < 0.01 '**', < 0.001 '***'.

| predictor | intercept | Standard error | z-value | Pr(>\|z\|) | |
|---|---|---|---|---|---|
| **Sampling method + DNA extraction kit (Supplementary Equation S1)** | *n* = 194 | | | | |
| | 0.1994 | 1.0522 | 0.189 | 0.849711 | ns |
| sampling_methodfull_EtOH | -3.6215 | 1.0373 | -3.491 | 0.000481 | *** |
| sampling_methodswab_buffer_ASL | -2.0819 | 1.3121 | -1.587 | 0.112589 | ns |
| sampling_methodswab_buffer_InhibitEX | -5.3668 | 1.2478 | -4.301 | 1.7e-05 | *** |
| sampling_methodswab_dry | -0.2634 | 1.1227 | -0.235 | 0.814489 | ns |
| sampling_methodswab_EtOH | -2.7152 | 1.1081 | -2.450 | 0.014278 | * |
| DNA_extraction_kitInhibitEX | -1.0372 | 0.6175 | -1.680 | 0.093020 | . |
| Random effect groups (lab#): *n* = 68 | | | | | |
| **Storage duration + DNA extraction kit (Supplementary Equation S2)** | *n* = 67 | | | | |
| | -4.1505 | 0.7781 | -5.334 | 9.59e-08 | *** |
| storage_time5 | 0.4102 | 0.1801 | 2.278 | 0.02274 | * |
| DNA_extraction_kitInhibitEX | -2.1223 | 0.7342 | -2.891 | 0.00384 | ** |
| Random effect groups (lab#): *n* = 24 | | | | | |
| **Faecal part + sampling method + DNA extraction kit (Supplementary Equation S3)** | *n* = 127 | | | | |
| | -1.7393 | 1.1149 | -1.560 | 0.1187 | ns |
| faecal_partsurface | 1.1082 | 0.6068 | 1.826 | 0.0678 | . |
| sampling_methodswab_buffer_ASL | -0.2933 | 1.3000 | -0.226 | 0.8215 | ns |
| sampling_methodswab_buffer_InhibitEX | -5.2974 | 1.1475 | -4.616 | 3.9e-06 | *** |
| sampling_methodswab_dry | 0.5210 | 1.0205 | 0.511 | 0.6097 | ns |
| sampling_methodswab_EtOH | -2.0247 | 1.0130 | -1.999 | 0.0456 | * |
| DNA_extraction_kitInhibitEX | 0.4006 | 0.8029 | 0.499 | 0.6178 | ns |
| Random effect groups (lab#): *n* = 44 | | | | | |

With only 6 observed FAs (in 6 genotypes, 5 samples and 3 markers) in 1564 successful scored PCR reactions no meaningful statistical dependency testing could be implemented. In the experimental setup of the GLMM including the explanatory variable of storage duration (Supplementary Equation S2) no FA was detected at all. In two single PCR reactions exclusively, FAs were amplified and scored and therefore are only accounted in the genotyping success but not in the amplification success.

Included in every model, the two DNA extraction kits do not show a significant different impact on the success rates (Supplementary Table S3 – Supplementary Table S5). Only on the ADO rate in the GLMM including the storage duration the utilised DNA extraction kit showed significant differences (Supplementary Table S5). Especially, in DNA extractions after five weeks compared to DNA extractions after one week (Supplementary Figure S8). In contrast, sampling with a swab directly in InhibitEX buffer is significant positively different to sampling with a swab directly in ASL buffer in all models. The latter method was even significantly negatively different to every dry sampling method. All dry sampling methods were significantly negatively different to sampling in EtOH as either a swab sample, a full sample, or InhibitEX buffer. Collecting a faecal sample from a wisent pat surface was highly significant negatively different to take samples from the faecal interior in the success rates. However, it showed only a marginal significant effect on the ADO rate. The storage duration of four more weeks showed a significant negative effect on the success and ADO rates (Supplementary Figure S8; Supplementary Table S3 – Supplementary Table S5).

### Supplementary Discussion

#### Marker system: microsatellites

Ten autosomal polymorphic microsatellite marker and the applicable sex marker are utilised for the first time for European bison. Before, MM12 was successfully tested in American bison[13]. BM203 was previously determined as monomorphic in LL[14], but shows private alleles in three individuals of LC (allele frequency: 0.0306).

Previously it was shown that 17 microsatellites do not provide enough informative power to assess issues like individual identification and paternity assignment in European bison whereas SNP panel of down to 50 – 60 most heterozygous loci would be sufficient[15]. Expectably, with a new set of autosomal microsatellite markers this study can support these previous results. No sibling (Supplementary Figure S4) and breeding line discrimination (Supplementary Figure S6) was possible. The PID suggests that seven markers are sufficient to discriminate individuals but is disproven due to the fact that two individuals of LC from Russia (1100_LC and 1116_LC) showed the same microsatellite genotype. But the microsatellites used here were not preselected to address any issues for the European bison. Instead, they were implemented for the methodology evaluation regarding the optimal faecal sampling.

### Pilot Study: best practice for faecal sampling, preservation and DNA extraction

#### Sampling and sample storage methodology

Besides other studies[16–18], this pilot study provides a comprehensive discussion regarding the complications of non-invasive sampling for genetics with a focus on faecal sampling.

##### *Faecal sampling*

Utility of faecal samples for a genetic population assessment brings specific problems but provides a frequent informative DNA source for genetic wildlife monitoring and is well-established in population and conservation genetics. Due to very different success rates of DNA amplification from faeces caused by methodology, including the sampling method, the storage method and duration as well as the extraction method, but also dependent towards other factors like environmental conditions[19] and also by the biology of the studied organism such studies remain taxa-specific[20]. Pilot studies for examine the best practice are recommended[19,21].

To find the best practice for faecal sampling, preservation and DNA extraction in European bison different methods and scenarios were tested with GLMMs. Statistical research on GLMMs is still in progress and e.g. model selection is less defined as in other modelling techniques[22]. But with a set of altogether categorical predictors, not normal distributed, overdispersed data and random effects, the GLMM provided the most suitable approach. It is recommended to keep every model as simple as possible[22]. Therefore, three independent simple models were simulated and models with interaction terms were tested but reasonably rejected (Supplementary Table S3; Supplementary Table S4; Supplementary Table S5).

The microsatellite marker set, not specifically selected for European bison, was only used for the evaluation of the sampling and storage methodology since it has limited explanatory power e.g. for the species' population structures (e.g. Supplementary Figure S6). The inclusion of only two individuals in this methodology evaluation leads to certain explanatory weaknesses. Especially the presented ADO rate lacks informative power due to many homozygous loci in both individuals and the calculation based on the amplification success instead on the number of successful amplified and scored heterozygous loci[23,24]. But with the consistent calculation this error rate is still comparable and conclusive within the experiment because its trend shown in the faecal samples is consistent with former studies on other species: here too, ADO seems to be the most serious GE[25]. This goes along with our results of only six FAs along in total 1 564 amplified and scored alleles.

Maudet *et al.*[26] showed a significant seasonal dependence in GE rates from faecal samples of two caprine species. Samples were collected in spring months showed higher error rates compared with samples collected in winter maybe due to high forage quality in less harsh months. European bison also show such a seasonal diet selection due to their distribution in the European temperate zone[27,28]. The faecal consistency of European bison dung is highly variable and depends on this seasonal food composition in the wild. During spring and summer, while feeding on mostly herbaceous plants, their dung show a very loose consistency compared to winter months while feeding on shoots of trees, shrubs and hay[29]. In some holdings such seasonal foraging is present due to big enclosures with naturalistic vegetation and less supplementary feeding. The proportion of supplementary hay and concentrated feed depends on each institution's husbandry. All faecal samples of the pilot study were collected in August 2018 during environmental conditions with high temperatures and humidity not advantageous for non-invasive genetic sampling in general. Still, the present study shows successful genotyping with non-invasive samples. Since the heterogeneous husbandries as well as additional feeding with e.g. hay is a common practice even in free-roaming herds[30], the present data are applicable for wild populations regarding nutrition concerns. Likewise, Gardipee[31] sampled faecal material of wild plains bison (*B. b. bison*) in two summers with an overall high amplification success and low error rates. Hájková *et al.*[32] point out that Maudet *et al.*[26] did not mentioned the lower temperatures during winter and its potential effect on DNA degradation. High temperatures and humidity increase the activity of hydrolytic enzymes in faeces causing more rapid degradation of DNA[20,32]. Based on study results[32] it is generally recommendable to sample in colder seasons, if possible even frozen faeces. This might be unrealistic in continuous monitorings due to temporal, geographical or project-related circumstances. Based on our experience, sampling wisent dung during winter with temperatures below 0 °C could be difficult due to their sheer size. Splitting off collectable samples from frozen dung pats are potentially sources of genetic contamination, because heavy tools are required and disinfection in the field is often impractical.

Precipitation is another, probable more severe factor than temperature or humidity, negatively influencing DNA quality in faeces[33–35]. Sampling and preserving shortly after defaecation is recommended to prevent such negative impacts and improve the ultimate genetic assessment[21,33,35–38]. Previous studies showed success with sampling within 1 h up to 24 h after defaecation in captive conditions and approximately 6 h up to 21 days under field conditions[19,39–42]. So far, plains bison dung of free-roaming individuals were collected within 10 – 15 min for genotyping[31]. The obvious advantage of faeces detection in the latter case was the weald of the habitat of *B. b. bison* and, not applicable for the more forest-dwelling *B. bonasus*[43]. In a genetic nutrition study for European bison fresh faecal samples were collected after GPS-tracking of collared individuals[44], which might also not be possible for a comprehensive genetic population monitoring. Thus, it might be pivotal for e.g. monitoring programs and studies to utilise faecal samples a few days old. Yet, the genotyping success is not always predictable by the physical faecal appearance[45]. In this regard, samples collected months after defaecation showed an expectable increasing uncertainty towards genetic assignment sensitivity estimations (0.786 for a 200-day-old faecal sample)[46]. Accordingly, this holds still a certain informative power but might not be transferred one to one regarding the faecal consistency and environmental associated with European bison. With two occasionally sampled aged faeces the present pilot study we could show that genotyping is potentially possible. Regarding this, marker sets consisting of more loci than microsatellite panels, such utilised in the SNP panel, would be favourable. A single biallelic SNP locus holds less information for the genotypes but in turn would cause a lower GE if failed. Also, the SNP amplicons of the system presented here are considerably shorter than needed for microsatellites. Therefore, fragmented DNA found in faeces can be utilised more efficiently.

It was presumed that diet has an impact on DNA extraction from faeces[47–49]. But overall, genotyping success from faecal samples seems to be not heavily affect by the diet, but most likely by other parameters[50]. Exemplary, in omnivorous brown bears (*Ursus arctos*, LINNAEUS 1758) general differences in dietary ratios of carnivory, herbivory and dietary fiber itself did not affect DNA yield[51]. But, the digestion system in ruminants is radically different in contrast to a monogastric mammal[52]. In general, wisent dung with a loose consistency was very common during our sample collection. Thus, compared with more compact faeces from e.g. Canoidea (KRETZOI 1943) species wisent pats may do not strip the same amount of mucosa cells from the intestinal wall visible as light grey slough resulting in higher chances to gain higher host DNA yield from the faecal surface[19,32,35,38,51]. Such slough was not determinable on pats of European bison. Nevertheless, differences in the amplification and genotyping successes of swab samples from the faecal surface versus the faecal interior were found here. The significant negative effect of samples from the faecal surface in comparison to sampling from the wisent pat interior might be connected to its environmental exposure and therefore to outer forces accelerating DNA degradation, such as UV light. Technically, it is more difficult to take up pure surface material than faecal substance from the wisent pat interior. Therefore, samples from the interior provides more faecal material, which in turn could positively influence the genetic analysis from such a sample. Consequently, in the following sampling it was recommended to cover the cotton tip of the swab decently with faecal material, no matter from which part of the wisent pat. Uneven distributions of intestinal cells in the faecal matrix itself, not showing explicitly mucosa slough and therefore randomly chosen for DNA extraction, leads to the additional problem of non-reproducible results of amplifiable DNA yields[49,53]. This can be avoided by homogenise the faecal sample during the collection process[39,40]. The advantage of preservation of full faecal samples in 96 % EtOH is that the comparable loose matrix of wisent dung could be homogenised afterwards within the permanent storage container if necessary.

Faecal sampling with swabs is cheaper, more practical and less prone to genetic contamination[38]. But the success of gaining genotypes also depends on the type of swab sample storage. Both methods presented in the pilot study including drying faecal samples (flocked swab and dried swab samples) not only showed significant lower amplification rates but the highest ADO rates resulting in decreased genotyping success (Supplementary Figure S7). Drying faeces for subsequent DNA extraction by silica gel has been shown to be an applicable preservation and storage alternative to Drierite, freezing, freeze-drying, the usage of 70 – 100 % EtOH, preservative solutions for nucleic acids (RNA) or directly in DNA lysis buffer partly on different temperature levels[20,39,54]. Especially, for long-term storage other studies found that silica gel together with RNA*later*™ Stabilization Solution (Invitrogen™) are more effective than 95 % EtOH[55]. A two-step storage procedure combining collecting in EtOH and subsequent drying with silica gel was shown to increase DNA amplification rates significantly[20]. In contrast, the less successful results presented here with faecal samples collected and subsequently dried goes along with studies on long-term storage and silica dried faeces, which showed the lowest success in amplification rates and highest GE rates compared with preservation and storage in EtOH, DMSO/EDTA/Tris/salt solution (DETs) and by freezing[40]. The DNA extraction for all faecal swab samples in the present study were done five weeks after collection. The observation that some faecal samples moulded after a few days and subsequently may not dry fast enough to prevent DNA from further degradation were mentioned before[20,40]. Especially, in large species like the European bison the amount of collected faecal matrix is a considerable factor for the silica drying approach. Even with collecting and drying swabs with relatively small amounts of faeces in this study the latter obstacle could not be eliminated. Thus, higher amounts of non-invasive samples could increase extracted DNA quantities. But concentrations of PCR inhibiting substances would also increase particularly in faeces[21]. The faecal moisture content of taurine cattle varies between less than 80 to 90 %[56] with a comparable faecal structure to the closely related *B. bonasus*[30,57–61]. In contrast, the faecal moisture content of e.g. dogs (*Canis lupus familiaris* (LINNAEUS 1758)) also dependent on diet, body size and breed, but was measured to be mostly under 80 %[62]. Here, this relatively high moisture content in wisent dung might be a further complication regarding the storage of dried faecal samples for genetic assessments compared with other species. This also applies potentially for storing faecal samples in EtOH or other solutions regarding a certain dilution effect by the contained water. But if used in a comprehensive genetic monitoring dried faecal samples like handled in this study are not recommended due to the risk of partial informative loss of the collection. If sampling in silica is required, subsequently desiccation with another method is recommended[40].

Beside the common practice to store faecal samples in EtOH, collecting swab samples directly in DNA lysis buffer are already established[35,38,63]. In general, preservation in liquid solutions (DNA lysis buffer and 96 % EtOH) was more successful in the present pilot study. Faecal samples (full faecal and faecal swab samples) stored in 96 % EtOH showed a higher variance in success rates compared with faecal samples stored in DNA lysis buffer (Supplementary Figure S7). Consequently, storage of faecal samples in EtOH represents an overall less consistently reliable method.

Other DNA extraction kits were used for faecal samples before, for instance the Blood & Tissue kit and DNeasy Tissue Kit (Qiagen Inc. Valencia, CA) for faecal full samples[26,33]. Both DNA extraction kits utilised in the present study did not show a significant different impact on the amplification or genotyping success. Notably, storing faecal swab samples in ASL buffer shows highly significant negatively different success and ADO rates to all other storage methods in liquids. The major difference between these two DNA extraction kits is that the ingredients that remove PCR inhibiting substances present in faeces are separately added to the ASL buffer during the extraction process in the form of a tablet, while the InhibitEX buffer already contains such inhibitors removing chemistry (see Qiagen kit instructions). The ingredients of both this InhibitEX tablet (QIAamp DNA Stool Mini Kit) and the InhibitEX buffer (QIAamp Fast DNA Stool Mini Kit), are not fully provided by the manufacturer (Qiagen) but might be very similar and consequently show a similar result for the DNA extraction itself. The fact that the InhibitEX buffer already contains these ingredients, might be the explanation for the significant positive effect of sampling faeces directly in the latter buffer (Supplementary Figure S7). DNA of the faecal samples collected directly in the ASL buffer might be less protected by lacking direct inhibitor suppression. The dehydrate quality of 96 % EtOH leads to a similar positive effect in this experiment, by inhibiting enzymatic activity degrading DNA[64]. However, both faecal swab sample types (ASL/InhibitEX) were extracted in separate QIAcube runs. Thus, the possibility of an impact by handling failures during DNA extraction on the effect in the results regarding the strong differentiation cannot be ruled out. Since, searching for not only successful but practical methods the noticeably longer handling time in the extraction procedure for the samples stored in ASL buffer must be considered. Longer handling durations as an important factor regarding sampling methodology evaluations was mentioned before[38]. Collecting faecal swab samples in InhibitEX buffer showed not only the highest success rates but also low variance and is therefore the best practice. Thus, it is totally reasonable to choose the latter DNA extraction kit without further experiments. Additionally, the QIAamp DNA Stool Mini Kit with the ASL buffer will become not commercially available in the near future (Qiagen pers. comm.).

An important aspect that this pilot study could show concerns the practicability to store the samples at room temperature. This fact is of interest because it is not always possible to provide a continuous cooling chain for shipping samples.

Expectably, the storage duration had a negative effect on the success rates and positive effect on the ADO rate (Supplementary Figure S8). Here, the presented data does not represent a continuous rate of DNA degradation but enables to recommend a contemporary DNA extraction based on a significant difference in success and error rates. Notably, the amplification and genotyping success was higher after five weeks after collection from samples stored in InhibiteEX buffer than the amplification success from full samples in EtOH after only one week after collection, which represents the both best practices tested here.

Others showed that the sample collector's field experience showed an effect especially on genotyping success from nuclear DNA of faecal samples. Initial sampling training is recommended to reduce the negative effect from heterogenic skilled collectors[65]. Since the collection for this pilot study was done by only two collectors the possible error is equal and therefore neglectable. For further sampling throughout Europe simple but detailed and standardised instructions were provided for the cooperation partners.

##### *Non-faecal non-invasive sampling*

The comparatively error-prone nature of non-invasive samples regarding correct genotypes could be a reason to collect and utilise different sample types within genetic monitoring: A genotype generated by several sample types per individual reduces the potential of negative effects of single sample type. It was possible to generate a complete consensus genotype from every non-faecal sample presented here including from urine, saliva and nasal secretion.

Though, collection of frozen dung rises potential difficulties during winter months as discussed above, low temperatures can be beneficial regarding DNA degradation: sampling in snow opens the possibility to utilise urine for genetic assessment[66]. Occasional swab samples from urine-soaked snow in the our study represent a further collection opportunity without any additional preparation. Nevertheless, this sampling method relies on snow and might only be complementary in a comprehensive genetic population monitoring. Two urine swab samples directly taken from a meadow after urination in summer were also successful. Nevertheless, it was possible to generate the entire genotype from only this urine sample. But due to difficult visibility and evaporation pure urine samples could only be occasionally found and are not a promising frequent sample source in European bison.

Hair as a well proven and potential non-invasive source for genotyping was collected in the further comprehensive sampling. During moulting wisent intensively rubbing against tree trunks and stumps sometimes called 'bison combs'[30]. Especially for free-roaming herds those bison combs are potentially sources for non-invasive hair samples but also characterised by become polished, therefore heavily used and prone for genetic contamination. Such bison combs were not sampled within this study. Only in occasional cases hair samples were collected non-invasively utilising comparable objects like brushes and stable walls (Supplementary Figure S10).

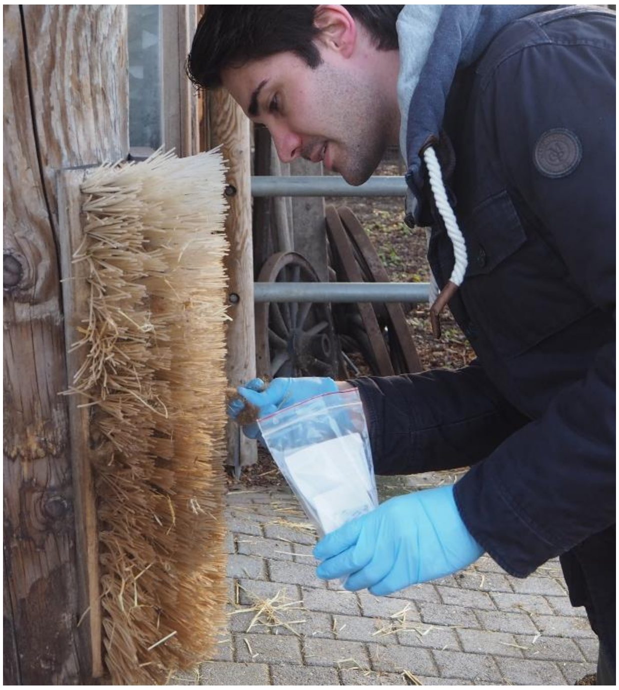

**Supplementary Figure S10: Non-invasive sampling of wisent hair from a rubbing brush into a sip-lock bag with silica gel.** Photo: Victoria Reuber

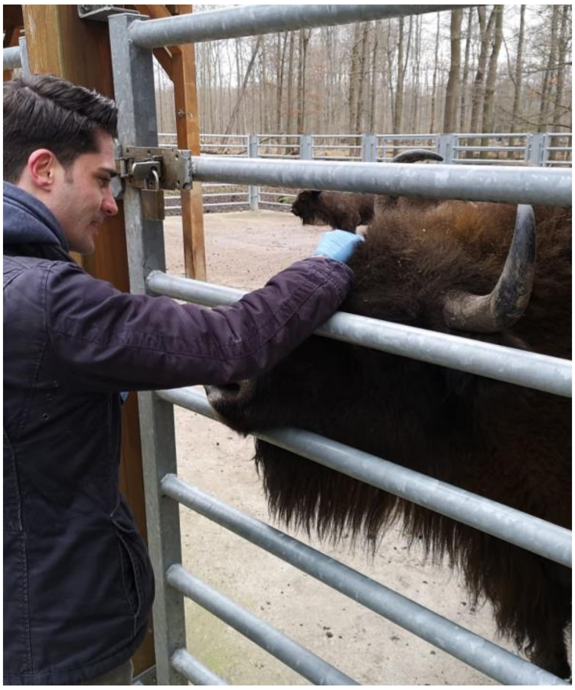

**Supplementary Figure S11: Invasive sampling of hair from the forehead of a female wisent (LL).** This exact method is not possible in the majority of collecting. Most invasive hair samples were taken while anaesthetisation or in corral systems (Supplementary Figure S12). Photo: Felix Rudzinski

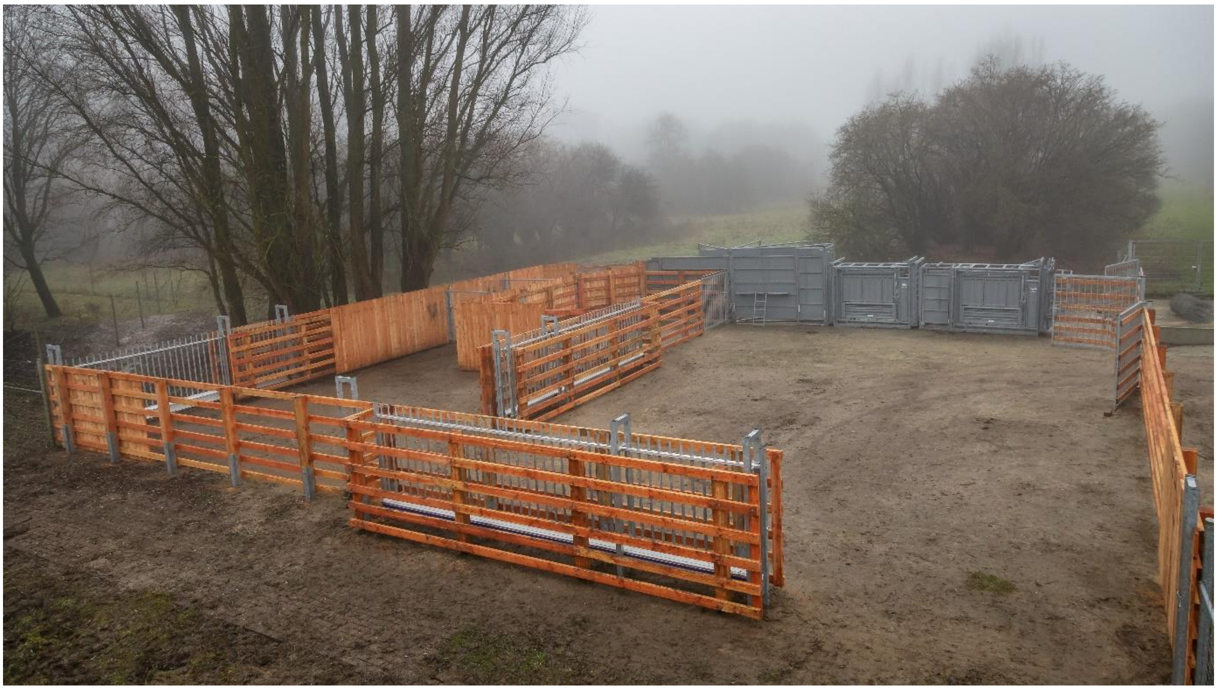

**Supplementary Figure S12: Corral systems like in Lelystad (Natuurpark) are recommendable installations to sample e.g. invasive but innocuous hair samples without anaesthetisation.** Photo: Randy van Domselaar

##### *Conclusion: Dung as source for genetic analyses in European bison*

Faeces is a frequently used environmental sample type that has been proven to be a viable source for DNA in numerous genetic assessments[21,31,39,51,67]. Many bovids (Bovidae GRAY 1821) utilise specific localised defaecation sites or latrines for urination and/or defaecation often resulting in dung piles known as 'dung heaps' or 'dung middens'[68–70]. While this behaviour might hamper genetic assessments due to intraspecific cross-contamination among individuals, it has not yet been observed in European bison. In contrast, Taylor[71] noted that taurine cattle do not even show any sign of field division regarding foraging and defaecation. In this regard, Bovini is the only tribus within the Bovinae which does not contain species which are reported to utilise such latrines[72–81]. Additionally, taurine cattle showed avoidance towards their own faeces to a certain extent[71]. Therefore, pristine Bovini dung pats provide a viable sample type for genetic monitoring with a relatively low risk of intraspecific cross-contamination but might hold complications in other bovids. Furthermore, the wisent, with a daily intake of as much as 30 kg of vegetable biomass with low digestibility defaecates between 5 – 7 kg of dung per day[82] providing an exceptional frequent and therefore pivotal source for non-invasive sampling. Even though, dung probably represents the optimal sample source in non-invasive genetic monitoring in the European bison, it is not completely free from potential contamination risks which should always be considered: licking each other exhibited in grooming behaviour especially between mother and calf or in sexual behaviour between male and female[30,43] are potential intraspecific sources of cross-contaminants in e.g. faecal samples[39].

Even if often proven as a viable DNA source for genetic studies, faeces still represent low-quality samples and need to be evaluated regarding the optimal sampling and sample storage strategy as measured by their PCR amplification success[21]. During quantification of nucleic acids in the samples, different initial concentrations and longevity of both DNA types during storage could lead to, to a certain extent, false conclusions to use amplification values of mtDNA to evaluate nuclear DNA in faeces[83]. Consequently, we used (nuclear) microsatellites to evaluate the reliability of different faecal sampling, DNA extraction and storage methods, considering that the amplification success of microsatellites was identified as good indicator for genotype qualities of SNPs[84] beyond their actual meaningfulness for population genetics in the European bison.

The pilot study provides a faecal sampling methodology evaluation on which (i) DNA extraction kit, (ii) sampled faecal part, (iii) sample storage duration, (iv) sampling and storage type is the most promising for genotyping and applicability in a comprehensive genetic population monitoring of European bison. Additional comments on other (non-)invasive sample types are included. In conclusion, dung samples of European bison were identified as a suitable source for genetic analyses. The generated high-quality SNP genotypes allowed for investigating several population genetic questions in European bison. While collecting faecal swab samples directly in InhibitEX buffer had shown to entail the best success rates with relatively low error rates, the collection of full samples in EtOH clearly has the advantage of providing back-up material for further genetic analysis as well as additional investigations, such as diet studies if needed. Thus, the collection of decent faecal swab samples in InhibitEX buffer and full faecal samples in 96 % EtOH with a contemporary DNA extraction are recommended and were used for further comprehensive sampling in the main study, consistent with others[38]. In general, minor evidence of cross-contamination was found in dung samples genotyped with microsatellites. Beside the fact that Bovini do not use latrines, the pilot study represents the molecular proof of the viability of dung as a DNA source for genetic assessment in this group.

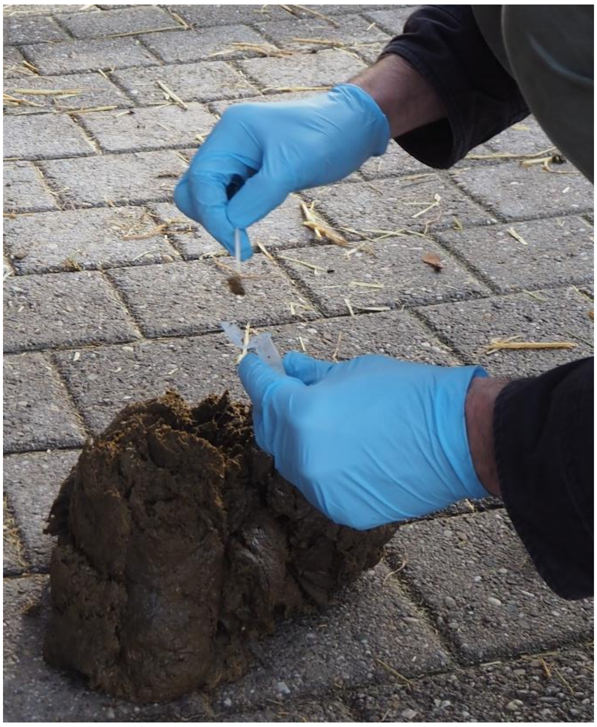

**Supplementary Figure S13: Collecting faecal swab sample into a 2.0 ml reaction tube with lysis buffer.** A decent amount of faecal matrix should be transferred to 100 µl buffer. The wisent pat shown here has a comparable solid consistence. Photo: Victoria Reuber

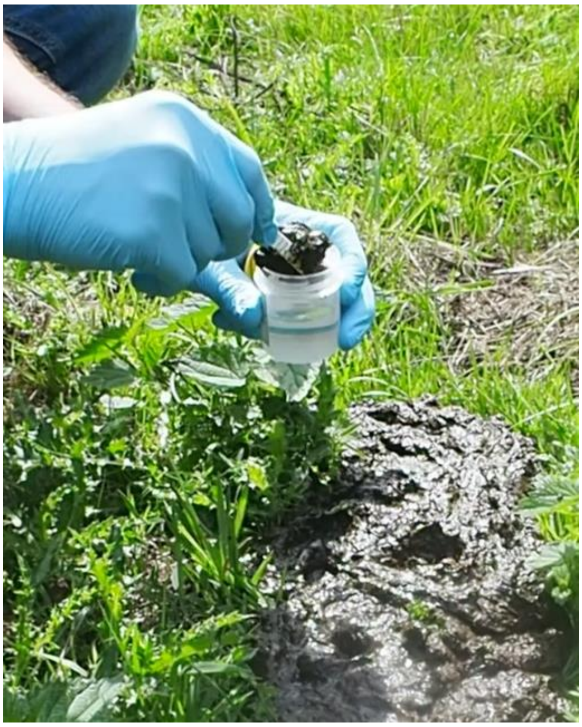

**Supplementary Figure S14: Collecting full faecal sample into a collection cub with 33 ml 96 % EtOH with a one-way forceps.** The wisent pat shown here had a relatively loose consistence, which is more common. Photo: Kaja Heising

### Supplementary Methods

#### Pilot Study: best practice for faecal sampling, preservation and DNA extraction

##### *Sampling and preservation procedures*

For the main study objective, 38 faecal samples from two wisent pats with several sampling types were collected at the 8$^{th}$ August 2018 in the *Wildpark Alte Fasanerie* in Hanau-Klein-Auheim, Hesse, Germany. Therefore, occasional defaecation of every wisent was observed to secure individual assignment of the sampled dung pats. Accordingly, those two dung pats originating from two ('Falka' EBPB#9318 and 'Abia' EBPB#13637) out of four captive individuals were obtained in order to test for an optimal faecal sampling as following: one-way forceps were used to isolate a portion of up to 10 – 15 g faecal matrix for full conservation in 33 ml 96 % EtOH (70 ml cup, SARSTEDT) (in the following *full faecal sample*). For the methodological preservation evaluation three types of swab storing conditions were used: (i) directly in DNA lysis buffer (ASL buffer (QIAamp DNA Stool Mini Kit (Qiagen)) and InhibitEX buffer (QIAampFast DNA Stool Mini Kit (Qiagen)), (ii) in dry bags and (iii) in 33 ml 96 % EtOH. Faecal cotton swab samples were taken separately from the faecal surface and faecal interior. Secondary, dung pats were sampled with dry flocked nylon swabs (4N6FLOQSwabs genetics™ regular size tips in 109 mm long tube with Active Drying System (COPAN flock technologies)). The environmental temperatures ranged from 29 – 32 °C whereas the humidity was measured between 40 – 55 % relative humidity (RH) during sampling. Environmental temperature and RH were assessed with a WindMate$^{tm}$ 300 (Speedtech Instruments).

##### *Microsatellite genotyping and analysis*

The 21 microsatellite primers were allocated in three multiplex mixes (Multiplex A to C) with adjusted concentrations of each marker (Supplementary Table S7). Each primer premix (in total 800 µl) was prepared with 640 µl DNA-free water, 80 µl reveres primer (Rev) and 80 µl forward primer (For). The concentration ratios of the fluorescence-labelled forward primers (ForLab) and the non-labelled forward primers (For) were also adjusted for each marker beforehand (Supplementary Table S7).

The multiplex PCR premix included 5 µl 2× Hot StarTaq Master Mix (QIAgen), 1.4 µl primer premix, 4 µg Bovine serum albumin (BSA, Molecular Biology Grade, B9000S, New England Biolabs® Inc.), 1.4 µl DNA-free water and 3 µl DNA extract.

The microsatellite sequencing target DNA was amplified with a PCR in T1 thermocyclers (Biometra, Analytik Jena) with the following program: Initial denaturation at 95 °C (15 min); followed by 40 cycles denaturation at 94 °C (30 s), annealing at 56 °C (1 min) and extension at 72 °C (1 min); followed by a final extension at 72 °C (10 min); followed by cooling at 10 °C.

PCR products were separated and detected on the ABI 3730 Genetic Analyzer (Thermo Fisher Scientific, Waltham, MA, USA). Allele sizes were determined based on the GeneScan™ 600 LIZ size standard (ThermoFisher Scientific) using GeneMarker® v2.6.3 (Softgenetics®). All automatic scorings were visually checked and if applicable manually corrected. Threshold for calling was set at a minimal florescence of 100 Relative Fluorescent Units (RFUs) for peaks in markers < 200 bp and a florescence of 80 RFUs for peaks in markers > 200 bp if the background noise of fluorescence is moderate around zero. Scoring errors and null alleles were evaluated with *Micro-Checker* v2.2.3[7] with a confidence interval of 95 %.

**Supplementary Table S6: Overview of all utilised microsatellite markers.** All bold printed markers ($n$ = 14) were used for the faecal sampling and storage methodological experiment. All other markers were rejected by disfunction or homozygoty. Marker premix and multiplex protocols from Westekemper *et al.*[85].

| *Locus* | *Primer sequence* ($\frac{forward}{reverse}$) | *Motif* | *Fluorescent label* | *Multiplex* | *References* | *Accession#* |
|---|---|---|---|---|---|---|
| BM4208 | **TCAGTACACTGGCCACCATG** | **(GT)** | **PET** | **A** | [86,87] | **G18509.1** |
| | **CACTGCATGCTTTTCCAAAC** | | | | | |
| *CSSM66* | AATTTAATGCACTGAGGAGCTTGG | (GT) | NED | A | [88–90] | AF232764.1 |
| | ACACAAATCCTTTCTGCCAGCTTGA | | | | | |
| DIK082 | **CCCACTCTGTCTCCAGTTTG** | **(GT)** | **6-FAM** | **A** | [91] | **D83304.1** |
| | **TATCCTGAGAAAAGCTGCTAGA** | | | | | |
| IDVGA59 | **CAGTCCCTCAACCCTCTTTTC** | **$(AC)_{23}$** | **VIC** | **A** | [92] | **X85074.1** |
| | **AACCCAAATATCCATCAATAG** | | | | | |
| *NVHRT21* | GCAGCGGAGAGGAACAAAAG | $(GT)_{16}(GC)_4GT$ | VIC | A | [93] | AF068207.1 |
| | GGGGAGGAGCAGGGAAATC | | | | | |
| *NVHRT48* | CGTGAATCTTAACCAGGTCT | (GT) | NED | A | [93] | AF068214.1 |
| | GGTCAGCTTCATTTAGAAAC | | | | | |
| *RT1* | TGCCTTCTTTCATCCAACAA | (GT) | PET | A | [94] | U90737.1 |
| | CATCTTCCCATCCTCTTTAC | | | | | |
| BM1818 | **AGTGCTTTCAAGGTCCATGC** | **(GT)** | **PET** | **B** | [86] | **G18391.1** |
| | **AGCTGGGAATATAACCAAAGG** | | | | | |
| BM203 | **GGGTGTGACATTTTGTTCCC** | **(GT)** | **6-FAM** | **B** | [86] | **G18500** |
| | **CTGCTCGCCACTAGTCCTTC** | | | | | |
| BMC1009 | **GCACCAGCAGAGAGGACATT** | **$(AC)_{15}$?** | **NED** | **B** | [86] | **?** |
| | **ACCGGCTATTGTCCATCTTG** | | | | | |
| CSSM14 | **AAATGACCTCTCAATGGAAGCTTG** | **(GT)** | **NED** | **B** | [88–90] | **AF232759** |
| | **GAATTCTGGCACTTAATAGGATTCA** | | | | | |
| CSSM19 | **TTGTCAGCAACTTCTTGTATCTTT** | **(GT)** | **VIC** | **B** | [88–90] | **AF232761** |
| | **TGTTTTAAGCCACCCAATTATTTG** | | | | | |
| CSSM22 | **TCTCTCTAATGGAGTTGGTTTTTG** | **(GT)** | **NED** | **B** | [88–90] | **AF232762** |
| | **CTTTCTCTTCAATCAATCCTCATC** | | | | | |

| | | | | | | |
|---|---|---|---|---|---|---|
| *ETH225* | ACATGACAGCCAGCTGCTACT | (GT) | 6-FAM | B | [90,95] | AF232767.1 |
| | GATCACCTTGCCACTATTTCCT | | | | | |
| *CER14* | TCTCTTGCGTCTCCTGCATTGAC | (GT) | 6-FAM | C | [90,96] | L35583.1 |
| | AATGGCACCCACTCCAGTATTCTTC | | | | | |
| CSSM16 | **AGAGCCACTTGTTACACCCCAAAG** | **(GT)** | **NED** | **C** | [90] | **AF232760** |
| | **GATGCAGTCTCCACTTGATTCAAA** | | | | | |
| Haut14 | **CCAGGGAAGATGAAGTGACC** | **(GT)** | **VIC** | **C** | [90] | **AF236378** |
| | **TGACCTTCACTCATGTTATTAA** | | | | | |
| IDVGA55 | **GTGACTGTATTTGTGAACACCTA** | **$(AC)_{12}$** | **NED** | **C** | [92] | **X85071** |
| | **TCTAAAACGGAGGCAGAGATG** | | | | | |
| *INRA35* | TTGTGCTTTATGACACTATCCG | (GT) | PET | C | [90,97] | X68049 |
| | ATCCTTTGCAGCCTCCACATTC | | | | | |
| MM12 | **CAAGACAGGTGTTTCAATCT** | **(GT)** | **6-FAM** | **C** | [90,98] | **Z30343** |
| | **ATCGACTCTGGGGATGATGT** | | | | | |
| KY1/2 | **GCCCAGCAGCCCTTCCAG** | **AmelY/AmelX** | **PET** | **C** | [99] | **FJ434497.1** |
| | **TGGCCAAGCTTCCAGAGGCA** | | | | | |

**Supplementary Table S7: Overview of primer concentrations in the multiplex mixes A to C (µl/reaction) and ratios of the not labelled forward primers (For) and fluorescence-labelled forward primers (ForLab).**

| **Multiplex A** | | | **Multiplex B** | | | **Multiplex C** | | |
|---|---|---|---|---|---|---|---|---|
| Primer | µl/rxn | For:ForLab | Primer | µl/rxn | For:ForLab | Primer | µl/rxn | For:ForLab |
| NVHRT48 | 0.2 | 1:5 | CSSM19 | 0.2 | 1:5 | KY1/2 | 0.2 | 1:10 |
| DIK082 | 0.2 | 1:5 | CSSM22 | 0.2 | 1:10 | MM12 | 0.2 | 1:10 |
| NVHRT21 | 0.2 | 1:5 | ETH225 | 0.2 | 1:10 | Haut14 | 0.3 | 1:4 |
| BM4208 | 0.3 | 1:3 | BMC1009 | 0.2 | 1:5 | CSSM16 | 0.2 | 1:20 |
| CSSM66 | 0.4 | 1:2 | BM203 | 0.2 | 1:5 | CER14 | 0.3 | 1:3 |
| RT1 | 0.4 | 1:3 | BM1818 | 0.2 | 1:5 | INRA35 | 0.2 | 1:5 |
| IDVGA59 | 0.4 | 1:5 | CSSM14 | 0.2 | 1:5 | IDVGA55 | 0.2 | 1:5 |

#### Faecal sampling validation

All faecal samples of both wisent individuals from the wildlife park ('Falka' EBPB#9318 and 'Abia' EBPB#13637) were used for the validation of faecal sampling and preservation methods as well as two DNA extraction kits. Based on these multiple-time sampled and genotyped non-invasive samples, reference genotypes were built to determine the GE rates for each single triplicate genotype. In total, 207 genotypes ('Falka' EBPB#9318: *n* = 105; 'Abia' EBPB#13637: *n* = 102) from 41 samples were generated. Those are comprised of 38 faecal samples and a sole saliva sample collected at the same day in Hanau Klein-Auheim, complemented by two saliva samples from the same two individuals collected in a former sampling to verify the reference individual genotype. The full faecal samples in EtOH were extracted six times (three after one week and three after five weeks). With the exception of the faecal swab samples in lysis buffer all samples were extracted with both the ASL and InhibitEX buffer. All extracts were triplicated for genotyping, while single non-informative triplicates due to missing data or technical error were excluded. The threshold for a valid allele per locus were matching alleles $n > 10$ over all genotypes per individual (≙ 10 % of all genotypes per individual). Three triplicates of a saliva sample (lab#X180120) collected from a feeding trough surface were excluded from analysis due to contamination presumably with DNA of a second individual.

The *amplification success rate*, *genotyping success rate*, *allelic dropout rate* and *false allele rate* as proportional response variables were used to evaluate the applicability of every faecal sampling method, storage and DNA extraction (predictors). The amplification success rate is the number of successful amplified and scored loci per genotype over the total number of loci (*n* = 14). This response variable reflects the applicability measured in successful sequenced PCR products disregarding the final consensus genotype[50]. The genotyping success rate is the number of successful amplified and scored loci per genotype over the total number of loci matching with the individual consensus genotype. Thus, the latter response variable reflects the applicability measured in the result of providing the true genotype, excluding amplification failure (missing data) and genotyping errors (ADOs and FAs)[50]. The ADO rate is the number of observed ADO over the number of total successfully amplified and scored loci per genotype (= amplification success). It was recommended to determine GE rates only as the observed number of GEs over the number of loci where those error can be detected[23,24]. Since all samples in the present study originate from only two individuals and the 14 microsatellite maker were not specifically selected to be heterozygous at every locus, the ADO rate is not calculated from the total number of heterozygous loci based on the consensus genotype ('Falka' EBPB#9318: *n* = 5; 'Abia' EBPB#13637: *n* = 2). ADOs in homozygous loci are a minor problem since they cause no erroneous genotypes like FAs[24]. The FA rate is the number of observed FA over the number of total successfully amplified and scored loci per genotype[24]. The definition of the ADO rate facilitate a comparison with the FA rate in this context and was used before in similar approaches[17,18,100].

Three models per response variable were implemented to test all predictors of the sampling methodology due to the fact that not all samples were allocable to every predictor character (Supplementary Equation S1 - Supplementary Equation S3). Single reactions showing erroneous raw data files (Mix A: *n* = 5; Mix B: *n* = 5; Mix C: *n* = 0) were rejected entirely (total: *n* = 10; 'Falka' EBPB#9318: *n* = 8; 'Abia' EBPB#13637: *n* = 2) for the GLMMs to not bias the explanatory power of the predictor variables on the success or error rates. Those erroneous raw data files were not possible to be displayed into *GeneMarker* v2.6.3 (SoftGenetics) and represent digital errors.

Several sampling and storage methods were tested on the dependence of their success and GE rates to find out a convenient best practice for a genetic study based on non-invasive faecal samples in the European bison. First, the full faecal samples in 96 % EtOH and faecal uptake by swabs were compared. Several containment types of the swab samples are included: cotton swab sample directly in (i) DNA lysis buffer, (ii) cotton swab samples in dry filter paper arranged in a dry bag, (iii) cotton swab samples in 96 % EtOH and (iv) flocked nylon swabs with integrated drying agent (Supplementary Equation S1; Supplementary Equation S3). The impact of two different DNA extraction kits on all sampling methods were tested with the exception of the flocked nylon swabs (only extracted with the InhibitEX buffer) (Supplementary Equation S1 - Supplementary Equation S3). Furthermore, the possible impact of swabs samples from the faecal surface and faecal interior (Supplementary Equation S3) and storage duration on the full samples in EtOH (Supplementary Equation S2) were also tested. Additional to the amplification triplets for each sample, three extraction triplicates of two faecal full samples (lab#X180110; lab#X180111) from two individuals are included per DNA extraction kit and storage duration.

Two random effect groups were implemented in every GLMM: genotypes generated from amplification triplets of the identical sample (laboratory number (lab#)) and the sample assemblage for each automated extraction run in the QIAcube (QIAcube run, 12 samples per run).

**Supplementary Equation S1: GLMM for testing the influence of the categorical predictor variables (sampling method and DNA extraction kit) on the response variable (amplification/genotyping success per locus ($S_i$) of all microsatellite markers (*n* = 14) per reaction/genotyping errors per locus ($E_i$) of successfully amplified and scored microsatellite markers per locus ($AmpS_i$) per reaction).** The random effect variable of the lab number (lab#) represents the single triplicated samples. The random effect variable of the DNA extraction run (QIAcube run) represents the sample assemblage of the automated DNA extraction (this variable was additionally tested). The distribution of the response variable is assumed to be binomial.

$$glmer(cbind(S_i, 14 - S_i)$$
$$glmer(cbind(E_i, AmpS_i - E_i)$$
$$\sim sampling\_method + DNA\_extraction\_kit + (1|Lab\#)[+(1|run)], family\ =\ binomial)$$

Due to the limited number of samples the impact of the storage duration (one week and five weeks after collection) of faecal samples were tested only on full samples in EtOH. The DNA extraction kit is an additional predictor for the success and GE rate (Supplementary Equation S2).

**Supplementary Equation S2: GLMM for testing the influence of the categorical variables (storage duration and DNA extraction kit) on the response variable (amplification/genotyping success of the sequenced microsatellite markers (*n* = 14) per reaction/genotyping errors per locus ($E_i$) of successfully amplified and scored microsatellite markers per locus ($AmpS_i$) per reaction).** The random effect variable of the lab number (lab#) represents the single triplicated samples. The random effect variable of the DNA extraction run (QIAcube run) represents the sample assemblage of the automated DNA extraction (this variable was additionally tested). The distribution of the response variable is assumed to be binomial.

$$glmer(cbind(S_i, 14 - S_i)$$
$$glmer(cbind(E_i, AmpS_i - E_i)$$
$$\sim storage\_duration + DNA\_extraction\_kit + (1|Lab\#)[+(1|run)], family = binomial)$$

Models with the predictors faecal part, sample method and DNA extraction kit were utilised on the success and GE rates to test the reliance of the sampled part of the dung pat. Since a differentiation between faecal surface and faecal interior in the full faecal samples in EtOH was not possible, the latter sampling method was excluded from this model (Supplementary Equation S3).

**Supplementary Equation S3: GLMM for testing the influence of the categorical variables (faecal part, sampling method and DNA extraction kit) on the response variable (amplification/genotyping success of the sequenced microsatellite markers (*n* = 14) per reaction/genotyping errors per locus ($E_i$) of successfully amplified and scored microsatellite markers per locus ($AmpS_i$) per reaction).** The random effect variable of the lab number (lab#) represents the single triplicated samples. The random effect variable of the DNA extraction run (QIAcube run) represents the sample assemblage of the automated DNA extraction (this variable was additionally tested). The distribution of the response variable is assumed to be binomial.

$$glmer(cbind(S_i, 14 - S_i)$$
$$glmer(cbind(E_i, AmpS_i - E_i)$$
$$\sim faecal\_part + sampling\_method + DNA\_extraction\_kit + (1|Lab\#)[+(1|run)], family = binomial)$$

#### Protocol adjustments in SNP panel development

**Supplementary Table S8: Protocol adjustments for preparing the 10X STA Primers for SNP genotyping.** Font colour code: grey = protocols during testing phase; black = protocols for general genotyping of this 96× SNP panel (recommended for adopting in other laboratoires). Abbreviations: LSP = Locus-specific primer (Rev); STA = Specific target amplification primer. Nomenclature was taken from Fluidigm protocol for comparison. For not adjusted protocols see manufacture's documentation for SNPtypeTM Assays for SNP Genotyping (Advanced Development Protocol 34, Fluidigm corp.).

| Preparing the 10X STA Primers | | | | |
|---|---|---|---|---|
| | testing phase (reduction for performing > 96 assays together) | | general genotyping (original protocol) | |
| **Component** | **Volume (µl)** | **final concentration** | **Volume (µl)** | **final concentration** |
| 100 µM SNPtype Assay STA Primer (for each of 96 assasys) | 1 | 250 nM | 2 | 500 nM |
| 101 µM SNPtype Assay LSP Primer (for each of 96 assasys) | 1 | 250 nM | 2 | 500 nM |
| DNA Suspension Buffer | 8 | | 16 | |
| total (aliquot) | 200 | | 400 | |

**Supplementary Table S9: Protocol adjustments for preparing the 10X STA Primers for SNP genotyping.** Colour code: grey = protocols during testing phase; black = protocols for general genotyping of this 96× SNP panel (recommended for adopting in other laboratoires). Abbreviations: STA = Specific target amplification primer; gDNA = genomic DNA. Nomenclature was taken from Fluidigm protocol for comparison. For not adjusted protocols see manufacture's documentation for SNPtypeTM Assays for SNP Genotyping (Advanced Development Protocol 34, Fluidigm corp.).

| Performing STA [pre-amplification PCR Mix] | | | | | | | | |
|---|---|---|---|---|---|---|---|---|
| | original protocol (not utilised) | | invasive reference samples (reduced volumes for resource-efficiency) | | non-invasive samples (testing phase) | | non-invasive samples (general genotyping) | |
| **Component** | **Volume (µl)** | **STA Pre-Mix for 96.96 with Overage (µl)** | **Volume (µl)** | **STA Pre-Mix for 96.96 with Overage (µl)** | **Volume (µl)** | **STA Pre-Mix for 96.96 with Overage (µl)** | **Volume (µl)** | **STA Pre-Mix for 96.96 with Overage (µl)** |
| Qiagen 2X Multiplex PCR Master Mix (Quiagen, PN206143) | 2.5 | 300 | 2.5 | 300 | 5 | 600 | 5 | 600 |
| 10X SNPtype Assay STA Primers (500 nM each) | 0.5 | 60 | 0.5 | 60 | 2 | 240 | 1 | 120 |
| PCR-certified water | 0.75 | 90 | 0 | 0 | 0 | 0 | 1 | 120 |
| gDNA | 1.25 | | 2 | | 3 | | 3 | |
| total | 5 | 450 | 5 | 360 | 10 | 840 | 10 | 840 |

**Supplementary Table S10: Protocol adjustments for preparing the 10X STA Primers for SNP genotyping.** Colour code: grey = protocols during testing phase; black = protocols for general genotyping of this 96× SNP panel (recommended for adopting in other laboratoires). Abbreviations: ASP = Allele-Specific Primers; LSP = Locus-specific primer (Rev). Nomenclature was taken from Fluidigm protocol for comparison. For not adjusted protocols see manufacture's documentation for SNPtypeTM Assays for SNP Genotyping (Advanced Development Protocol 34, Fluidigm corp.).

| Prepering SNPtype Assay Mixes | | | | |
|---|---|---|---|---|
| | original protocol | reduced volume to 30 µl | reduced volume to 20 µl | |
| **Component** | **Volume (µl)** | **Volume (µl)** | **Volume (µl)** | **Final Concentration** |
| SNPtype Assay ASP1/ASP2 (100 µM) | 3 | 2.25 | 1.5 | 7.5 µM |
| SNPtype Assay LSP (100 µM) | 8 | 6 | 4 | 20 µM |
| DNA Suspension Buffer | 29 | 21.75 | 14.5 | |
| total | 40 | 30 | 20 | |

### Supplementary References

1. Tokarska, M., Pertoldi, C., Kowalczyk, R. & Perzanowski, K. Genetic status of the European bison Bison bonasus after extinction in the wild and subsequent recovery. *Mammal Review* **41**, 151–162 (2011).
2. Flisikowski, K., Krasińska, A. M. & Siadkowska, T. S. Genetic polymorphism in selected gene loci in a sample of Białowieża population of European bison (Bison bonasus). *Animal Science Papers and Reports* **25**, 221–230 (2007).
3. Lucy, M. C. *et al.* Variants of somatotropin in cattle: gene frequencies in major dairy breeds and associated milk production. *Domestic Animal Endocrinology* **10**, 325–333 (1993).
4. Mikhailova, M. & Voitsukhovskaya, Y. Comparison of the Lowland and Lowland-Caucasian lines of the European bison (Bison bonasus). in *Abstract eBook: The 3rd International Symposium on EuroAsian Biodiversity* (eds. Semiz, G. & Akyildiz, G. K.) 24 (2017).
5. Chakraborty, R., Andrade, M. D. E., Daiger, S. P. & Budowle, B. Apparent heterozygote deficiencies observed in DNA typing data and their implications in forensic applications. *Ann Human Genet* **56**, 45–57 (1992).
6. Brookfield, J. F. Y. A simple new method for estimating null allele frequency from heterozygote deficiency. *Mol Ecol* **5**, 453–455 (1996).
7. van Oosterhout, C., Hutchinson, W. F., Wills, D. P. M. & Shipley, P. micro-checker: software for identifying and correcting genotyping errors in microsatellite data. *Mol Ecol Notes* **4**, 535–538 (2004).
8. Oleński, K. *et al.* Genome-wide association study for posthitis in the free-living population of European bison (Bison bonasus). *Biol Direct* **10**, 2 (2015).
9. Waits, L. P., Luikart, G. & Taberlet, P. Estimating the probability of identity among genotypes in natural populations: cautions and guidelines. *Mol Ecol* **10**, 249–56 (2001).
10. Inglis, G. D. & Kalischuk, L. D. Use of PCR for Direct Detection of Campylobacter Species in Bovine Feces. *Applied and Environmental Microbiology* **69**, 3435–3447 (2003).

11. Gioffré, A., Meichtri, L., Zumárraga, M., Rodriguez, R. & Cataldi, A. Evaluation of a QIAamp DNA stool purification kit for Shiga-toxigenic Escherichia coli detection in bovine fecal swabs by PCR. *Revista Argentina de microbiologia* **36**, 1–5 (2004).

12. Inglis, G. D., Kalischuk, L. D. & Busz, H. W. Chronic shedding of Campylobacter species in beef cattle. *J Appl Microbiol* **97**, 410–20 (2004).

13. Mommens, G., van Zeveren, A. & Peelman, L. J. Effectiveness of bovine microsatellites in resolving paternity cases in American bison, Bison bison L. *Anim Genet* **29**, 12–18 (1998).

14. Luenser, K., Fickel, J., Lehnen, A., Speck, S. & Ludwig, A. Low level of genetic variability in European bisons (Bison bonasus) from the Bialowieza National Park in Poland. *Eur J Wildl Res* **51**, 84–87 (2005).

15. Tokarska, M. *et al.* Effectiveness of microsatellite and SNP markers for parentage and identity analysis in species with low genetic diversity: The case of European bison. *Heredity (Edinb)* **103**, 326–32 (2009).

16. Launhardt, K., Epplen, C., Epplen, J. T. & Winkler, P. Amplification of microsatellites adapted from human systems in faecal DNA of wild Hanuman langurs (Presbytis entellus). *Electrophoresis* **19**, 1356–61 (1998).

17. Bayes, M. K., Smith K. L., Alberts, S. C., Altmann, J. & Bruford, M. W. Testing the reliability of microsatellite typing from faecal DNA in the savannah baboon. *Conserv Genet* **1**, 173–176 (2000).

18. Smith, K. L. *et al.* Cross-species amplification, non-invasive genotyping, and non-Mendelian inheritance of human STRPs in Savannah baboons. *Am. J. Primatol.* **51**, 219–227 (2000).

19. Miles, K. A., Holtz, M. N., Lounsberry, Z. T. & Sacks, B. N. A paired comparison of scat-collecting versus scat-swabbing methods for noninvasive recovery of mesocarnivore DNA from an arid environment. *Wildl. Soc. Bull.* **39**, 797–803 (2015).

20. Nsubuga, A. M. *et al.* Factors affecting the amount of genomic DNA extracted from ape faeces and the identification of an improved sample storage method. *Mol Ecol* **13**, 2089–94 (2004).

21. Taberlet, P., Waits, L. P. & Luikart, G. Noninvasive genetic sampling: Look before you leap. *Trends in Ecology & Evolution* **14**, 323–327 (1999).

22. Zuur, A. F., Ieno, E. N., Walker, N. J., Saveliev, A. A. & Smith, G. M. *Mixed effects models and extensions in ecology with R*. (Springer, 2009).

23. Creel, S. *et al.* Population size estimation in Yellowstone wolves with error-prone noninvasive microsatellite genotypes. *Mol Ecol* **12**, 2003–2009 (2003).

24. Broquet, T. & Petit, E. J. Quantifying genotyping errors in noninvasive population genetics. *Mol Ecol* **13**, 3601–8 (2004).

25. Gagneux, P., Boesch, C. & Woodruff, D. S. Microsatellite scoring errors associated with noninvasive genotyping based on nuclear DNA amplified from shed hair. *Mol Ecol* **6**, 861–868 (1997).

26. Maudet, C., Luikart, G., Dubray, D., von Hardenberg, A. & Taberlet, P. Low genotyping error rates in wild ungulate faeces sampled in winter. *Mol Ecol Notes* **4**, 772–775 (2004).

27. Cromsigt, J. P. G. M., Kemp, Y. J. M., Rodriguez, E. & Kivit, H. Rewilding Europe's large grazer community: How functionally diverse are the diets of European bison, cattle, and horses? *Restor Ecol* **90**, 2910 (2017).

28. Zielke, L., Wrage-Mönning, N. & Müller, J. Seasonal preferences in diet selection of semi-free ranging European bison (Bison bonasus). *European Bison Conservation Newsletter* 61–70 (2017).

29. Jędrzejewski, W., Sìdarovìč, V., Siemaszko-Skiendziul, A. & Jędrzejewska, B. *The art of tracking animals*. (Mammal Research Institute Polish Academy of Sciences, 2010).

30. Krasińska, M. & Krasiński, Z. A. *European bison: The nature monograph*. (Springer Berlin Heidelberg, 2013).

31. Gardipee, F. M. Development of fecal DNA sampling methods to assess genetic population structure of Greater Yellowstone bison. (2007).

32. Hájková, P. *et al.* Factors affecting success of PCR amplification of microsatellite loci from otter faeces. *Mol Ecol Notes* **6**, 559–562 (2006).

33. Brinkman, T. J., Schwartz, M. K., Person, D. K., Pilgrim, K. L. & Hundertmark, K. J. Effects of time and rainfall on PCR success using DNA extracted from deer fecal pellets. *Conserv Genet* **11**, 1547–1552 (2010).

34. Wedrowicz, F., Karsa, M., Mosse, J. & Hogan, F. E. Reliable genotyping of the koala (Phascolarctos cinereus) using DNA isolated from a single faecal pellet. *Mol Ecol Resour* **13**, 634–41 (2013).

35. Agetsuma-Yanagihara, Y., Inoue, E. & Agetsuma, N. Effects of time and environmental conditions on the quality of DNA extracted from fecal samples for genotyping of wild deer in a warm temperate broad-leaved forest. *Mamm Res* **62**, 201–207 (2017).

36. Santini, A., Lucchini, V., Fabbri, E. & Randi, E. Ageing and environmental factors affect PCR success in wolf (Canis lupus) excremental DNA samples. *Mol Ecol Notes* **7**, 955–961 (2007).

37. Schultz, A. J., Cristescu, R. H., Littleford-Colquhoun, B. L., Jaccoud, D. & Frère, C. H. Fresh is best: Accurate SNP genotyping from koala scats. *Ecol Evol* **8**, 3139–3151 (2018).

38. Velli, E. *et al.* Ethanol versus swabs: what is a better tool to preserve faecal samples for non-invasive genetic analyses? *Hystrix, the Italian Journal of Mammalogy* **30**, 0–6 (2019).

39. Wasser, S. K., Houston, C. S., Koehler, G. M., Cadd, G. G. & Fain, S. R. Techniques for application of faecal DNA methods to field studies of Ursids. *Mol Ecol* **6**, 1091–1097 (1997).

40. Murphy, M. A. *et al.* An evaluation of long-term preservation methods for brown bear (Ursus arctos) faecal DNA samples. *Conservation Genetics* **3**, 435–440 (2002).

41. Berry, O., Sarre, S. D., Farrington, L. & Aitken, N. Faecal DNA detection of invasive species: the case of feral foxes in Tasmania. *Wildl. Res.* **34**, 1 (2007).

42. Canu, A., Mattioli, L., Santini, A., Apollonio, M. & Scandura, M. 'Video-scats': combining camera trapping and non-invasive genotyping to assess individual identity and hybrid status in gray wolf. *Wildlife Biology* **2017**, 00355 (2017).

43. Krasińska, M., Krasiński, Z. A., Perzanowski, K. & Olech, W. European bison Bison bonasus (Linnaeus, 1758). in *Ecology, Evolution and Behaviour of Wild Cattle* (eds. Melleti, M. & Burton, J.) 115–173 (Cambridge University Press, 2014). doi:10.1017/CBO9781139568098.012.

44. Kowalczyk, R. *et al.* Foraging plasticity allows a large herbivore to persist in a sheltering forest habitat: DNA metabarcoding diet analysis of the European bison. *Forest Ecology and Management* **449**, 117474 (2019).

45. Brown, W. E., Ramsey, D. S. L. & Gaffney, R. Degradation and detection of fox (Vulpes vulpes) scats in Tasmania: evidence from field trials. *Wildl. Res.* **41**, 681 (2014).

46. Ramsey, D. S. L., MacDonald, A. J., Quasim, S., Barclay, C. & Sarre, S. D. An examination of the accuracy of a sequential PCR and sequencing test used to detect the incursion of an invasive species: the case of the red fox in Tasmania. *J Appl Ecol* **52**, 562–570 (2015).

47. Reed, J. Z., Tollit, D. J., Thompson, P. M. & Amos, W. Molecular scatology: the use of molecular genetic analysis to assign species, sex and individual identity to seal faeces. *Mol Ecol* **6**, 225–234 (1997).

48. Farrell, L. E., Roman, J. & Sunquist, M. E. Dietary separation of sympatric carnivores identified by molecular analysis of scats. *Mol Ecol* **9**, 1583–1590 (2000).

49. Goossens, B., Chikhi, L., Utami, S. S., de Ruiter, J. & Brunford, M. W. A multi-samples, multi-extracts approach for microsatellite analysis of faecal samples in an arboreal ape. *Conserv Genet* **1**, 157–162 (2000).

50. Broquet, T., Ménard, N. & Petit, E. J. Noninvasive population genetics: a review of sample source, diet, fragment length and microsatellite motif effects on amplification success and genotyping error rates. *Conserv Genet* **8**, 249–260 (2006).

51. Murphy, M. A., Waits, L. P. & Kendall, K. C. The influence of diet on faecal DNA amplification and sex identification in brown bears (Ursus arctos). *Mol Ecol* **12**, 2261–2265 (2003).

52. *Hoofed mammals*. vol. 2 (Lynx, 2011).

53. Kohn, M., Knauer, F., Stoffella, A., Schröder, W. & Pääbo, S. Conservation genetics of the European brown bear - a study using excremental PCR of nuclear and mitochondrial sequences. *Mol Ecol* **4**, 95–104 (1995).

54. Frantzen, M. A. J., Silk, J. B., Ferguson, J. W. H., Wayne, R. K. & Kohn, M. H. Empirical evaluation of preservation methods for faecal DNA. *Mol Ecol* **7**, 1423–1428 (1998).

55. Soto-Calderón, I. D. *et al.* Effects of storage type and time on DNA amplification success in tropical ungulate faeces. *Mol Ecol Resour* **9**, 471–9 (2009).

56. Valiela, I. An Experimental Study of the Mortality Factors of Larval Musca autumnalis DeGeer. *Ecological Monographs* **39**, 199–225 (1969).

57. Janecek, L. L., Honeycutt, R. L., Adkins, R. M. & Davis, S. K. Mitochondrial gene sequences and the molecular systematics of the artiodactyl subfamily bovinae. *Mol Phylogenet Evol* **6**, 107–19 (1996).

58. Hassanin, A. & Ropiquet, A. Molecular phylogeny of the tribe Bovini (Bovidae, Bovinae) and the taxonomic status of the Kouprey, Bos sauveli Urbain 1937. *Mol Phylogenet Evol* **33**, 896–907 (2004).

59. Verkaar, E. L. C., Nijman, I. J., Beeke, M., Hanekamp, E. & Lenstra, J. A. Maternal and paternal lineages in cross-breeding bovine species. Has wisent a hybrid origin? *Mol Biol Evol* **21**, 1165–70 (2004).

60. Soubrier, J. *et al.* Early cave art and ancient DNA record the origin of European bison. *Nat Commun* **7**, 13158 (2016).

61. Palacio, P. *et al.* Genome data on the extinct Bison schoetensacki establish it as a sister species of the extant European bison (Bison bonasus). *BMC Evol Biol* **17**, 48 (2017).

62. Meyer, H., Zentek, J., Habernoll, H. & Maskell, I. Digestibility and Compatibility of Mixed Diets and Faecal Consistency in Different Breeds of Dog. *J Vet Med Series A* **46**, 155–166 (1999).

63. Hayaishi, S. & Kawamoto, Y. Low genetic diversity and biased distribution of mitochondrial DNA haplotypes in the Japanese macaque (Macaca fuscata yakui) on Yakushima Island. *Primates* **47**, 158–64 (2006).

64. Beja-Pereira, A., Oliveira, R., Alves, P. C., Schwartz, M. K. & Luikart, G. Advancing ecological understandings through technological transformations in noninvasive genetics. *Mol Ecol Resour* **9**, 1279–301 (2009).

65. Ruiz-González, A., Madeira, M. J., Randi, E., Urra, F. & Gómez-Moliner, B. J. Non-invasive genetic sampling of sympatric marten species (Martes martes and Martes foina): assessing species and individual identification success rates on faecal DNA genotyping. *Eur J Wildl Res* **59**, 371–386 (2013).

66. Valiere, N. & Taberlet, P. Urine collected in the field as a source of DNA for species and individual identification. *Mol Ecol* **9**, 2150–2152 (2003).

67. Waits, L. P. & Paetkau, D. Noninvasive Genetic Sampling Tools for Wildlife Biologists:: A Review of Applications and Recommendations for Accurate Data Collection. *Journal of Wildlife Management* **69**, 1419–1433 (2005).

68. Hassanin, A. *et al.* Pattern and timing of diversification of Cetartiodactyla (Mammalia, Laurasiatheria), as revealed by a comprehensive analysis of mitochondrial genomes. *C R Biol* **335**, 32–50 (2012).

69. Bibi, F. A multi-calibrated mitochondrial phylogeny of extant Bovidae (Artiodactyla, Ruminantia) and the importance of the fossil record to systematics. *BMC Evol Biol* **13**, 166 (2013).

70. Zurano, J. P. *et al.* Cetartiodactyla: Updating a time-calibrated molecular phylogeny. *Mol Phylogenet Evol* **133**, 256–262 (2019).

71. Taylor, E. L. Grazing behaviour and helminthic disease. *The British Journal of Animal Behaviour* **2**, 61–62 (1954).

72. Walther, F. R. Einige Verhaltensbeobachtungen an Thomsongazellen (Gazella thomsoni Günther, 1884) im Ngorongoro-Krater. *Zeitschrift für Tierpsychologie* **21**, 871–890 (1964).

73. Jarman, P. J. & Jarman, M. V. Impala Behaviour and its Relevance to Management. in *The Behaviour of Ungulates and its relation to management: Ungulate Behaviour Papers* (eds. Geist, V. & Walther, F. R.) vol. 2 871–881 (1974).

74. Hendrichs, H. Changes in a Population of Dikdik, Madoqua (Rhynchotragus) kirki (Günther 1880). *Zeitschrift für Tierpsychologie* **38**, 55–69 (1975).

75. Essghaier, M. F. A. & Johnson, D. R. Distribution and use of dung heaps by Dorcas gazelle in western Libya. *Mammalia* **45**, (1981).

76. Schütze, H. *Field guide to the mammals of the Kruger National Park*. (Struik, 2002).

77. Wronski, T., Apio, A. & Plath, M. The communicatory significance of localised defecation sites in bushbuck (Tragelaphus scriptus). *Behav Ecol Sociobiol* **60**, 368–378 (2006).

78. Leslie, D. M. Boselaphus Tragocamelus (Artiodactyla: Bovidae). *Mammalian Species* **813**, 1–16 (2008).

79. Sharma, K., Rahmani, A. R. & Chundawat, R. S. Natural History Observations of the four-horned antelope Tetracerus quadricornis. *Journal of the Bombay Natural History Society* **106**, 72–82 (2009).

80. Lunt, N. The role of small antelope in ecosystem functioning in the Matobo Hills, Zimbabwe. (2011).

81. Wronski, T., Apio, A., Plath, M. & Ziege, M. Sex difference in the communicatory significance of localized defecation sites in Arabian gazelles (Gazella arabica). *J Ethol* **31**, 129–140 (2013).

82. Olech, W. & Perzanowski, K. Best practice manual for protection of European Bison. (2015).

83. Morin, P. A., Chambers, K. E., Boesch, C. & Vigilant, L. Quantitative polymerase chain reaction analysis of DNA from noninvasive samples for accurate microsatellite genotyping of wild chimpanzees (Pan troglodytes verus). *Mol Ecol* **10**, 1835–1844 (2001).

84. von Thaden, A. *et al.* Assessing SNP genotyping of noninvasively collected wildlife samples using microfluidic arrays. *Sci Rep* **7**, 10768 (2017).

85. Westekemper, K., Signer, J., Cocchiararo, B., Nowak, C. & Balkenhol, N. Understanding effective isolation of intensively managed red deer populations across Germany.

86. Bishop, M. D. *et al.* A genetic linkage map for cattle. *Genetics* **136**, 619–39 (1994).

87. Talbot, J., Haigh, J. & Plante, Y. A parentage evaluation test in North American Elk (Wapiti) using microsatellites of ovine and bovine origin. *Anim Genet* **27**, 117–119 (1996).

88. Moore, S. S., Barendse, W., Berger, K. T., Armitage, S. M. & Hetzel, D. J. S. Bovine and ovine DNA microsatellites from the EMBL and GENBANK databases. *Anim Genet* **23**, 463–467 (1992).

89. Moore, S. S. *et al.* Characterization of 65 bovine microsatellites. *Mammalian Genome* **5**, 84–90 (1994).

90. Kuehn, R., Schroeder, W., Pirchner, F. & Rottmann, O. Genetic diversity, gene flow and drift in Bavarian red deer populations (Cervus elaphus). *Conserv Genet* **4**, 157–166 (2003).

91. Hirano, T. *et al.* Characterization of 42 highly polymorphic bovine microsatellite markers. *Anim Genet* **27**, 365–368 (1996).

92. Mezzelani, A. *et al.* Chromosomal localization and molecular characterization of 53 cosmid-derived bovine microsatellites. *Mammalian Genome* **6**, 629–635 (1995).

93. Røed, K. H. & Midthjell, L. Microsatellites in reindeer, Rangifer tarandus, and their use in other cervids. *Mol Ecol* **7**, 1773–6 (1998).

94. Wilson, G. A., Strobeck, C., Wu, L. & Coffin, J. W. Characterization of microsatellite loci in caribou Rangifer tarandus , and their use in other artiodactyls. *Mol Ecol* **6**, 697–699 (1997).

95. Steffen, P. *et al.* Isolation and mapping of polymorphic microsatellites in cattle. *Anim Genet* **24**, 121–124 (1993).

96. DeWoody, J. A., Honeycutt, R. L. & Skow, L. C. Microsatellite markers in white-tailed deer. *J Hered* **86**, 317–9 (1995).

97. Vaiman, D. *et al.* A set of 99 cattle microsatellites: characterization, synteny mapping, and polymorphism. *Mammalian Genome* **5**, 288–297 (1994).

98. Mommens, G., Coppieterst, W., Weghe, A., Zeveren, A. & Bouquet, Y. Dinucleotide repeat polymorphism at the bovine MM12E6 and MM8D3 loci. *Anim Genet* **25**, 368–368 (1994).

99. Brinkman, T. J. & Hundertmark, K. J. Sex identification of northern ungulates using low quality and quantity DNA. *Conserv Genet* **10**, 1189–1193 (2009).

100. Dallas, J. F. *et al.* Similar estimates of population genetic composition and sex ratio derived from carcasses and faeces of Eurasian otter Lutra lutra. *Mol Ecol* **12**, 275–282 (2003).
